# Pan-cancer analysis of pyrimidine metabolism reveals signaling pathways connections with chemoresistance role

**DOI:** 10.1101/2023.12.06.570388

**Authors:** Vignesh Ramesh, Mert Demirdizen, Luisa Pinna, Thomas Koed Doktor, Mohammad Aarif Siddiqui, Paolo Ceppi

**Affiliations:** Department of Biochemistry and Molecular Biology, University of Southern Denmark, Campusvej 55, 5230 Odense, Denmark; Interdisciplinary Centre for Clinical Research, University Hospital Erlangen, FAU-Erlangen-Nuremberg, Erlangen, Germany; Department of Molecular Biology and Genetics, Bilkent University, 06800, Ankara, Turkey

**Author notes:** These authors contributed equally to the work (Vignesh Ramesh, Mert Demirdizen). Correspondence:* *(Paolo Ceppi).

**Keywords:** Pyrimidine metabolism, signaling pathways, metabolic process, brequinar, pan-cancer

## Abstract

Deregulated nucleotide metabolism, and in particular increased pyrimidine metabolism (PyMet), has been shown to contribute to various pathological features of cancer including chemoresistance and epithelial-to-mesenchymal transition. However, cancer often encompasses complex signaling and metabolic pathway cascades for its progression, and understanding of these molecular regulatory processes in pyrimidine metabolism is quite limited. Therefore, a comprehensive pan-cancer analysis in around 10,000 gene expression profiles of 32 cancer types was employed using a pathway-based approach utilizing gene-sets representing various signaling and metabolic pathways. The analysis identified several top connections with PyMet including TERT, MTOR, DAX1, HOXA1, TP53 and TNC implying an inter-dependency of regulations which in turn was linked to the chemoresistance mechanisms. PyMet-signaling interactions were validated with *in vitro* derived gene-sets from endogenous thymidylate synthase (*TYMS)*-promoter activity reporter, from *TYMS* knockdown and from brequinar treatment, and further at single cell transcriptome level. Strikingly, brequinar treatment profile showed a strong inverse association pattern with doxorubicin chemoresistance in multiple cancer types. The study highlights the PyMet-pathway interactions and its role in chemoresistance, thereby providing an effective tool for improving PyMet targeting strategy in cancer. The analysis as an accessible resource is available at: www.pype.compbio.sdu.dk

**Highlights:** Pan-cancer analysis showed pyrimidine metabolism connections with signaling pathways Top pathway interactors of pyrimidine metabolism were TERT, HOXA1, TP53 and TNC In vitro derived pyrimidine gene-sets recapitulate cancer patients’ pathway analysis Pyrimidine associated pathways confer chemoresistance in multiple cancer types Pyrimidine metabolic inhibitor brequinar reversed doxorubicin chemoresistance feature

## Introduction

Cancer, a highly heterogenous disease, often characterized with genetic and epigenetic aberrations lead to the accumulation of mutations, deregulated signaling pathways and aberrant metabolic processes in the cell, which in turn facilitates uncontrolled proliferative ability, immune evasion, metastasis and chemoresistance (Hanahan & Weinberg, 2011). Breast, lung and colorectal cancers are the most commonly diagnosed cancers, with lung cancer being the leading cause of cancer-related death (Sung *et al*, 2021). Various curative options have been developed in recent years with survival improvement, although found effective when the disease is diagnosed at early stages (Crosby *et al*, 2022). Improvements in treatment options such as CAR-T cell therapy for hematologic cancers (Dagar *et al*, 2023), immunotherapy for NSCLC and melanoma (André *et al*, 2020; Wang *et al*, 2022), molecular targeted therapy for EGFR mutants with osimertinib in lung cancer (Ramalingam *et al*, 2020), and for HER2-positive breast cancer with tucatinib (Lee, 2020) have shown remarkable success in cancer combating. However, the overall survival rate and response rate remains low for many cancer types. For instance, 5-year overall survival rate of NSCLC is 25% (Caini *et al*, 2022) and for advanced endometrial cancer is 17% (Marth *et al*, 2022). Furthermore, the growing concern about the emergence of drug resistance in cancer patients underscores the challenges associated with the promising chemotherapeutic regimen, leading to disease recurrence and contributing to as much as 90% of cancer-related mortality (Bukowski *et al*, 2020; Ramos *et al*, 2021). All these implying the need for better characterization of cellular and molecular level processes of cancer to exploit therapeutic interventions.

Among the hallmarks of cancer, the foremost feature of cancer cell is to induce uncontrolled proliferation, which in turn is heavily relied on the metabolism of purines and pyrimidines (Mullen & Singh, 2023). Several studies showed that higher expression levels of rate-limiting enzymes of pyrimidine metabolism were observed in more than 80% of cancer types including in lung and liver cancer (Wang *et al*, 2020), breast cancer (Luo *et al*, 2022), gastric cancer (Wu *et al*, 2021) and in acute myeloid leukemia (Wang *et al*, 2021) implicating a strong involvement of pyrimidine metabolism in cancer. Of note, high urea cycle dysregulation (UCD) in cancer contributes to poor prognosis, and showed a significant higher pyrimidine-to-purine ratio in multiple tumors (Lee *et al*, 2018) suggesting the importance of pyrimidine metabolism in tumorigenesis. Besides cellular proliferation, pyrimidine metabolism has also been shown to have prominent roles in several other tumorigenic features such as thymidylate synthase (TS) induced epithelial-to-mesenchymal transition (EMT) mediated metastasis and chemoresistance in breast and lung cancer (Siddiqui *et al*, 2017; Siddiqui *et al*, 2019; Siddiqui *et al*, 2021), and dihydroorotate dehydrogenase (DHODH) involvement in liver metastasis in colorectal cancer (Yamaguchi *et al*, 2019). Indeed, during doxorubicin or cisplatin resistance, an increased *de novo* pyrimidine metabolic flux of dCTP and dTTP pools facilitates DNA repair under genotoxic stress have been observed in triple negative breast cancer (Brown *et al*, 2017). However, tumor development is highly governed by the deregulation in multiple transcriptional programs via signaling pathways and metabolic processes, and oncogenes are tightly connected with the signaling process in mediating the utilization of pyrimidine metabolism both during transcriptional gene activities and at post-translational levels on the metabolic enzymes (Wang *et al*, 2021). Therefore, a comprehensive understanding of the regulatory network behind pyrimidine metabolism is required to provide a better opportunity for therapeutic exploitation of cancer (Sanchez-Vega *et al*, 2018), and to combat drug resistance in cancer.

Advent of next generation sequencing platforms offered massive molecular insights of cancers, by utilizing the transcriptomic profiles of cancer patients, from unraveling the functional level biological processes behind tumorigenesis to clinical outcome (Bass *et al*, 2014; Demircioğlu *et al*, 2019; Frost *et al*, 2020; Thennavan *et al*, 2021; Fäldt Beding *et al*, 2022). Therefore, in the present study, an unprecedented intricate role of pyrimidine metabolism in cancer has been addressed by utilizing the transcriptomic profiles of 32 cancer types from The Cancer Genome Atlas (TCGA), and explored the signaling pathways and metabolic processes that could be involved in the regulation of pyrimidine metabolism (PyMet).

## Results

### A comprehensive pan-cancer analysis has identified pyrimidine metabolic-pathway connections

Initially, the dependency of pyrimidine metabolism genes in cancer was analyzed by exploring a genome-wide CRISPR/Cas9 knockout screening for fitness genes in 324 cancer cell lines from 30 cancer types (Behan *et al*, 2019). Pyrimidine metabolic gene-set was obtained from KEGG database with the removal of polymerase containing genes to have more metabolism-focused analysis (**Supplementary Table S1**). Pan-cancer fitness analysis of PyMet genes showed that around 20% of the genes were cancer-dependent genes based on the priority scores (**Figure 1A**), and among them the majority were rate-limiting enzymes with thymidylate synthase (*TYMS)* as the top ranked gene (30/2852). This implies a strong role for pyrimidine metabolism in cancer development and progression across cancer types. To comprehensively understand the molecular aspects of pyrimidine metabolic genes role in cancer, identifying signaling pathways and metabolic processes associated with pyrimidine metabolism is essential. Therefore, a pathway-based activity pattern analysis was employed in assessing the signaling pathways or metabolic processes using gene-sets in the tumor mRNA expression profiles of The Cancer Genome Atlas (TCGA). First, a total of 10,071 normalized gene expression profiles of TCGA representing 32 cancer types of 12 different cancer origins were collected from cBioportal search engine (**Figure 1B** and **Supplementary Table S2**). Second, multiple gene-sets (n = 585) representing various signaling pathways and metabolic processes were collected from MSigDB (**Supplementary Tables S3 and S4**). The activation scores of the gene-sets for PyMet, signaling pathways and metabolic processes were then assessed in each tumor sample using *z*-score based method. Third, the activation scores of the signaling and metabolic pathways were associated with the pyrimidine metabolic process activity scores in each cancer type using Pearson’s correlation with clinically accepted correlation range (r < -0.3 and r > 0.3), and further analyzed significantly associated pathways of PyMet across 32 cancer types by meta-correlation (**Supplementary Tables S5 and S6**). From the analysis, we identified top strong positively (meta-r = 0.41 to 0.57; meta-*P*-value < 0.0001) and negatively (meta-r = - 0.39 to -0.52; meta-*P*-value < 0.0001) associated pathways of PyMet in more than 20 cancer types (∼65% of the cancer types) as threshold (**Figure 1C** and **Supplementary Table S5**). TERT, ESRRA and CAMP pathways were found positively associated with PyMet in around 80% of the cancer types, while TNC target gene-set was found negatively associated in 78% of the cancer types. Pathways associations in lung adenocarcinoma (N = 510), as a representative of the cancer types, clearly showed the trend in the positive and negative pathway activity levels with the increase in the PyMet activation score (**Figure 1D**). Few connections with inflammatory signaling were also observed. While around 65% of the cancer types showed negative association of PyMet with interferon beta (meta-r = -0.46), few positive connections were found with IL15 (meta-r = 0.40) and IL2 (meta-r = 0.37) in around 55 to 60% of the cancer types (**Supplementary Table S5**). As expected, across 32 cancer types, we also observed other signaling pathways or transcription factor-mediated associations with PyMet such as MYC (meta-r = 0.32 to 0.40), CCND1 & CDK4 (meta-r = 0.43), KRAS (meta-r = 0.43), EMT process or cellular transformation signatures (meta-r = 0.25 to 0.38) and a negative association with TP53 targets (meta-r = -0.39). We have included, when available, multiple gene-sets for the same pathways derived from different experimental settings or sources such as with MYC and TERT which concordantly showed a similar positive association patterns with PyMet indicating the sensitivity of the approach (**Supplementary Table S5**). The top positively and negatively associated metabolic processes with PyMet were also validated with 2 additional PyMet gene-sets (KEGG pyrimidine and REACTOME pyrimidine) in the largest cancer type cohorts of breast invasive carcinoma (N = 1082), colon adenocarcinoma (N = 592) and uterine corpus endometrial carcinoma (N = 527) revealing the sensitivity in detecting the regulatory connections using pathway-based approach (**Supplementary Figures 1A-C** and **Supplementary Table S7**).

**Figure 1.**
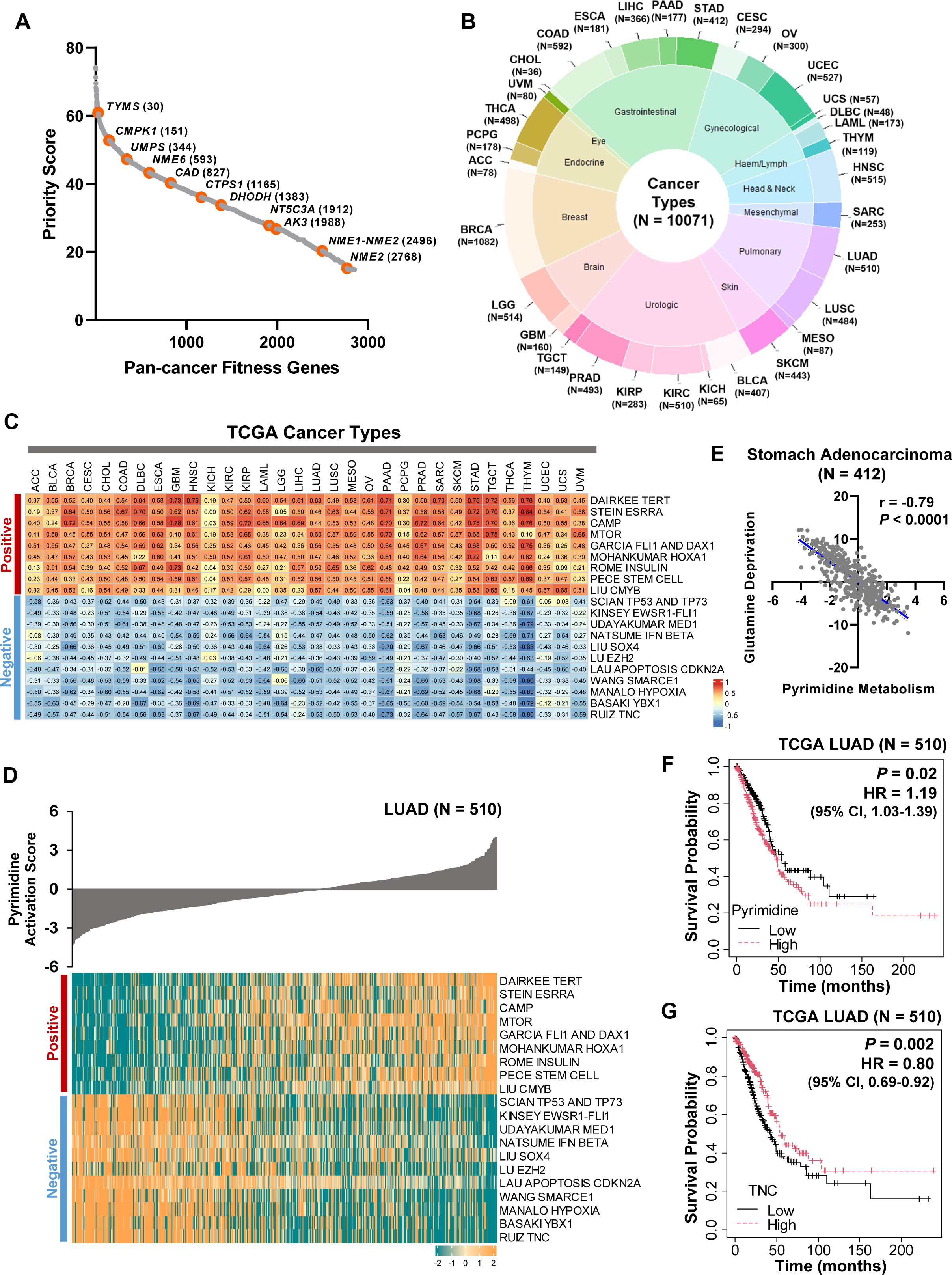
Pathway-based activation analysis of pan-cancer transcriptome profiles has identified novel signaling pathways and metabolic processes of pyrimidine metabolism. A. Ranked plot depicting the priority scores of the pan-cancer fitness genes essential for cancer growth and survival. Highlighted dots represent the pyrimidine metabolic genes essential for cancer with the rank indicated in the brackets. B. Pie-Donut representation of 32 cancer types from 12 different tissue origins with the number of samples used in the study. C. Heatmap visualization of correlation values between top positively and negatively associated signaling pathways with PyMet across 32 cancer types. D. Representative heatmap visualization of the activation pattern of top positively and negatively associated pathways of PyMet in the lung adenocarcinoma patient samples (N = 510) with the increasing trend of PyMet activation levels as a bar chart above heatmap. E. Correlation plot depicting the negative association between the activation scores of glutamine deprivation and PyMet in stomach adenocarcinoma gene expression profile (N = 412). F, G. Overall survival analysis in the TCGA lung adenocarcinoma patient samples categorized as low- and high-levels of pyrimidine metabolic process (F) or TNC signaling (G) based on the median of the activation scores of the gene-sets. HR – Hazard ratio indicated for high-activation scores containing group was calculated using Cox proportional hazards model and *P*-value was calculated using log-rank method.

In parallel, the metabolic processes associated with PyMet were also explored. As anticipated, positive associations were observed between PyMet and nucleotide metabolisms, synthesis of DNA, TCA cycle, OXPHOS and mitotic or cell cycle processes in about 80 to 90% of the cancer types. A pronounced significant negative associations of glutamine deprivation (meta-r = -0.62), leucine deprivation (meta-r = -0.54) and calcium signaling (meta-r = -0.50) with PyMet were observed across cancer types (**Figure 1E**, **Supplementary Figures S1D,E** and **Supplementary Table S6**). Pyrimidine metabolism showed poor prognosis while TNC signaling showed good prognosis in lung adenocarcinoma patient samples (**Figures 1F and 1G**).

Collectively, a comprehensive analysis of pathway activation pattern in different types of cancers has identified several positively and negatively associated signaling pathways and metabolic processes of pyrimidine metabolism.

### Pathways associations with PyMet from cancer patients are recapitulated in cancer cell lines

Having identified the top signaling pathway regulators of PyMet in patient derived tumor samples, we further went onto validate its findings *in vitro*. Gene set enrichment analysis (GSEA) of few of the top positive (HOXA1, ESRRA, and FLI1-DAX1) and negative (TNC and SOX4) interactors of PyMet showed a clear increase and decrease, respectively, in the pyrimidine metabolism in a cancer cell line expression profiles (N = 917) categorized as high and low based on the pathway activation levels (**Figures 2A,B** and **Supplementary Figures S2A-C**) recapitulating the patient derived tumor expression profiles of TCGA. Independently, a previously reported *de novo* pyrimidine synthesis gene, *TYMS* characterized with the non-proliferative role in EMT mediated metastasis (Siddiqui *et al*, 2021), was utilized to validate the findings. *TYMS*-specific gene-set was derived from the FACS sorting of high (TS^HIGH^) and low (TS^LOW^) cell population of Calu-1 cell line (lung squamous carcinoma) stably expressing mCherry reporter gene under the control of *TYMS* promoter followed by RNA-seq profiling (**Figure 2C and Supplementary Table S8**). The derived gene-set (TS^LOW^ and TS^HIGH^) was then used as a proxy of pyrimidine metabolism to examine the functional association between the positive and negative pathways of PyMet, and as a result majority of the pathways showed a similar trend of significant associations as observed in the primary tumors of lung squamous expression profiles (**Figure 2D**). In addition, we used another TS gene-set derived by stable shRNA knockdown of *TYMS* gene in A549 cell line followed by RNA-seq profiling (**Figure 2E** and **Supplementary Table S9**). A similar trend of positive and negative association pattern, although with less significance, was observed with TS knockdown gene-set compared to Calu-1 TS-promoter activity (**Figure 2F**).

**Figure 2.**
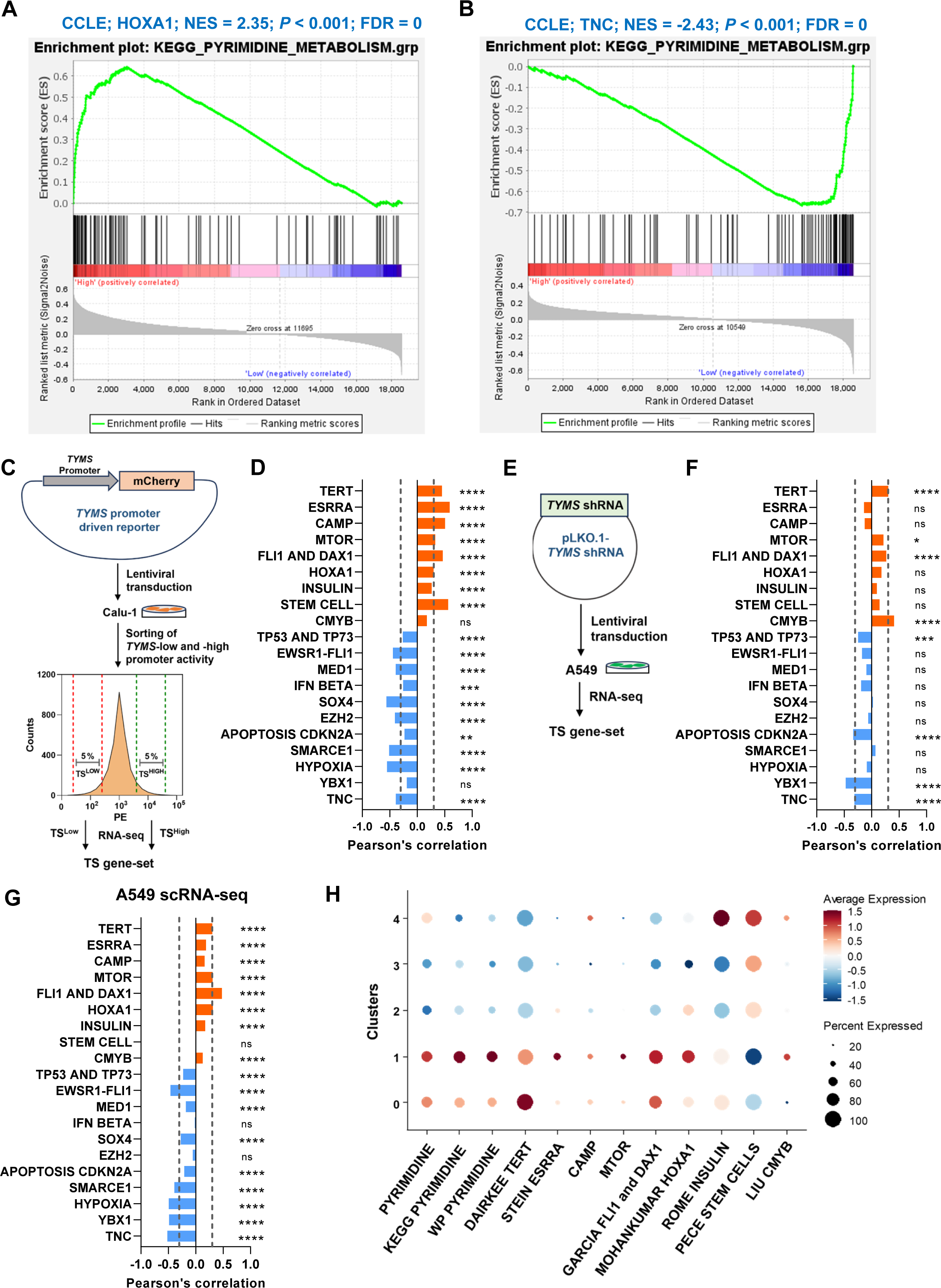
Pathway-based activation analysis in cancer cell lines recapitulate signaling pathway associations with pyrimidine metabolic process. A. Gene-set enrichment analysis of KEGG pyrimidine metabolism gene-set in the gene expression profile of cancer cell line encyclopedia (CCLE) panel (N = 917) categorized as low and high based on the HOXA1 gene-set activation levels showed pyrimidine metabolic process enrichment in high-HOXA1 categorized cell lines. Ranking of genes with signal2noise metric was used for GSEA. B. Gene-set enrichment analysis of KEGG pyrimidine metabolism gene-set in the gene expression profile of cancer cell line encyclopedia (CCLE) panel (N = 917) categorized as low and high based on the TNC gene-set activation levels showed pyrimidine metabolic process enrichment in low-TNC categorized cell lines. Ranking of genes with signal2noise metric was used for GSEA. C. Schematic representation of TS gene-set derived from the RNA-seq expression profiling of Calu-1 cells stably transduced with a lentiviral construct containing mCherry gene under the control of *TYMS* promoter, and sorted the cells based on the low (5%) and high (5%) *TYMS*-driven promoter activity. D. Bar plot of correlation scores calculated between Calu-1 TS gene-set and the positive (orange) or negative (blue) pathways of PyMet. **** - *P*-value < 0.0001; *** - *P*-value < 0.001; ** - *P*-value < 0.01; ns – non-significant. E. Schematic representation of TS gene-set derived from the RNA-seq expression profiling of A549 cells knockdown for *TYMS* gene using stable shRNA lentiviral transduction. F. Bar plot of correlation scores calculated between A549 TS gene-set and the positive (orange) or negative (blue) pathways of PyMet. **** - *P*-value < 0.0001; *** - *P*-value < 0.001; ** - *P*-value < 0.01; * - *P*-value < 0.05; ns – non-significant. G. Bar plot of correlation scores calculated between PyMet and the positive (orange) or negative (blue) pathways of PyMet in parental A549 cells sequenced using single cell RNA-seq. **** - *P*-value < 0.0001; ns – non-significant. H. Dot plot pattern analysis of positively associated signaling pathways of PyMet using AddmoduleScore function in Seurat showed the enrichment of pyrimidine and its associated pathways in cluster 1 of parental A549 cells from scRNA sequencing.

Furthermore, to examine the associations pattern at a single cell expression level, a single cell RNA sequencing of parental A549 cell line (untreated) was explored from our previous study (Ramesh *et al*, 2023), and the analysis revealed a consistent results of positive and negative association of the pathways with the pyrimidine metabolism across 5,203 cells (**Figure 2G**). Remarkably, a cluster-based analysis showed that pyrimidine and its associated positive signaling pathways or metabolic processes were strongly expressed in cluster 1 (**Figure 2H** and **Supplementary Figure S2D**). The negative interacting partners of pyrimidine were only reflected at a minimal level of percent negative expression (**Supplementary Figures S2E** and **S2F**). Overall, co-occurrence pattern of signaling pathway processes and PyMet at the single cell level points out the inherent molecular level regulations and dependency of these identified pathways on pyrimidine metabolism.

### Functional level depletion of pyrimidine metabolism substantiates the signaling pathways interactions

To further substantiate and characterize the involvement of the identified signaling pathways in pyrimidine metabolism, A549 cell line was treated with brequinar (BRQ), an inhibitor of DHODH, a rate limiting enzyme in the initial steps of pyrimidine metabolism (**Figure 3A**). Western blot analysis confirmed the inhibition of TS, a downstream end-product catalyzing enzyme of PyMet, to emphasize the depletion of pyrimidine metabolism in the cells (**Figure 3B**). Differentially expressed genes were then analyzed from the RNA-seq expression profiling of BRQ treated A549 cells with 1 µM for 72 hours which showed maximum depletion of pyrimidine metabolism (**Figure 3C**, **Supplementary Figures S3A** and **Supplementary Table S10**). A significant number of PyMet genes, both *de novo* and salvage, were found in the down-regulated gene list of BRQ including *DHODH*, *TYMS*, *TK1*, *RRM1*, *RRM2* and *UPP1* (**Supplementary Figure S3B**) implying the efficient inhibition of pyrimidine metabolism. Additional independent approaches were carried out to validate the top associated pathways of PyMet in the BRQ treated samples. First, by overlap gene enrichment analysis a significant enrichment of mTORC1 and cell proliferative pathways (MYC and E2F) were observed in the BRQ down-regulated genes, while p53 signaling was enriched in the BRQ up-regulated genes (**Supplementary Figures S3C** and **S3D**). Second, comparison of normalized gene expression of the pathway target genes in the RNA-seq profile of BRQ treated samples with the control revealed a significant up-regulation of TNC and TP53-TP73 target genes with the down-regulation of pyrimidine and FLI1-DAX1 genes (**Figure 3D** and **Supplementary Figures S3E-G**). Third, GSEA analysis was performed in the RNA-seq expression profile of LUAD patient samples categorized as BRQ-high and BRQ-low based on the BRQ gene-set pathway activity scores. The results showed a significant enrichment of HOXA1 down-regulated genes (**Figure 3E**) and TNC up-regulated genes (**Supplementary Figure S3H**) in BRQ-high samples, while HOXA1 up-regulated genes (**Figure 3F**) and pyrimidine (**Supplementary Figure S3I**) enrichment in the BRQ-low samples. Finally, a correlation analysis of the pathway activation scores was also performed in the LUAD patient gene expression profile for PyMet and its associated pathways along with the BRQ gene-set. Interestingly, a striking inverse association pattern was observed between BRQ and pyrimidine’s associated pathways except for EZH2 (**Figure 3G**). In fact, BRQ gene-set showed a strong positive associations with TNC (r = 0.88) and TP53-TP73 (r = 0.85), and negative associations with HOXA1 (r = -0.87) and FLI1-DAX1 (r = -0.86) (**Figure 3H**). Further, BRQ treated A549 and SKMES1 cell lines showed an increase in TNC and decrease in DAX1 protein levels (**Figure 3I** and **Supplementary Figures 3J,K**). Similarly, functionally active phosphorylated form of P53 protein level was found increased with BRQ treatment with a decrease in the KRAS (**Figure 3J**). In addition, knockdown of TS in A549 cell line markedly decreased the phosphorylated form of MTOR and DAX1 while increased the TNC and P16 protein levels compared to the control (**Figure 3K**) concordant with the transcriptome data (**Figure 2F**). Collectively, *in vitro* pyrimidine metabolic inhibitor treatment with BRQ confirms the pathways associations identified from the patients derived transcriptome analysis.

**Figure 3.**
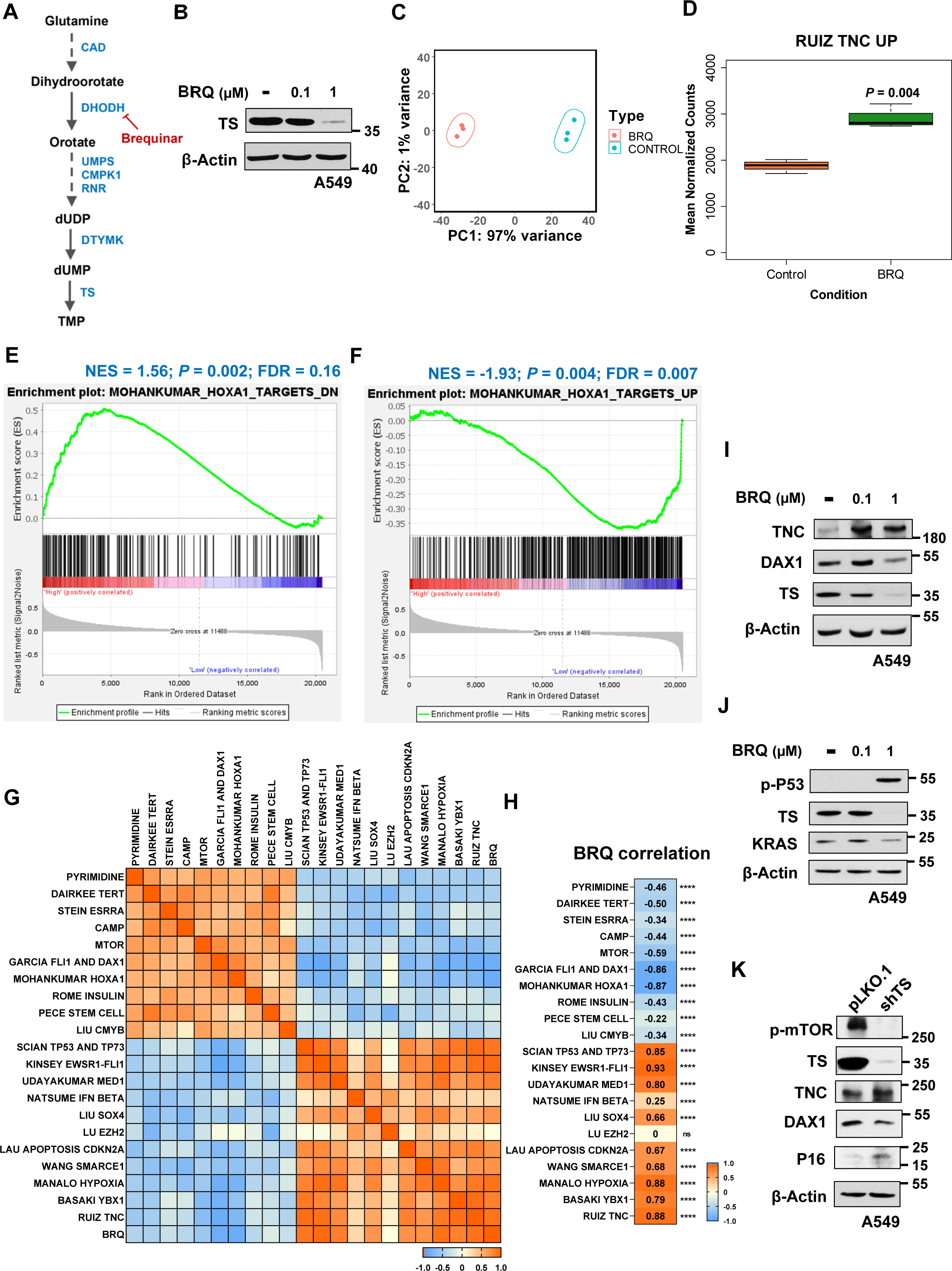
RNA-seq expression profiling of brequinar (BRQ) treated cells validates the pyrimidine associated pathways. A. Schematic representation of pyrimidine metabolic pathway and targeting of DHODH with brequinar (BRQ). B. Western blot analysis of TS protein levels in A549 cells treated with brequinar (BRQ) in the indicated dose dependent concentration for 72 h. β-actin was used as an internal control. C. Principal component analysis of RNA-seq expression profile of A549 cells treated with 1 µM of BRQ for 72 h reveals a high variance in the gene expression profiles between the treated and the control samples (n = 3 per group). D. Box plot visualization of the increased up-regulated gene-set of TNC signaling pathway in the BRQ (1 μM) treated A549 cells compared to the control. The central band inside the box represents the median value of the data (n = 3) obtained using the lower (bottom) and upper (top) quartile values of the box. The maximum and minimum values of the data are displayed with the vertical lines (whiskers) connecting the box. Significance was calculated using un-paired *t*-test between the BRQ treatment and the control conditions. E, F. Gene set enrichment analysis of down-regulated (E) and up-regulated (F) gene-sets of HOXA1 signaling pathway in TCGA lung adenocarcinoma patient gene expression profiles (N = 510) categorized as BRQ-high and BRQ-low based on the median of the BRQ gene-set activation scores. Ranking of genes with signal2noise metric was used for GSEA. G. Correlation matrix plot depicting the associations between BRQ gene-set, PyMet and its associated signaling pathways in lung adenocarcinoma patients (N = 510). H. Correlation values between BRQ gene-set and positively (or negatively) associated pathways of PyMet calculated from the activation scores of the gene-sets in lung adenocarcinoma patients (N = 510). **** - *P*-value < 0.0001; ns – non-significant. I. Western blot analysis of TNC, DAX1 and TS protein levels in A549 cells treated with brequinar (BRQ) in the indicated dose dependent concentration for 72 h. β-actin was used as an internal control. J. Western blot analysis of phosphorylated P53, TS and KRAS protein levels in A549 cells treated with brequinar (BRQ) in the indicated dose dependent concentration for 72 h. β-actin was used as an internal control. K. Western blot analysis of phosphorylated mTOR, TS, TNC, DAX1 and P16 protein levels in A549 cells knockdown for *TYMS* compared to the control (pLKO.1). β-actin was used as an internal control.

### Pyrimidine metabolism and its associated pathways confer chemoresistance in multiple cancer types

As pyrimidine metabolism is involved in various clinical manifestations of tumorigenesis from uncontrolled proliferation, EMT and chemoresistance, we investigated whether the identified PyMet associated signaling pathways also have any involvement with chemoresistance. Therefore, we assessed pathway-based activation scores for multiple gene-sets related to various drugs used in the clinic for multiple cancer types such as fluorouracil, cisplatin, gefitinib, gemcitabine and doxorubicin, the activity of which is well known to be limited by chemoresistance. Among the investigated chemoresistance gene-sets, we identified a very strong positive pattern between doxorubicin chemoresistance and PyMet and its associated pathways. Interestingly, this pattern was found to have inverse association with BRQ in pancreatic and lung adenocarcinoma patient samples (**Figures 4A-D**) indicating that BRQ could potentially reverse the effect of doxorubicin chemoresistance. This pattern of positive interaction between doxorubicin chemoresistance and PyMet’s associated pathways, and the reversal effect of BRQ was found in multiple cancer types including in breast, bladder and low grade glioma (**Supplementary Figure S4**). Notably, the correlation was significantly strong between doxorubicin resistance and PyMet (r > 0.5) or HOXA1 (r > 0.6). Conversely, a strong negative correlation was observed between doxorubicin resistance and BRQ (r = - 0.76 to -0.87) or TNC (r = -0.86 to -0.91) (**Figures 4B** and **4D**). Independently, TNC signaling pathway was found enriched in the gene expression profile of neuroblastoma cell line IMR32 treated with another DHODH inhibitor, teriflunomide (GSE67338) (**Figure 4E**). Minimal levels of positive associations were also observed with other chemoresistance drugs such as cisplatin and gemcitabine. However, analyzing the gene expression pattern of rate-limiting enzymes of PyMet in cisplatin-resistant A549 cell line (**Figure 4F**) or gemcitabine-resistant Calu-3 cell line (**Figure 4G**) revealed an increased expression with the chemoresistance development.

**Figure 4.**
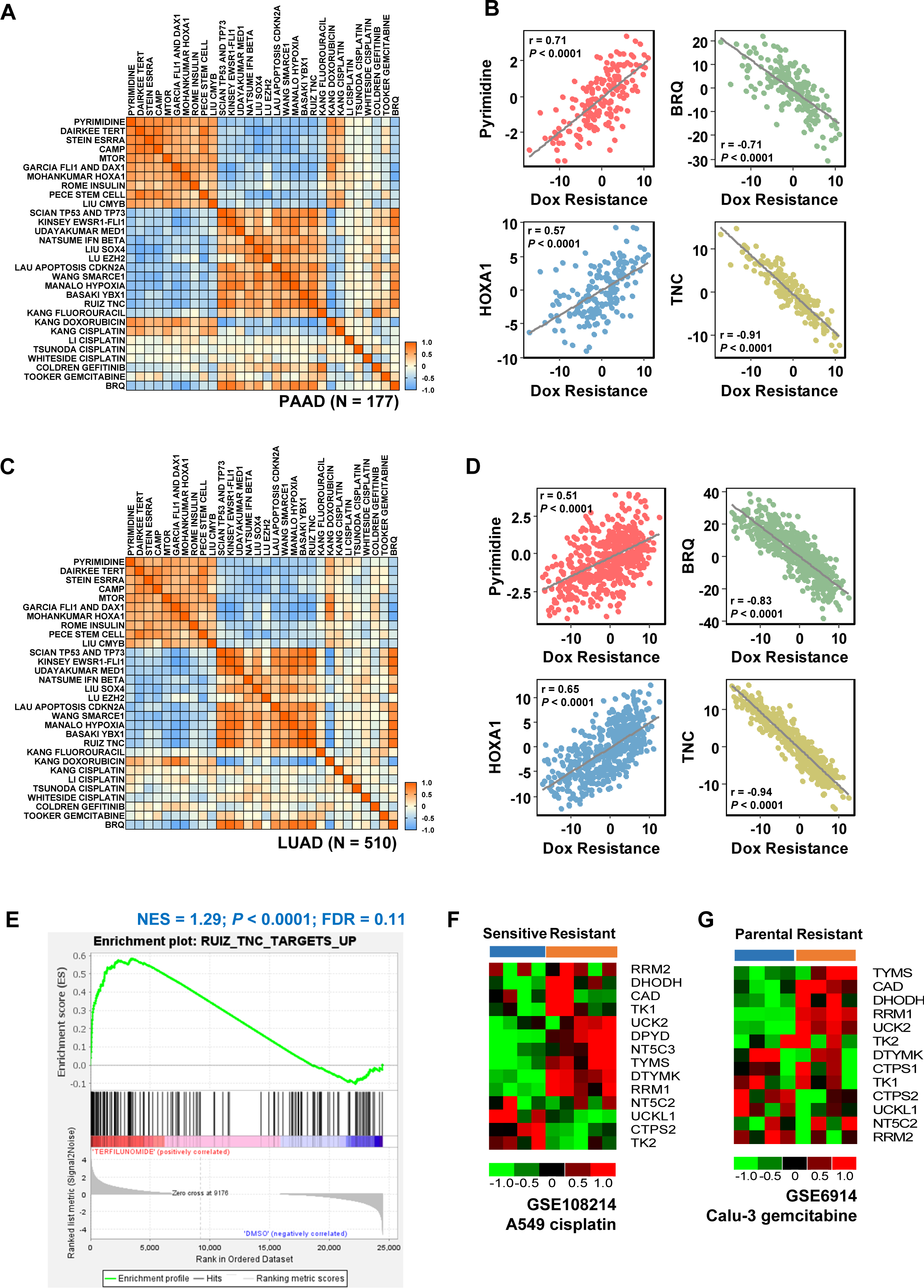
Pyrimidine metabolic process and its associated signaling pathways poses chemoresistance in cancer. A. Correlation matrix plot depicting the associations between drug resistance gene-sets, BRQ, pyrimidine and its associated signaling pathways in pancreatic adenocarcinoma (N = 177). B. Scatter plots of the activation scores between doxorubicin drug resistance and pyrimidine metabolism, its associated signaling pathways (HOXA1 and TNC) and BRQ gene-sets in pancreatic adenocarcinoma (N = 177). Doxorubicin drug resistance showed positive associations with pyrimidine and HOXA1, while negative associations were observed with DHODH inhibitor (BRQ) and TNC signaling pathway. C. Correlation matrix plot depicting the associations between drug resistance gene-sets, BRQ, pyrimidine and its associated signaling pathways in lung adenocarcinoma (N = 510). D. Scatter plots of the activation scores between doxorubicin drug resistance and pyrimidine metabolism, its associated signaling pathways (HOXA1 and TNC) and BRQ gene-sets in lung adenocarcinoma (N = 510). Doxorubicin drug resistance showed positive associations with pyrimidine and HOXA1, while negative associations were observed with DHODH inhibitor (BRQ) and TNC signaling pathway. E. Gene set enrichment analysis of up-regulated gene-set of TNC signaling pathway in the gene expression profile of IMR32 cell line treated with teriflunomide obtained from GEO (GSE67338). Ranking of genes with signal2noise metric was used for GSEA. F. Heatmap visualization of the rate-limiting genes of pyrimidine metabolism in the gene expression profile of A549 cells resistant to cisplatin obtained from GEO (GSE108214). G. Heatmap visualization of the rate-limiting genes of pyrimidine metabolism in the gene expression profile of Calu-3 cells resistant to gemcitabine obtained from GEO (GSE6914).

Taken together, pathway-based analysis revealed a strong involvement of pyrimidine and its associated pathways in drug resistance, and importantly this involvement could be sensitized or inhibited with DHODH inhibitors.

### Pyrimidine Pathway Explorer (PYPE) as a web resource to examine pyrimidine metabolism regulatory process

Having observed the potential of pathway-based approach in identifying the association patterns between PyMet and signaling pathways or metabolic processes, we developed a web portal as a resource for the scientific community (**Figure 5A**). Pyrimidine Pathway Explorer (PYPE) allows exploration of connections between the signaling pathways or metabolic processes at the gene expression based level: First, signaling pathways or metabolic processes correlations with PyMet in individual cancer types and multiple cancer types can be explored using the PYPE tool. Each separate section for signaling pathways (247 gene-sets) and metabolic processes (335 gene-sets) were included in the entry page. By selecting the cancer type and correlation type, the result output generates a list of correlated signaling or metabolic processes gene-sets with PyMet. Further selecting the gene-set in the obtained list, the activation scores across samples, correlation plot between the signaling (or metabolic process) and PyMet, and finally the overlap of gene list between the gene-sets are obtained. Second, a custom correlation analysis section has been provided wherein the users can input their own gene-sets derived from microarray or RNA-seq from different experimental settings. Both options of inputting single or paired direction gene-sets can be imputed by the web tool. The results include the correlation between PyMet gene-set and user-defined gene-set for each individual cancer type. Further, when selected multiple cancer types, a Forest plot of meta-correlation values across selected cancer types can be obtained. In another tab, an overlap analysis report will also be generated between the user-defined gene-sets and PyMet gene-set. We cross-validated our gene-sets derived from TS^LOW^/TS^HIGH^ from Calu-1 and TS-knockdown gene-set from A549 cells in PYPE across cancer types, and identified that majority of the cancer types showed PyMet associations in a significant manner for TS specific gene-sets (**Figures 5B,C**). The developed web server provides a flexible analysis of gene expression based analysis of pyrimidine metabolism in the aspects of signaling pathways and metabolic processes (**Figure 5D**). The findings as an accessible resource is available at: www.pype.compbio.sdu.dk

**Figure 5.**
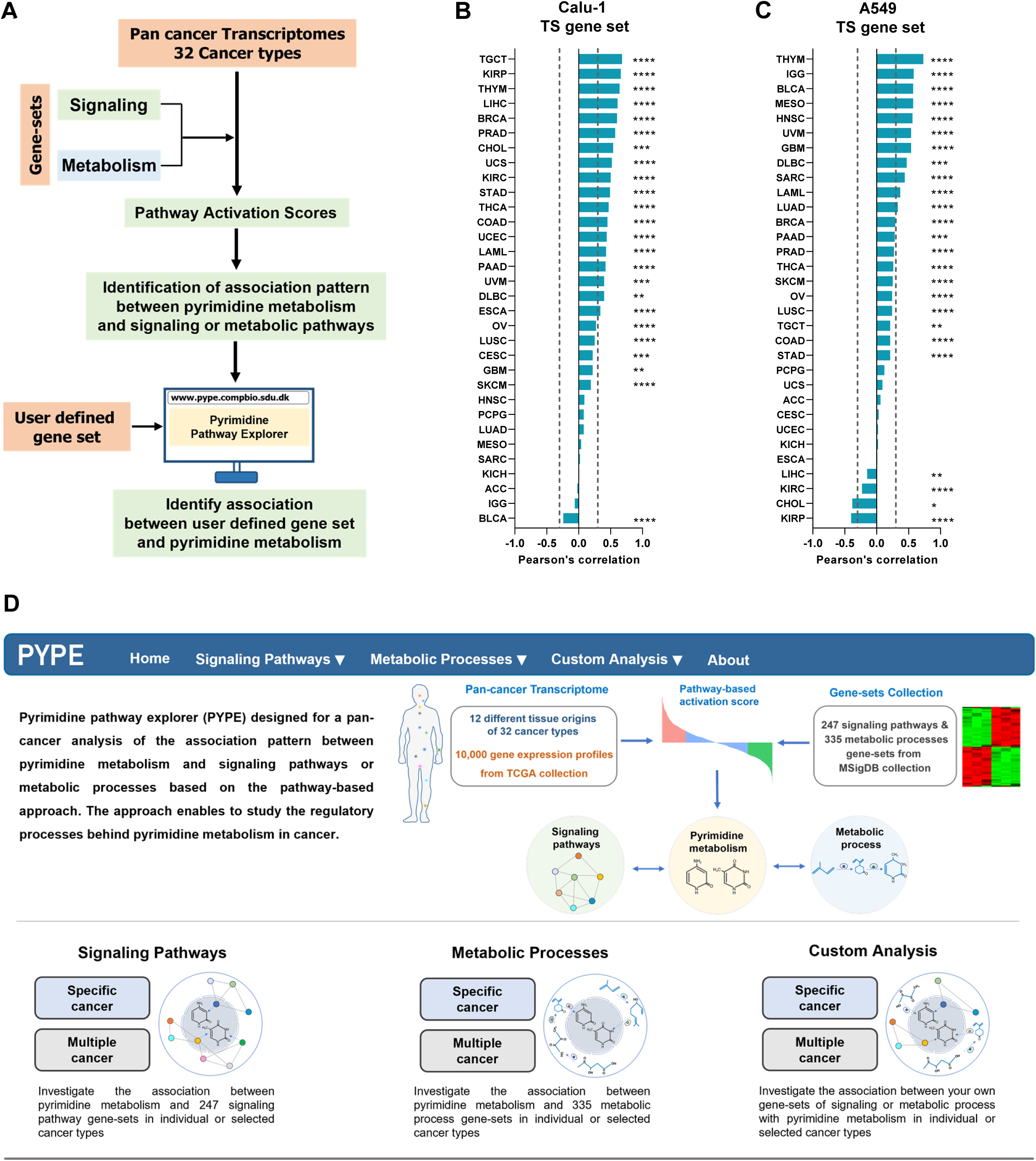
Pyrimidine pathway explorer (PYPE), a web resource for identifying the associations of pyrimidine metabolic process. A. Schematic representation of the pipeline of pyrimidine pathway explorer (PYPE) to analyze the user-defined gene-set and pyrimidine metabolism. B,C. Identification of significant associations of Calu-1 TS gene-set (B) and A549 TS gene-set (C) gene-sets with PyMet in a pan-cancer analysis using PYPE web tool. D. Screenshot of PYPE web resource tool for performing pan-cancer analysis of pyrimidine metabolism with signaling pathways and metabolic processes across 32 cancer types.

## Discussion

Deregulated pyrimidine metabolism contributes to sustained cell proliferation, genome instability, deregulation of cellular bioenergetics, and further conferring advanced tumorigenic features including EMT-mediated metastasis with worse prognosis (Wang *et al*, 2020). In the current pan-cancer study, by utilizing several thousands of gene expression profiles of 32 cancer types, we explored the intricate regulatory network of signaling and metabolic pathways with pyrimidine metabolism using pathway-based approach. Many of the identified associations in this study were also reported previously like with TERT (Mannherz & Agarwal, 2023), KRAS (Koundinya *et al*, 2018) and CCND1 (Qian *et al*, 2020). For instance, thymidine nucleotide metabolic genes (*TYMS*, *DTYMK*, *TK1* and *DCTPP1*) were found to strongly regulate telomere length maintenance identified from a genome-wide CRISPR-Cas9 functional screening (Mannherz & Agarwal, 2023). And PyMet negative partner p53 signaling pathway identified from the current study was also observed previously wherein p53 pathway was activated upon the depletion of pyrimidines relied on DHODH (Khutornenko *et al*, 2010). Similarly, at the metabolic level, dependency of oxidative phosphorylation on pyrimidine metabolism was also in line with the earlier observation wherein regulation of pyrimidine metabolism driven by DHODH is essential in functional oxidative phosphorylation for tumorigenesis (Bajzikova *et al*, 2019). This confirms the sensitivity of the pathway-based approach and its reliability in exploring novel molecular regulatory connections. The involvement of proliferative pathways and EMT associations with pyrimidine metabolism from the pathway analysis indeed support the model of differential contributions of pyrimidine metabolism in cancer (Siddiqui *et al*, 2019). Interestingly, the analysis also identified few novel signaling pathways such as HOXA1 and TNC signaling pathways in the positive and negative regulation of pyrimidine metabolism, respectively, in more than 24 cancer types (>75%) which would be of relevance for future explorations.

Next, a strong recapitulation of the connections between pyrimidine and signaling pathways were also validated using multiple different genomic approaches. First, with CCLE containing 917 cell lines derived from different cancer subtypes. Second, by functionally modulating a principal component gene, *TYMS*, of pyrimidine metabolism both at the gene knockdown level and by capturing endogenous pyrimidine metabolic level from *TYMS-* driven promoter activity. The observation of low significance with *TYMS* knockdown gene-set compared to *TYMS*-driven promoter gene-set in pathways associations could be due to the capturing of the upstream and downstream pathways from undisrupted endogenous level of pyrimidine metabolism with *TYMS*-promoter activity while *TYMS* knockdown could capture only the downstream associated molecular processes. Third, at the single cell RNA sequencing of a parental lung cancer cell line (without any treatment) implying that these regulatory processes are inherent at the cellular level. Previously, chemical synthetic lethal screening has identified DHODH as a target of mutant KRAS, and inhibition of DHODH with brequinar selectively inhibited *in vitro* and *in vivo* KRAS driven cancerous growth including lung (Koundinya *et al*, 2018). Strikingly, this is in line with our study on the significant inverse association between the brequinar gene-set and KRAS mutant activation levels in lung cancer (**Supplementary Figure S3L**). In addition, the derived brequinar gene-set has also shown a precise inverse pattern to the pyrimidine-associated pathways such as HOXA1, MTOR, CCND1-CDK4 and FLI1-DAX1 in the patient derived gene expression profiles indicating a strong regulatory phenomenon by pyrimidine metabolism at the cellular level and high relevance of these targets in synthetic lethality in cancers dependent on pyrimidine metabolism. In fact, a recent screening of inhibitors for FLI1-DAX1 has identified K-234, a DHODH inhibitor, for Ewing’s sarcomas (Watanabe *et al*, 2023). Negative effect of TP53 with pyrimidine was also observed before when DHODH inhibitor treatment increases TP53 levels (Ladds et al., 2018) and substantiated in the current study with an increased phosphorylated-p53 with BRQ treatment. TNC, an extracellular matrix molecule, was found correlated with tumor progression with poor prognosis in multiple cancer types (Liot *et al*, 2020). However, our pathway-based analysis showed a negative association pattern with pyrimidine metabolism across cancer types, and notably the regulation of TNC signaling by DHODH inhibitors would be worth investigating in the future. All these convincingly highlight the importance of the identified signaling pathways and pyrimidine metabolism in targeting several types of cancer.

Cancer drug resistance is an important feature in the recurrence of cancer in patients contributing to poor survival (Ramos *et al*, 2021), and our gene expression analysis of pyrimidine rate limiting enzymes showed a strong pattern of positive regulation of chemoresistance. Strong evidences have shown the importance of DHODH inhibition in sensitizing the cancer cells to overcome the chemoresistance in several cancer types (Madak *et al*, 2019; Zhang *et al*, 2022). Similarly, doxorubicin treatment has been shown to stimulate the increase in the *de novo* pyrimidine synthesis, and pharmacological inhibition of pyrimidine synthesis with leflunomide in combination with doxorubicin has sensitized the TNBC cells (Brown *et al*, 2017). Furthermore, *in vivo* combinatorial treatment of melanoma xenograft with BRQ and doxorubicin had a substantial tumor growth reduction and improved the efficacy of doxorubicin potency (Dorasamy *et al*, 2019). In accordance with these studies, the current study for the first time uncovered the pan-cancer role of signaling pathways and metabolic processes strongly associated with pyrimidine metabolism. And most importantly, it shows that DHODH inhibitor, brequinar, can overcome chemotherapy resistance especially by doxorubicin and cisplatin by targeting multiple signaling pathways dependent on pyrimidine metabolism in several cancer types. Thus, further exploration of the identified signaling regulatory network or metabolic processes of pyrimidine metabolism would help uncover the development of combinatorial therapeutic treatment options for drug resistance cancer types specifically for doxorubicin.

The pathway-based approach in the study includes advantages and few limitations in the utilization of the gene-sets derived from different sources. The current study included multiple numbers of gene-sets for the same pathway or metabolic processes derived from different studies to have concordant results for the pathway associations. For example, all 5 gene-sets of TERT signaling from different studies using different cell line origins showed a consistent positive association with pyrimidine (meta-r = 0.33 to 0.69). Similarly, 5 out of 7 hypoxia different gene-sets obtained by HIF1A overexpression or hypoxia mimic treatments showed a consistent negative association with pyrimidine metabolism (meta-r = -0.47 to -0.62) indicating the increased confidence level of the association patterns. On the other hand, we also observed few pathway gene-sets performed in a context-specific manner. For instance, majority of the KRAS mutant gene-sets derived from different tissue types (lung, breast, prostate) showed a similar positive trend in association with pyrimidine metabolism except for kidney, and KRAS gene-set from lung showed strong positive associations with pyrimidine metabolism in around 20 cancer types (62%) compared to other tissue types derived gene-sets. In another instance, pyrimidine metabolism showed a negative association with YBX1 gene-set from ovarian cancer cell line in 75% cancer types and this outperformed breast cancer cell line derived YBX1 gene-set showing positive association in only 38% cancer types. Nevertheless, when tight meta-correlation thresholds were applied, the later was excluded to reduce false positives. The differential effect in association pattern could have been due to the tissue-specific nature and genetic makeup of the cell line used to derive the gene-sets. With stringent parameters set in the pathway-based analysis and by experimentally validating the pathway associations such context-specific variabilities could be eliminated. Further, in future, the sensitivity of the pathway-based approach could be strengthened by expanding the number of gene-sets for signaling and metabolic pathways in pan-cancer manner to derive a conserved or hallmark gene signatures. A conserved pathway gene signature derived from a pan-cancer perspective could be useful to capture a refined and defined biological processes of the cell which in turn can be used to analyze pathways independent of context-specific nature of the studies.

The current study’s results projected as a web tool resource for the scientific community would be helpful in exploring the association between the pyrimidine metabolism and signaling pathways or metabolic processes across cancer types. The major advantage of pathway-based analysis over the conventional over-representation analysis is that the earlier performs at the gene expression levels in the patient samples which provides more reliable output than the later based on gene content overlap. Besides the in-built features in the PYPE tool, the web resource also allows to analyze the associations between the user defined gene list (for instance experimentally derived) and pyrimidine metabolic pathway across several types of cancers providing a platform to explore the oncogenic processes mediated by pyrimidine metabolism and its associated pathways in multiple cancers. Furthermore, the user defined gene-set analysis could also pave a way in identifying a potential impact of chemoresistance and therapeutic targeting of cancer.

## Conclusion

A comprehensive pan-cancer analysis of pyrimidine metabolism in the context of signaling pathways and metabolic processes showed molecular regulatory connections with clinical features including chemoresistance for oncogenic pyrimidine metabolism. Further, the study highlights the combinatorial therapeutic approach of using DHODH inhibitors to enhance the efficacy of routinely used chemotherapeutic drugs in clinic.

## Methods

### Cell culture

A549 and SKMES1 cell lines (ATCC) were cultured in DMEM high glucose medium (Gibco) and RPMI-1640 (Gibco), respectively, supplemented with 2 mM of L-glutamine (Gibco), 100 U/ml of Penicillin-Streptomycin (Gibco) and 10% Fetal bovine serum (Gibco). Cell lines were maintained at 37° C with 5% CO_2_ level in a humidified incubator. Cell lines were authenticated by STR profiling and routinely tested for mycoplasma contamination.

### Western blot

Around 1 x 10^5^ cells were seeded in a 6-well plate for the drug treatment with Brequinar (Tocris, 6196) in a dose-dependent manner for 72 h. After 72 h treatment, cells were harvested for protein using Pierce RIPA lysis buffer (Thermo Scientific) containing 1X Halt Protease & Phosphatase inhibitor cocktail (Thermo Scientific) and estimated the protein concentrations using Pierce BCA Protein Assay Kit as per the manufacturer’s protocol (ThermoFisher Scientific). Around 10 µg of proteins were allowed to resolve in 8% SDS-PAGE separating gel, and the proteins were then transferred from the separating gel to a PVDF membrane (Thermo Scientific). Membranes with the transferred proteins were blocked with the blocking buffer (5% non-fat dried milk powder or 3% bovine serum albumin in 1X TBS-T) for 1 hour at room temperature. After blocking, the membranes were then incubated overnight with the primary antibody at 4° C. Primary antibodies used in the study with dilutions were: TS (1:2500, rabbit, abcam, ab108995), TNC (1:1000, rabbit, abcam, ab108930), DAX1 (1:1000, rabbit, abcam, ab196649), KRAS (1:1000, rabbit, abcam, ab180772), p-mTOR (1:1000, rabbit, Cell Signaling, 2971S), p-P53 (1:1000, rabbit, Cell Signaling, 9284S), P16 (1:1000, mouse, abcam, ab54210) and β-Actin HRP conjugated (1:10000, Cell Signaling, 12262). Membranes were washed thrice with 1X TBS-T and incubated with the secondary antibody. Secondary antibodies used were: Goat Anti-rabbit IgG-HRP (1:10000, Southern Biotech, 4030-05) and Goat Anti-mouse IgG2b-HRP (1:10000, Southern Biotech, 1091-05). Pierce ECL western blotting solution (Thermo Scientific) was used to detect the protein bands with the development of X-ray films (Thermo Scientific).

### RNA-sequencing

A549 cell line treated with brequinar was harvested for RNA using 700 µl of QIAzol reagent (Qiagen), and the total RNA was isolated and eluted in 40 μl of nuclease free water (VWR) as per the manufacturer’s protocol using miRNeasy kit (Qiagen). 500 ng of total RNA was utilized to construct the sequencing libraries using NEBNext Ultra RNA Library Prep Kit for Illumina as per the manufacturer’s protocol (NEB) and paired-end sequencing was performed with NovaSeq 6000 platform (Illumina).

RNA-seq analysis was performed with the obtained Fastq files from the sequencing platform. Briefly, quality control (QC) analysis was carried out using FastQC program followed by the alignment of the reads to the reference genome GRCh38 release 105 along with the respective human gene annotation file using STAR aligner v2.7.9a. The aligned reads were then counted using the featureCounts function in Rsubread v2.10.5 package in R 4.2.1. Differential gene expression analysis was performed using the raw count reads with DESeq2 v1.36.0 package in R with an adjusted p-value<0.0001 and fold change of 2.

### Single cell RNA-sequencing

Single cell RNA sequencing for parental A549 cell line was performed using 10X Genomics guidelines at the Institute of Human Genetics, Friedrich-Alexander-University of Erlangen-Nürnberg, Erlangen, Germany. Approximately 6000 cells were used and obtained 25,000 reads per cell. In brief, cellranger-4.0.0 pipeline was used to obtain the filtered_feature_bc_matrix data and analyzed further with Seurat v4.2.1 package in R. Pre-processing of the file was performed after quality control metric analysis and filtered the data by eliminating cells with less than 500 genes, UMIs with <2500 & >45000 total number of molecules, >10% mitochondrial genes and with > 5% largest genes. Log_10_ genes per UMI was set at > 0.85 for filtering. Normalization of the counts was done with the library size normalization and selected 2000 variable genes to perform principal component analysis (PCA). For the downstream analysis, cells were regressed out for the variations including mitochondrial genes, cell cycle genes, and genes & UMI counts per cell. With the first 25 principal components, cell clusters were determined using Lovain algorithm with a resolution of 0.4, and tSNE dimensionality reduction method was employed to visualize the cell clusters. AddModuleScore function in Seurat was then applied to assess the percent activation pattern of pyrimidine and its associated signaling or metabolic processes gene-sets for the up-regulated gene-sets in the identified cell clusters.

### Gene enrichment analysis

Gene set enrichment analysis (GSEA) was performed for the patient samples and cancer cell line gene expression profile by categorizing the samples as low and high based on the median of the z-score activation levels of the gene-sets. Signal2Noise metric was used to rank the genes between groups comparison. Significance for the enrichment was set at a nominal p-value < 0.05 and FDR < 0.25. Representational overlap analysis of BRQ-regulated genes identified from RNA-seq analysis was performed with Enrichr web tool (Kuleshov *et al*, 2016) containing in-built gene-set libraries for various biological functions.

### Survival analysis

Samples from TCGA gene expression profiles obtained from cbioportal platform were categorized as low and high based on the median of the signaling pathways gene-set activity score calculated with the difference of up-regulated and down-regulated activation scores in the patient samples. Survival curves were generated based on Kaplan-Meier estimate analysis with log-rank test used to estimate the significant difference between the two categorized groups and hazard ratio was estimated using Cox proportional in R software.

### Gene-sets source

Signaling pathways and metabolic process associated gene-sets were collected from MSigDb v6.2. In total, 247 signaling associated gene-sets and 335 metabolic process associated gene-sets were collected (135 of KEGG and 200 of REACTOME gene-sets). Signaling pathway gene-sets represent oncogenic, tumor suppressor, epigenetic, stem cells, cell cycle and transcription factor mediated pathways. Metabolic process associated gene-sets collectively represent diverse metabolic processes of the cell such as fatty acids, steroids, vitamins, lipids, carbohydrates, organic, inorganic, catabolic and anabolic reactions, amino acids, hormones, nucleotide, enzymatic activities including coenzymes or co-factors, alcohol, amine, drug, acyl chain related metabolisms, biopolymer and macromolecular metabolism. Pyrimidine metabolism gene-set was collected from KEGG database and the polymerase containing genes were removed from the list to have more metabolism-centered analysis.

### Pathway-based activation analysis

Signaling pathway or metabolic process activity score was calculated as described previously (Ramesh & Ganesan, 2016). First, fold difference (log_2_) for each gene in each tumor sample was calculated relative to the median expression value across tumor samples of each cancer subtype as reference. Second, mean and standard deviation for each tumor’s whole gene expression profile was calculated. Third, mean of the fold expression value for each signaling pathway or metabolic processes genes in the gene-sets was calculated for each tumor sample. Next, pathway or metabolic process activity score (z-score) was estimated by subtracting the mean fold expression value of the whole gene expression from the mean fold expression value of the gene-set followed by dividing by standard deviation of the whole gene expression profile of the tumor sample. Finally, a normalized activation score of the gene-set was obtained by multiplying the obtained above score with the square root of the number of genes in the gene-set. In case of pathways containing up- and down-regulated gene-sets, the final normalized activation score was further obtained by subtracting down-score from the up-score. Normalized activation scores were then used to assess the association between PyMet and the signaling or metabolic process gene-sets by Pearson’s correlation method with significance calculation. Percent positive and negative rate of correlations across cancer types were calculated with adjusted p-value < 0.05 and r > 0.3 or r < -0.3. Then, the number of cancer types with PyMet associated gene-sets (signaling or metabolic) was counted as positive or negative after threshold filtering. To strengthen the observed association pattern, a meta-correlation analysis was performed using metacor.DSL function in R for the tumor types satisfying the initial Pearson’s correlation threshold. The number of samples in each dataset was included for the meta-correlation analysis and significantly associated gene-sets were identified with meta-r > 0.3 and meta-r < -0.3 as positively and negatively associated, respectively, with meta-p-value < 0.05. Heatmap representation of the activation scores or correlation values was carried out using R v4.2.1 or GraphPad Prism9 software. A detailed programming script for calculating the pathway activation scores, correlation analysis and meta-analysis is provided in the **Supplementary Method** section. PYPE web application was developed using Shiny package in the R environment.

## Statistical analysis

All statistical analysis was performed using GraphPad Prism9 software or using R v4.2.1 software.

## Data Availability

The RNA-seq profile of brequinar treatment produced in this study was deposited in Gene Expression Omnibus database with the accession number: GSE248686 (https://www.ncbi.nlm.nih.gov/geo/query/acc.cgi?acc=GSE248686)

## Acknowledgements

This work was supported by the Interdisciplinary Center for Clinical Research of the University of Erlangen-Nuremberg, the German Research Foundation (DFG, CE 281/6-1), the Novo Nordisk Foundation (Hallas-Møller Ascending Investigator Grant 0066909), and by the Danish Cancer Society (A18859). Research work of MAS is supported by Lundbeck Foundation, Denmark (R380-2021-1264). Sequencing was performed at the Center for Functional Genomics and Tissue Plasticity, Functional Genomics & Metabolism Research Unit, University of Southern Denmark. The authors thank Tenna P. Mortensen, Maibrith Wishoff and Ronni Nielsen for sequencing assistance.

## Author’s contribution

Conceived the concept: VR and PC; Designed the experiments: VR and PC; Performed the experiments: VR, LP and MAS. Performed computational and statistical analysis: VR and MD; Designed web server: VR, MD and TKD; Wrote the manuscript: VR and PC. All the authors read and approved the final manuscript.

## Supplementary figure legends

**Supplementary Figure S1. Pan-cancer transcriptome analysis has identified novel connections of signaling pathways and metabolic processes with pyrimidine metabolism.**

A-C. Correlation matrix plot of the top positively and negatively associated signaling pathways of PyMet along with KEGG pyrimidine and REACTOME pyrimidine metabolism in BRCA (N = 1082) (A), COAD (N = 592) (B) and UCEC (N = 527) (C) gene expression profiles using pathway-based approach.

D. Heatmap visualization of correlation values between top positively and negatively associated metabolic processes with PyMet across 32 cancer types.

E. Correlation plot depicting the negative association between the activation scores of leucine deprivation and PyMet gene-sets in stomach adenocarcinoma gene expression profile (N = 412).

**Supplementary Figure S2. Pyrimidine association with signaling pathways and metabolic processes were recapitulated in CCLE and parental single cell RNA-sequencing.**

A-C. Gene-set enrichment analysis of KEGG pyrimidine metabolism gene-set in the gene expression profile of cancer cell line encyclopedia (CCLE) panel (N = 917) categorized as low and high based on the ESRRA (A), FLI1-DAX1 (B) or SOX4 (C) gene-set activation levels showed pyrimidine metabolic process enrichment in high-ESRRA (A) or high-FLI1 and DAX1 (B) categorized cell lines and in low-SOX4 (C) categorized cell lines. Ranking of genes with signal2noise metric was used for GSEA.

D-F. Dot plot pattern analysis of negatively associated signaling pathways (D), positively associated metabolic processes (E) and negatively associated metabolic processes (F) of PyMet in the scRNA-seq of parental A549 cells. Percent expression and average expression was estimated using AddmoduleScore function in Seurat.

**Supplementary Figure S3. RNA-seq expression profiling of BRQ treated samples substantiate the pyrimidine associated pathways**

A. MA plot of differentially expressed genes identified in A549 cell line treated with BRQ for 72 h compared to the control (n = 3 per group). Blue dots represent the significantly differentially expressed genes with a fold change of log_2_ (1).

B. Venn diagram representation of overlap analysis of the gene content between pyrimidine metabolism genes and BRQ down-regulated genes. Significance was calculated using the hypergeometric distribution method.

C,D. Enrichr based gene enrichment analysis of BRQ up-regulated genes in MSigDB gene-sets collection (C), and BRQ down-regulated genes overlap with Bioplanet 2019 (D).

E-G. Box plot visualization of decreased activation levels of pyrimidine metabolism (E), and FLI1-DAX1 down-regulated gene-set (F) in the RNA-seq samples of BRQ (1 μM) treated A549 cells compared to the control. TP53 and TP73 target gene-sets were found expressed with BRQ treatment (G). The central band inside the box represents the median value of the data (n = 3) obtained using the lower (bottom) and upper (top) quartile values of the box. The maximum and minimum values of the data are displayed with the vertical lines (whiskers) connecting the box. Significance was calculated using un-paired *t*-test between the BRQ treated cells and the control.

H,I. Gene set enrichment analysis of up-regulated gene-set of TNC signaling pathway (H) and KEGG pyrimidine metabolism (I) in TCGA lung adenocarcinoma (N = 510) patients categorized as BRQ-high and BRQ-low based on the median of BRQ gene-set activation scores. Ranking of genes with signal2noise metric was used for GSEA.

J. Western blot analysis of TNC and TS protein levels in SKMES1 cells treated with brequinar (BRQ) in the indicated dose-dependent manner for 48 h. β-actin was used as an internal control.

K. Western blot analysis of DAX1 and TS protein levels in SKMES1 cells treated with brequinar (BRQ) in the indicated dose-dependent manner for 72 h. β-actin was used as an internal control.

L. Scatter plot of activation scores of KRAS and BRQ gene-sets in lung adenocarcinoma patient samples (N = 510) showed a negative association pattern between KRAS and DHODH inhibitor, BRQ.

**Supplementary Figure S4. Pyrimidine metabolic process and its associated signaling pathways pose drug resistance in pan-cancer analysis.**

A. Correlation matrix plot depicting the associations between drug resistance gene-sets, BRQ, pyrimidine and its associated signaling pathways in bladder carcinoma (N = 407).

B. Scatter plots of the activation scores between doxorubicin drug resistance and pyrimidine metabolism, its associated signaling pathways (HOXA1 and TNC) and BRQ gene-sets in bladder carcinoma (N = 407). Doxorubicin drug resistance showed positive associations with pyrimidine and HOXA1, while negative associations were observed with DHODH inhibitor (BRQ) and TNC signaling pathway.

C. Correlation matrix plot depicting the associations between drug resistance gene-sets, BRQ, pyrimidine and its associated signaling pathways in breast invasive carcinoma (N = 1082).

D. Scatter plots of the activation scores between doxorubicin drug resistance and pyrimidine metabolism, its associated signaling pathways (HOXA1 and TNC) and BRQ gene-sets in breast invasive carcinoma (N = 1082). Doxorubicin drug resistance showed positive associations with pyrimidine and HOXA1, while negative associations were observed with DHODH inhibitor (BRQ) and TNC signaling pathway.

E. Correlation matrix plot depicting the associations between drug resistance gene-sets, BRQ, pyrimidine and its associated signaling pathways in low grade glioma (N = 514).

F. Scatter plots of the activation scores between doxorubicin drug resistance and pyrimidine metabolism, its associated signaling pathways (HOXA1 and TNC) and BRQ gene-sets in low grade glioma (N = 514). Doxorubicin drug resistance showed positive associations with pyrimidine and HOXA1, while negative associations were observed with DHODH inhibitor (BRQ) and TNC signaling pathway.

