## Supplementary Figures for "Pan-cancer analysis of pyrimidine metabolism reveals signaling pathways connections with chemoresistance role"

**Supplementary Figure S1**

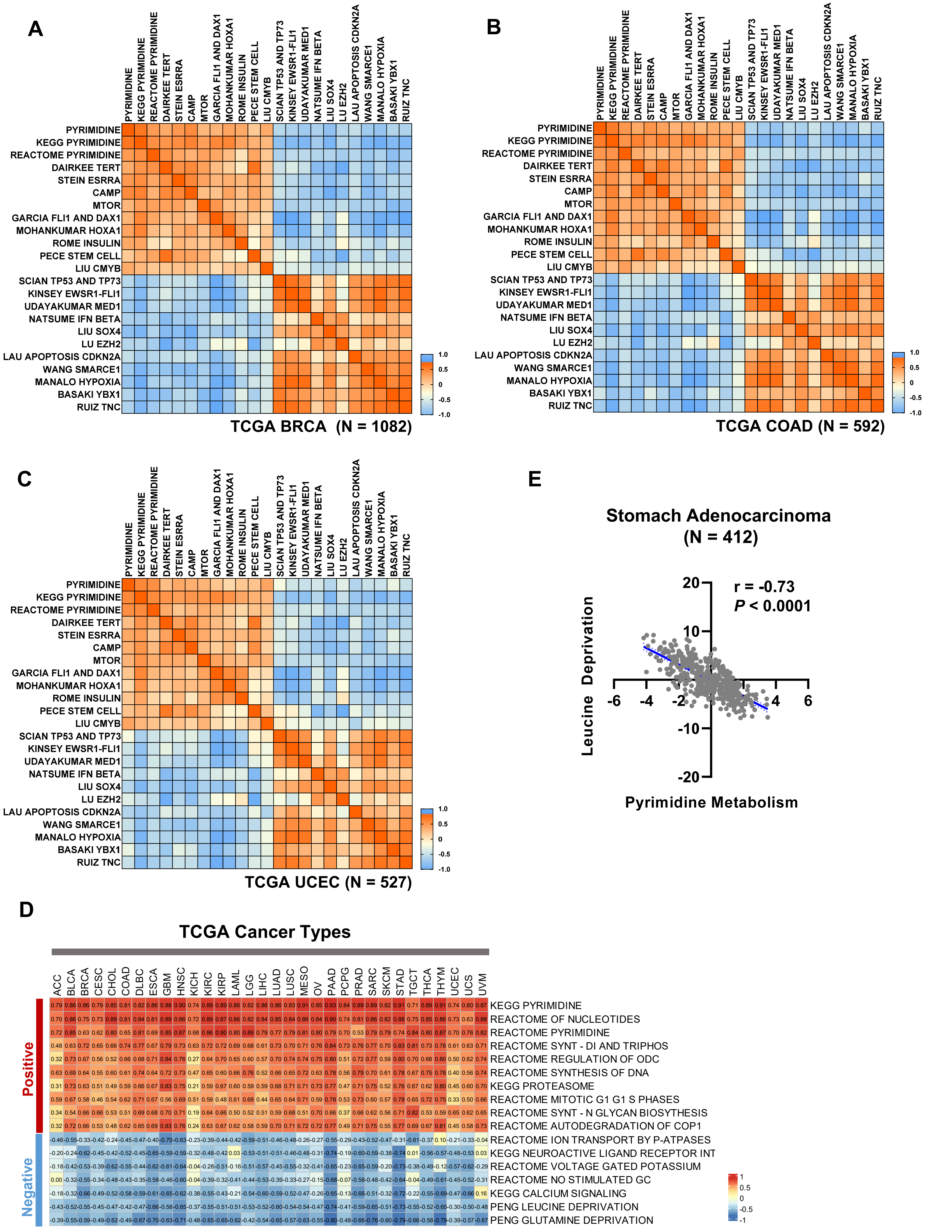

#### Supplementary Figure S2

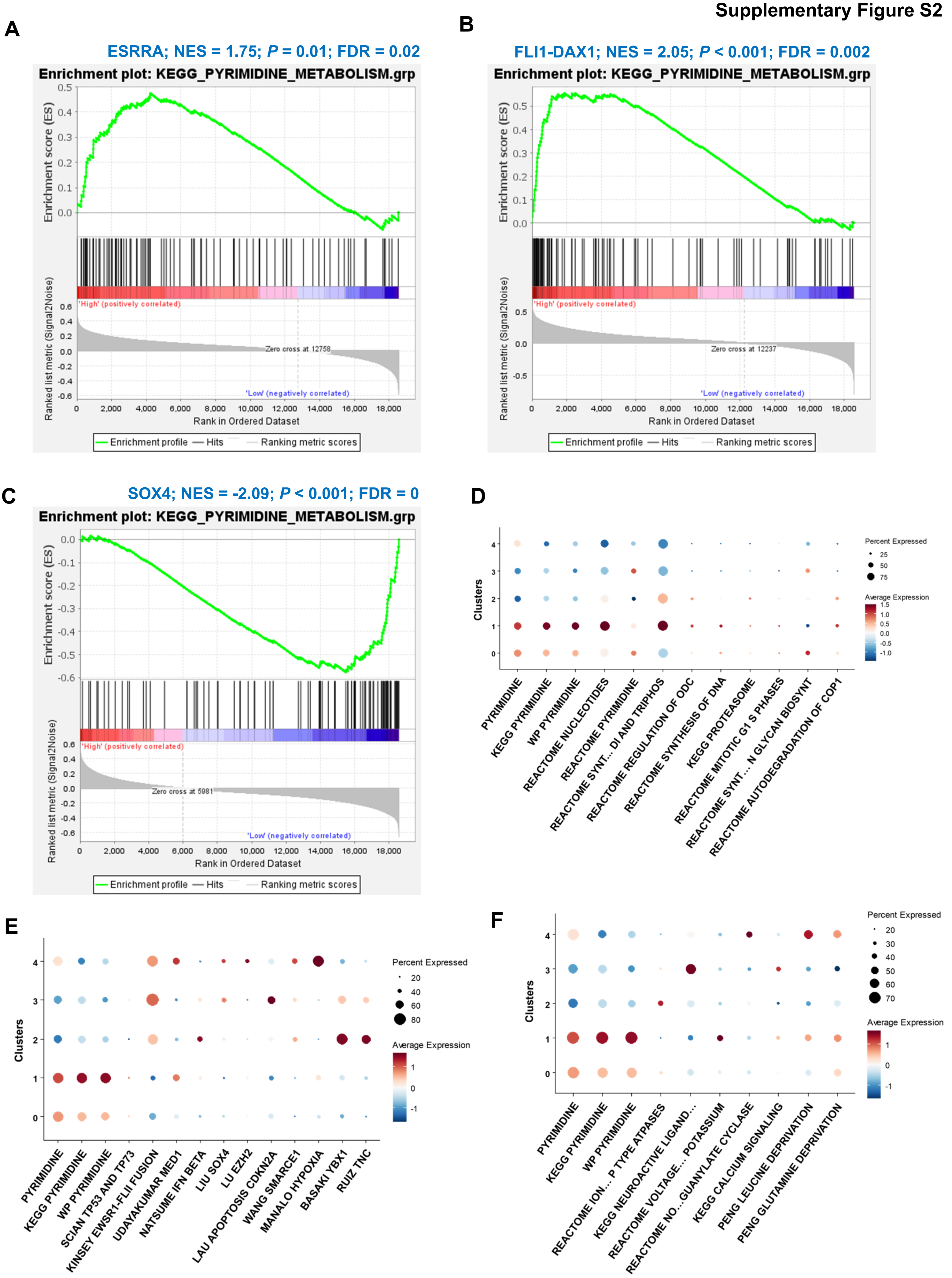

### Supplementary Figure S3

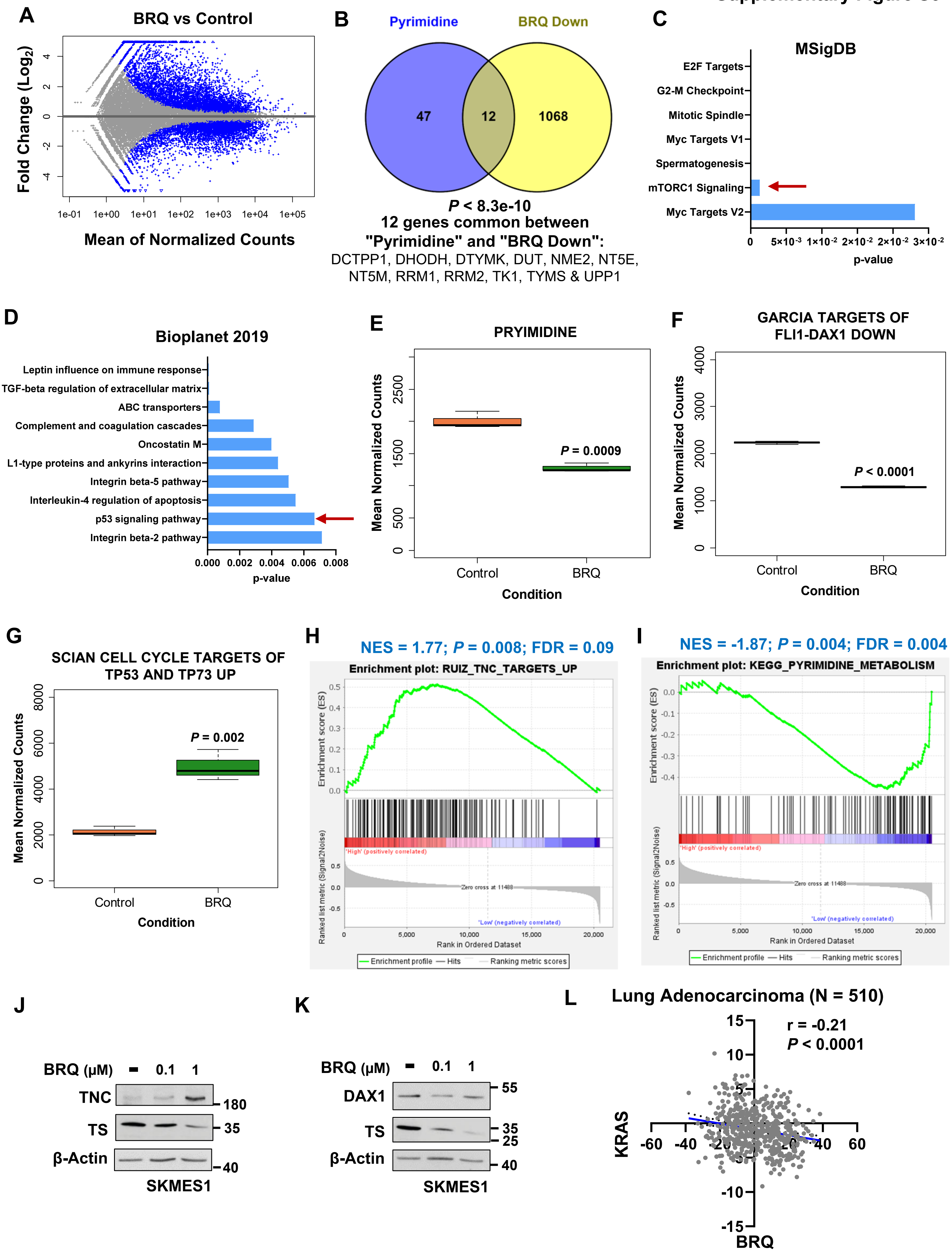

### Supplementary Figure S4

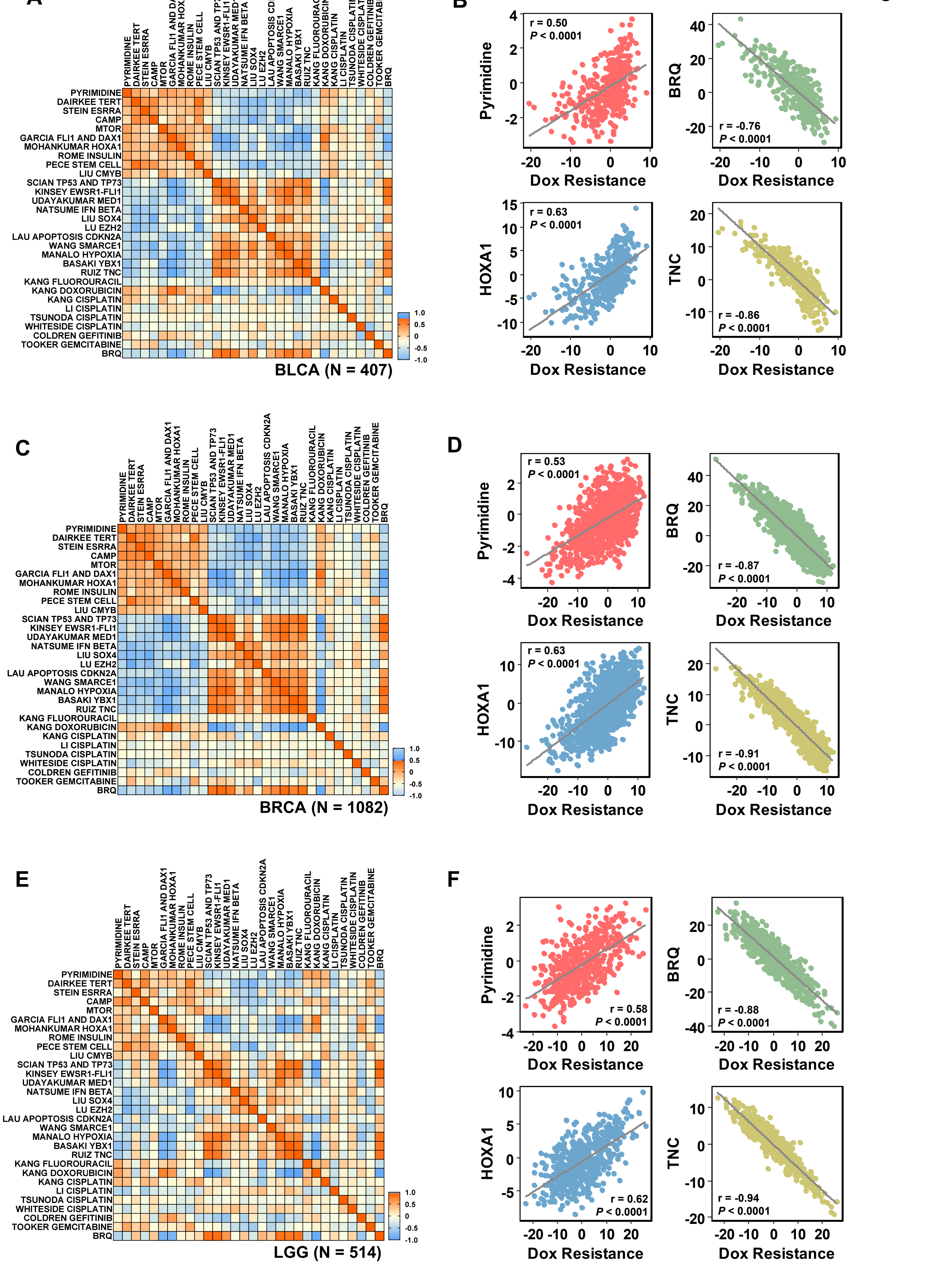
