## Supplementary Tables for "Pan-cancer analysis of pyrimidine metabolism reveals signaling pathways connections with chemoresistance role"

**Supplementary Table S1. List of pyrimidine metabolism genes obtained from KEGG database utilized for the pancancer analysis.**

| | |
|---|---|
| AK3 | NME6 |
| CAD | NME7 |
| CANT1 | NT5C |
| CDA | NT5C1A |
| CMPK1 | NT5C1B |
| CMPK2 | NT5C1B-RDH14 |
| CTPS1 | NT5C2 |
| CTPS2 | NT5C3 |
| DCK | NT5C3A |
| DCTD | NT5C3B |
| DCTPP1 | NT5E |
| DHODH | NT5M |
| DPYD | NUDT2 |
| DPYS | PNP |
| DTYMK | RRM1 |
| DUT | RRM2 |
| ENPP1 | RRM2B |
| ENPP3 | TK1 |
| ENTPD1 | TK2 |
| ENTPD3 | TYMP |
| ENTPD4 | TYMS |
| ENTPD5 | UCK1 |
| ENTPD6 | UCK2 |
| ENTPD8 | UCKL1 |
| NME1 | UMPS |
| NME1-NME2 | UPB1 |
| NME2 | UPP1 |
| NME3 | UPP2 |
| NME4 | UPRT |
| NME5 | |

**Supplementary Table S2. List of 32 cancer types representing 12 different tissue origins, TCGA codes and number of samples in each cancer type used for the signaling pathway and metabolic process analysis in pan-cancer expression profiles**

| Cancer Types | TCGA Code | No. of Samples |
|---|---|---|
| Brain | GBM | 160 |
| Brain | LGG | 514 |
| Breast | BRCA | 1082 |
| Endocrine | ACC | 78 |
| Endocrine | PCPG | 178 |
| Endocrine | THCA | 498 |
| Eye | UVM | 80 |
| Gastrointestinal | CHOL | 36 |
| Gastrointestinal | COAD | 592 |
| Gastrointestinal | ESCA | 181 |
| Gastrointestinal | LIHC | 366 |
| Gastrointestinal | PAAD | 177 |
| Gastrointestinal | STAD | 412 |
| Gynecological | CESC | 294 |
| Gynecological | OV | 300 |
| Gynecological | UCEC | 527 |
| Gynecological | UCS | 57 |
| Haem/ Lymph | DLBC | 48 |
| Haem/ Lymph | LAML | 173 |
| Haem/ Lymph | THYM | 119 |
| Head & Neck | HNSC | 515 |
| Mesenchymal | SARC | 253 |
| Pulmonary | LUAD | 510 |
| Pulmonary | LUSC | 484 |
| Pulmonary | MESO | 87 |
| Skin | SKCM | 443 |
| Urologic | BLCA | 407 |
| Urologic | KICH | 65 |
| Urologic | KIRC | 510 |
| Urologic | KIRP | 283 |
| Urologic | PRAD | 493 |
| Urologic | TGCT | 149 |

**Supplementary Table S3. List of gene-sets collected from MSigDB representing various cellular signaling pathways used for the signaling pathway activation pattern analysis in TCGA pan-cancer expression profiles.**

ACOSTA_PROLIFERATION_INDEPENDENT_MYC_TARGETS_UP
ACOSTA_PROLIFERATION_INDEPENDENT_MYC_TARGETS_DN
AIYAR_COBRA1_TARGETS_DN
AIYAR_COBRA1_TARGETS_UP
ALONSO_METASTASIS_EMT_UP
ALONSO_METASTASIS_EMT_DN
ATF2_UP.V1_UP
ATF2_UP.V1_DN
ALK_DN.V1_DN
ALK_DN.V1_UP
ATM_DN.V1_DN
ATM_DN.V1_UP
AZARE_NEOPLASTIC_TRANSFORMATION_BY_STAT3_UP
AZARE_NEOPLASTIC_TRANSFORMATION_BY_STAT3_DN
BAE_BRCA1_TARGETS_UP
BAE_BRCA1_TARGETS_DN
BASAKI_YBX1_TARGETS_DN
BASAKI_YBX1_TARGETS_UP
BASSO_CD40_SIGNALING_UP
BASSO_CD40_SIGNALING_DN
BCAT_BILD_ET_AL_UP
BCAT_BILD_ET_AL_DN
BEGUM_TARGETS_OF_PAX3_FOXO1_FUSION_UP
BEGUM_TARGETS_OF_PAX3_FOXO1_FUSION_DN
BEIER_GLIOMA_STEM_CELL_UP
BEIER_GLIOMA_STEM_CELL_DN
BILBAN_B_CLL_LPL_UP
BILBAN_B_CLL_LPL_DN
BMI1_DN.V1_DN
BMI1_DN.V1_UP
BMI1_DN_MEL18_DN.V1_DN
BMI1_DN_MEL18_DN.V1_UP
BOQUEST_STEM_CELL_CULTURED_VS_FRESH_UP
BOQUEST_STEM_CELL_CULTURED_VS_FRESH_DN
BOQUEST_STEM_CELL_UP
BOQUEST_STEM_CELL_DN
BRCA1_DN.V1_DN
BRCA1_DN.V1_UP
BUSA_SAM68_TARGETS_DN
BUSA_SAM68_TARGETS_UP
CAMP_UP.V1_UP
CAMP_UP.V1_DN
CEBALLOS_TARGETS_OF_TP53_AND_MYC_UP
CEBALLOS_TARGETS_OF_TP53_AND_MYC_DN
CERVERA_SDHB_TARGETS_1_DN
CERVERA_SDHB_TARGETS_1_UP
CHANG_POU5F1_TARGETS_UP
CHANG_POU5F1_TARGETS_DN
CHARAFE_BREAST_CANCER_BASAL_VS_MESENCHYMAL_DN
CHARAFE_BREAST_CANCER_BASAL_VS_MESENCHYMAL_UP
CHARAFE_BREAST_CANCER_LUMINAL_VS_BASAL_UP
CHARAFE_BREAST_CANCER_LUMINAL_VS_BASAL_DN
CHARAFE_BREAST_CANCER_LUMINAL_VS_MESENCHYMAL_DN
CHARAFE_BREAST_CANCER_LUMINAL_VS_MESENCHYMAL_UP
CHEN_HOXA5_TARGETS_6HR_UP
CHEN_HOXA5_TARGETS_6HR_DN
CHEN_HOXA5_TARGETS_9HR_UP
CHEN_HOXA5_TARGETS_9HR_DN
CHOW_RASSF1_TARGETS_UP
CHOW_RASSF1_TARGETS_DN
CHUANG_OXIDATIVE_STRESS_RESPONSE_UP
CHUANG_OXIDATIVE_STRESS_RESPONSE_DN
COLLER_MYC_TARGETS_UP
COLLER_MYC_TARGETS_DN
COULOUARN_TEMPORAL_TGFB1_SIGNATURE_UP
COULOUARN_TEMPORAL_TGFB1_SIGNATURE_DN
CREIGHTON_AKT1_SIGNALING_VIA_MTOR_UP
CREIGHTON_AKT1_SIGNALING_VIA_MTOR_DN
CROONQUIST_IL6_DEPRIVATION_DN
CROONQUIST_IL6_DEPRIVATION_UP
CTIP_DN.V1_DN
CTIP_DN.V1_UP
CYCLIN_D1_UP.V1_UP
CYCLIN_D1_UP.V1_DN
DAIRKEE_TERT_TARGETS_UP
DAIRKEE_TERT_TARGETS_DN
DASU_IL6_SIGNALING_SCAR_UP
DASU_IL6_SIGNALING_SCAR_DN
DASU_IL6_SIGNALING_UP
DASU_IL6_SIGNALING_DN
DAUER_STAT3_TARGETS_UP
DAUER_STAT3_TARGETS_DN
DAVICIONI_TARGETS_OF_PAX_FOXO1_FUSIONS_UP
DAVICIONI_TARGETS_OF_PAX_FOXO1_FUSIONS_DN
DELPUECH_FOXO3_TARGETS_UP
DELPUECH_FOXO3_TARGETS_DN
DER_IFN_ALPHA_RESPONSE_UP
DER_IFN_ALPHA_RESPONSE_DN
DER_IFN_BETA_RESPONSE_UP
DER_IFN_BETA_RESPONSE_DN
DER_IFN_GAMMA_RESPONSE_UP
DER_IFN_GAMMA_RESPONSE_DN
DE_YY1_TARGETS_DN
DE_YY1_TARGETS_UP
DITTMER_PTHLH_TARGETS_DN
DITTMER_PTHLH_TARGETS_UP
DORSAM_HOXA9_TARGETS_UP
DORSAM_HOXA9_TARGETS_DN
DOUGLAS_BMI1_TARGETS_DN
DOUGLAS_BMI1_TARGETS_UP
DUNNE_TARGETS_OF_AML1_MTG8_FUSION_DN
DUNNE_TARGETS_OF_AML1_MTG8_FUSION_UP
E2F3_UP.V1_UP
E2F3_UP.V1_DN
EBAUER_TARGETS_OF_PAX3_FOXO1_FUSION_DN
EBAUER_TARGETS_OF_PAX3_FOXO1_FUSION_UP
EGFR_UP.V1_UP
EGFR_UP.V1_DN
EIF4E_UP
EIF4E_DN
ELVIDGE_HIF1A_AND_HIF2A_TARGETS_DN
ELVIDGE_HIF1A_AND_HIF2A_TARGETS_UP
ELVIDGE_HIF1A_TARGETS_DN
ELVIDGE_HIF1A_TARGETS_UP
ELVIDGE_HYPOXIA_BY_DMOG_UP
ELVIDGE_HYPOXIA_BY_DMOG_DN
ELVIDGE_HYPOXIA_UP
ELVIDGE_HYPOXIA_DN
ERB2_UP.V1_UP
ERB2_UP.V1_DN
FEVR_CTNNB1_TARGETS_DN
FEVR_CTNNB1_TARGETS_UP
FORTSCHEGGER_PHF8_TARGETS_DN
FORTSCHEGGER_PHF8_TARGETS_UP
FUJII_YBX1_TARGETS_DN
FUJII_YBX1_TARGETS_UP

FURUKAWA_DUSP6_TARGETS_PCI35_UP
FURUKAWA_DUSP6_TARGETS_PCI35_DN
GAL_LEUKEMIC_STEM_CELL_UP
GAL_LEUKEMIC_STEM_CELL_DN
GARCIA_TARGETS_OF_FLI1_AND_DAX1_DN
GARCIA_TARGETS_OF_FLI1_AND_DAX1_UP
GARY_CD5_TARGETS_UP
GARY_CD5_TARGETS_DN
GHO_ATF5_TARGETS_UP
GHO_ATF5_TARGETS_DN
GOUYER_TATI_TARGETS_UP
GOUYER_TATI_TARGETS_DN
GOZGIT_ESR1_TARGETS_DN
GOZGIT_ESR1_TARGETS_UP
GRABARCZYK_BCL11B_TARGETS_DN
GRABARCZYK_BCL11B_TARGETS_UP
GRANDVAUX_IRF3_TARGETS_UP
GRANDVAUX_IRF3_TARGETS_DN
GUENTHER_GROWTH_SPHERICAL_VS_ADHERENT_UP
GUENTHER_GROWTH_SPHERICAL_VS_ADHERENT_DN
GU_PDEF_TARGETS_DN
GU_PDEF_TARGETS_UP
HALMOS_CEBPA_TARGETS_UP
HALMOS_CEBPA_TARGETS_DN
HAN_SATB1_TARGETS_DN
HAN_SATB1_TARGETS_UP
HASINA_NOL7_TARGETS_UP
HASINA_NOL7_TARGETS_DN
HENDRICKS_SMARCA4_TARGETS_UP
HENDRICKS_SMARCA4_TARGETS_DN
HINATA_NFKB_TARGETS_KERATINOCYTE_UP
HINATA_NFKB_TARGETS_KERATINOCYTE_DN
HIRSCH_CELLULAR_TRANSFORMATION_SIGNATURE_UP
HIRSCH_CELLULAR_TRANSFORMATION_SIGNATURE_DN
HOEBEKE_LYMPHOID_STEM_CELL_UP
HOEBEKE_LYMPHOID_STEM_CELL_DN
HOEGERKORP_CD44_TARGETS_DIRECT_UP
HOEGERKORP_CD44_TARGETS_DIRECT_DN
HOEGERKORP_CD44_TARGETS_TEMPORAL_UP
HOEGERKORP_CD44_TARGETS_TEMPORAL_DN
HOELZEL_NF1_TARGETS_DN
HOELZEL_NF1_TARGETS_UP
HOLLMANN_APOPTOSIS_VIA_CD40_UP
HOLLMANN_APOPTOSIS_VIA_CD40_DN
HOOI_ST7_TARGETS_UP
HOOI_ST7_TARGETS_DN
HORIUCHI_WTAP_TARGETS_DN
HORIUCHI_WTAP_TARGETS_UP
HOXA9_DN.V1_DN
HOXA9_DN.V1_UP
HUANG_FOXA2_TARGETS_UP
HUANG_FOXA2_TARGETS_DN
IGARASHI_ATF4_TARGETS_DN
IGARASHI_ATF4_TARGETS_UP
IL15_UP.V1_UP
IL15_UP.V1_DN
IL21_UP.V1_UP
IL21_UP.V1_DN
IL2_UP.V1_UP
IL2_UP.V1_DN
ITO_PTTG1_TARGETS_DN
ITO_PTTG1_TARGETS_UP
JAATINEN_HEMATOPOIETIC_STEM_CELL_UP
JAATINEN_HEMATOPOIETIC_STEM_CELL_DN
JAK2_DN.V1_DN
JAK2_DN.V1_UP
JAZAG_TGFB1_SIGNALING_UP
JAZAG_TGFB1_SIGNALING_DN
JAZAG_TGFB1_SIGNALING_VIA_SMAD4_UP
JAZAG_TGFB1_SIGNALING_VIA_SMAD4_DN
JEON_SMAD6_TARGETS_DN
JEON_SMAD6_TARGETS_UP
JIANG_TIP30_TARGETS_UP
JIANG_TIP30_TARGETS_DN
JOHNSTONE_PARVB_TARGETS_1_UP
JOHNSTONE_PARVB_TARGETS_1_DN
JOHNSTONE_PARVB_TARGETS_2_UP
JOHNSTONE_PARVB_TARGETS_2_DN
JOHNSTONE_PARVB_TARGETS_3_UP
JOHNSTONE_PARVB_TARGETS_3_DN
KANG_IMMORTALIZED_BY_TERT_UP
KANG_IMMORTALIZED_BY_TERT_DN
KANNAN_TP53_TARGETS_UP
KANNAN_TP53_TARGETS_DN
KIM_PTEN_TARGETS_DN
KIM_PTEN_TARGETS_UP
KINSEY_TARGETS_OF_EWSR1_FLII_FUSION_DN
KINSEY_TARGETS_OF_EWSR1_FLII_FUSION_UP
KOYAMA_SEMA3B_TARGETS_UP
KOYAMA_SEMA3B_TARGETS_DN
KRAS.300_UP.V1_UP
KRAS.300_UP.V1_DN
KRAS.50_UP.V1_UP
KRAS.50_UP.V1_DN
KRAS.600_UP.V1_UP
KRAS.600_UP.V1_DN
KRAS.600.LUNG.BREAST_UP.V1_UP
KRAS.600.LUNG.BREAST_UP.V1_DN
KRAS.AMP.LUNG_UP.V1_UP
KRAS.AMP.LUNG_UP.V1_DN
KRAS.BREAST_UP.V1_UP
KRAS.BREAST_UP.V1_DN
KRAS.DF.V1_UP
KRAS.DF.V1_DN
KRAS.KIDNEY_UP.V1_UP
KRAS.KIDNEY_UP.V1_DN
KRAS.LUNG.BREAST_UP.V1_UP
KRAS.LUNG.BREAST_UP.V1_DN
KRAS.LUNG_UP.V1_UP
KRAS.LUNG_UP.V1_DN
KRAS.PROSTATE_UP.V1_UP
KRAS.PROSTATE_UP.V1_DN
KRASNOSELSKAYA_ILF3_TARGETS_UP
KRASNOSELSKAYA_ILF3_TARGETS_DN
KREPPEL_CD99_TARGETS_DN
KREPPEL_CD99_TARGETS_UP
LAU_APOPTOSIS_CDKN2A_DN
LAU_APOPTOSIS_CDKN2A_UP
LEE_NEURAL_CREST_STEM_CELL_UP
LEE_NEURAL_CREST_STEM_CELL_DN
LEF1_UP.V1_UP
LEF1_UP.V1_DN
LIANG_SILENCED_BY_METHYLATION_UP
LIANG_SILENCED_BY_METHYLATION_DN
LINDVALL_IMMORTALIZED_BY_TERT_UP
LINDVALL_IMMORTALIZED_BY_TERT_DN
LIU_CDX2_TARGETS_UP
LIU_CDX2_TARGETS_DN
LIU_CMYB_TARGETS_UP
LIU_CMYB_TARGETS_DN
LIU_IL13_MEMORY_MODEL_UP
LIU_IL13_MEMORY_MODEL_DN
LIU_SOX4_TARGETS_UP
LIU_SOX4_TARGETS_DN

LIU_TARGETS_OF_VMYB_VS_CMYB_UP
LIU_TARGETS_OF_VMYB_VS_CMYB_DN
LTE2_UP.V1_UP
LTE2_UP.V1_DN
LUCAS_HNF4A_TARGETS_UP
LUCAS_HNF4A_TARGETS_DN
LU_EZH2_TARGETS_DN
LU_EZH2_TARGETS_UP
MACLACHLAN_BRCA1_TARGETS_UP
MACLACHLAN_BRCA1_TARGETS_DN
MAHAJAN_RESPONSE_TO_IL1A_UP
MAHAJAN_RESPONSE_TO_IL1A_DN
MAINA_VHL_TARGETS_UP
MAINA_VHL_TARGETS_DN
MANALO_HYPOXIA_UP
MANALO_HYPOXIA_DN
MARKS_HDAC_TARGETS_UP
MARKS_HDAC_TARGETS_DN
MARZEC_IL2_SIGNALING_UP
MARZEC_IL2_SIGNALING_DN
MEK_UP.V1_UP
MEK_UP.V1_DN
MEL18_DN.V1_DN
MEL18_DN.V1_UP
MELLMAN_TUT1_TARGETS_DN
MELLMAN_TUT1_TARGETS_UP
MIKHAYLOVA_OXIDATIVE_STRESS_RESPONSE_VIA_VHL_UP
MIKHAYLOVA_OXIDATIVE_STRESS_RESPONSE_VIA_VHL_DN
MIYAGAWA_TARGETS_OF_EWSR1_ETS_FUSIONS_UP
MIYAGAWA_TARGETS_OF_EWSR1_ETS_FUSIONS_DN
MIZUKAMI_HYPOXIA_UP
MIZUKAMI_HYPOXIA_DN
MOHANKUMAR_HOXA1_TARGETS_UP
MOHANKUMAR_HOXA1_TARGETS_DN
MOLENAAR_TARGETS_OF_CCND1_AND_CDK4_DN
MOLENAAR_TARGETS_OF_CCND1_AND_CDK4_UP
MTOR_UP.N4.V1_UP
MTOR_UP.N4.V1_DN
MYC_UP.V1_UP
MYC_UP.V1_DN
NAGASHIMA_NRG1_SIGNALING_UP
NAGASHIMA_NRG1_SIGNALING_DN
NATSUME_RESPONSE_TO_INTERFERON_BETA_UP
NATSUME_RESPONSE_TO_INTERFERON_BETA_DN
NEWMAN_ERCC6_TARGETS_UP
NEWMAN_ERCC6_TARGETS_DN
NGUYEN_NOTCH1_TARGETS_UP
NGUYEN_NOTCH1_TARGETS_DN
NOJIMA_SFRP2_TARGETS_UP
NOJIMA_SFRP2_TARGETS_DN
NUYTTEN_EZH2_TARGETS_DN
NUYTTEN_EZH2_TARGETS_UP
NUYTTEN_NIPP1_TARGETS_DN
NUYTTEN_NIPP1_TARGETS_UP
ODONNELL_TARGETS_OF_MYC_AND_TFRC_DN
ODONNELL_TARGETS_OF_MYC_AND_TFRC_UP
ODONNELL_TFRC_TARGETS_DN
ODONNELL_TFRC_TARGETS_UP
OLSSON_E2F3_TARGETS_DN
OLSSON_E2F3_TARGETS_UP
ONDER_CDH1_TARGETS_1_DN
ONDER_CDH1_TARGETS_1_UP
ONDER_CDH1_TARGETS_2_DN
ONDER_CDH1_TARGETS_2_UP
ONDER_CDH1_TARGETS_3_DN
ONDER_CDH1_TARGETS_3_UP
OSADA_ASCL1_TARGETS_UP
OSADA_ASCL1_TARGETS_DN
OXFORD_RALA_AND_RALB_TARGETS_DN
OXFORD_RALA_AND_RALB_TARGETS_UP
OXFORD_RALA_TARGETS_DN
OXFORD_RALA_TARGETS_UP
OXFORD_RALB_TARGETS_DN
OXFORD_RALB_TARGETS_UP
P53_DN.V2_DN
P53_DN.V2_UP
PARENT_MTOR_SIGNALING_UP
PARENT_MTOR_SIGNALING_DN
PASTURAL_RIZ1_TARGETS_UP
PASTURAL_RIZ1_TARGETS_DN
PDGF_ERK_DN.V1_UP
PDGF_ERK_DN.V1_DN
PDGF_UP.V1_UP
PDGF_UP.V1_DN
PEART_HDAC_PROLIFERATION_CLUSTER_UP
PEART_HDAC_PROLIFERATION_CLUSTER_DN
PECE_MAMMARY_STEM_CELL_UP
PECE_MAMMARY_STEM_CELL_DN
PETROVA_PROX1_TARGETS_UP
PETROVA_PROX1_TARGETS_DN
PHONG_TNF_TARGETS_UP
PHONG_TNF_TARGETS_DN
PIEPOLI_LGI1_TARGETS_UP
PIEPOLI_LGI1_TARGETS_DN
PIGF_UP.V1_UP
PIGF_UP.V1_DN
PRAMOONJAGO_SOX4_TARGETS_DN
PRAMOONJAGO_SOX4_TARGETS_UP
PRC1_BMI_UP.V1_DN
PRC1_BMI_UP.V1_UP
PRC2_EED_UP.V1_DN
PRC2_EED_UP.V1_UP
PRC2_EZH2_UP.V1_DN
PRC2_EZH2_UP.V1_UP
PRC2_SUZ12_UP.V1_DN
PRC2_SUZ12_UP.V1_UP
PTEN_DN.V1_DN
PTEN_DN.V1_UP
PTEN_DN.V2_DN
PTEN_DN.V2_UP
PURBEY_TARGETS_OF_CTBP1_AND_SATB1_DN
PURBEY_TARGETS_OF_CTBP1_AND_SATB1_UP
PURBEY_TARGETS_OF_CTBP1_NOT_SATB1_DN
PURBEY_TARGETS_OF_CTBP1_NOT_SATB1_UP
RADAEVA_RESPONSE_TO_IFNA1_UP
RADAEVA_RESPONSE_TO_IFNA1_DN
RAF_UP.V1_UP
RAF_UP.V1_DN
RELA_DN.V1_DN
RELA_DN.V1_UP
RODRIGUES_DCC_TARGETS_UP
RODRIGUES_DCC_TARGETS_DN
RODRIGUES_NTN1_TARGETS_UP
RODRIGUES_NTN1_TARGETS_DN
ROETH_TERT_TARGETS_UP
ROETH_TERT_TARGETS_DN
ROME_INSULIN_TARGETS_IN_MUSCLE_UP
ROME_INSULIN_TARGETS_IN_MUSCLE_DN
RORIE_TARGETS_OF_EWSR1_FLI1_FUSION_UP
RORIE_TARGETS_OF_EWSR1_FLI1_FUSION_DN
ROZANOV_MMP14_TARGETS_UP
ROZANOV_MMP14_TARGETS_DN
RUIZ_TNC_TARGETS_UP
RUIZ_TNC_TARGETS_DN

RUTELLA_RESPONSE_TO_CSF2RB_AND_IL4_UP
RUTELLA_RESPONSE_TO_CSF2RB_AND_IL4_DN
RUTELLA_RESPONSE_TO_HGF_UP
RUTELLA_RESPONSE_TO_HGF_DN
RUTELLA_RESPONSE_TO_HGF_VS_CSF2RB_AND_IL4_UP
RUTELLA_RESPONSE_TO_HGF_VS_CSF2RB_AND_IL4_DN
SAGIV_CD24_TARGETS_DN
SAGIV_CD24_TARGETS_UP
SANA_RESPONSE_TO_IFNG_UP
SANA_RESPONSE_TO_IFNG_DN
SANA_TNF_SIGNALING_UP
SANA_TNF_SIGNALING_DN
SARRIO_EPITHELIAL_MESENCHYMAL_TRANSITION_UP
SARRIO_EPITHELIAL_MESENCHYMAL_TRANSITION_DN
SCHRAMM_INHBA_TARGETS_UP
SCHRAMM_INHBA_TARGETS_DN
SCHUHMACHER_MYC_TARGETS_UP
SCHUHMACHER_MYC_TARGETS_DN
SCHURINGA_STAT5A_TARGETS_UP
SCHURINGA_STAT5A_TARGETS_DN
SCIAN_CELL_CYCLE_TARGETS_OF_TP53_AND_TP73_UP
SCIAN_CELL_CYCLE_TARGETS_OF_TP53_AND_TP73_DN
SCIAN_INVERSED_TARGETS_OF_TP53_AND_TP73_UP
SCIAN_INVERSED_TARGETS_OF_TP53_AND_TP73_DN
SCIBETTA_KDM5B_TARGETS_UP
SCIBETTA_KDM5B_TARGETS_DN
SENESE_HDAC1_AND_HDAC2_TARGETS_DN
SENESE_HDAC1_AND_HDAC2_TARGETS_UP
SENESE_HDAC1_TARGETS_DN
SENESE_HDAC1_TARGETS_UP
SENESE_HDAC2_TARGETS_DN
SENESE_HDAC2_TARGETS_UP
SENESE_HDAC3_TARGETS_DN
SENESE_HDAC3_TARGETS_UP
SHIRAISHI_PLZF_TARGETS_UP
SHIRAISHI_PLZF_TARGETS_DN
SHI_SPARC_TARGETS_DN
SHI_SPARC_TARGETS_UP
SIRNA_EIF4GI_UP
SIRNA_EIF4GI_DN
SMITH_TERT_TARGETS_UP
SMITH_TERT_TARGETS_DN
SRC_UP.V1_UP
SRC_UP.V1_DN
STEIN_ESRRA_TARGETS_RESPONSIVE_TO_ESTROGEN_UP
STEIN_ESRRA_TARGETS_RESPONSIVE_TO_ESTROGEN_DN
STEIN_ESRRA_TARGETS_UP
STEIN_ESRRA_TARGETS_DN
STK33_NOMO_DN
STK33_NOMO_UP
STK33_SKM_DN
STK33_SKM_UP
STK33_DN
STK33_UP
STREICHER_LSM1_TARGETS_UP
STREICHER_LSM1_TARGETS_DN
SWEET_KRAS_TARGETS_DN
SWEET_KRAS_TARGETS_UP
TANG_SENESCENCE_TP53_TARGETS_DN
TANG_SENESCENCE_TP53_TARGETS_UP
TAVOR_CEBPA_TARGETS_UP
TAVOR_CEBPA_TARGETS_DN
TGFB_UP.V1_UP
TGFB_UP.V1_DN
TSAI_DNAJB4_TARGETS_UP
TSAI_DNAJB4_TARGETS_DN
UDAYAKUMAR_MED1_TARGETS_DN
UDAYAKUMAR_MED1_TARGETS_UP
VANTVEER_BREAST_CANCER_ESR1_UP
VANTVEER_BREAST_CANCER_ESR1_DN
VEGF_A_UP.V1_UP
VEGF_A_UP.V1_DN
VETTER_TARGETS_OF_PRKCA_AND_ETS1_DN
VETTER_TARGETS_OF_PRKCA_AND_ETS1_UP
WANG_CLIM2_TARGETS_DN
WANG_CLIM2_TARGETS_UP
WANG_LMO4_TARGETS_UP
WANG_LMO4_TARGETS_DN
WANG_SMARCE1_TARGETS_UP
WANG_SMARCE1_TARGETS_DN
WEINMANN_ADAPTATION_TO_HYPOXIA_UP
WEINMANN_ADAPTATION_TO_HYPOXIA_DN
WELCSH_BRCA1_TARGETS_UP
WELCSH_BRCA1_TARGETS_DN
WIERENGA_STAT5A_TARGETS_UP
WIERENGA_STAT5A_TARGETS_DN
XU_GH1_AUTOCRINE_TARGETS_UP
XU_GH1_AUTOCRINE_TARGETS_DN
XU_GH1_EXOGENOUS_TARGETS_UP
XU_GH1_EXOGENOUS_TARGETS_DN
XU_HGF_SIGNALING_NOT_VIA_AKT1_48HR_UP
XU_HGF_SIGNALING_NOT_VIA_AKT1_48HR_DN
XU_HGF_TARGETS_INDUCED_BY_AKT1_48HR_UP
XU_HGF_TARGETS_INDUCED_BY_AKT1_48HR_DN
XU_HGF_TARGETS_REPRESSED_BY_AKT1_UP
XU_HGF_TARGETS_REPRESSED_BY_AKT1_DN
YAP1_UP
YAP1_DN
ZHU_SKIL_TARGETS_DN
ZHU_SKIL_TARGETS_UP

**Supplementary Table S4. List of metabolic processes gene-sets (KEGG and REACTOME) collected from MSigDB representing various metabolic processes used for the metabolic process activation pattern analysis in TCGA pan-cancer expression profiles.**

PENG_GLUCOSE_DEPRIVATION_UP
PENG_GLUCOSE_DEPRIVATION_DN
PENG_GLUTAMINE_DEPRIVATION_UP
PENG_GLUTAMINE_DEPRIVATION_DN
PENG_LEUCINE_DEPRIVATION_UP
PENG_LEUCINE_DEPRIVATION_DN
KEGG_GLYCOLYSIS_GLUCONEOGENESIS
KEGG_CITRATE_CYCLE_TCA_CYCLE
KEGG_PENTOSE_PHOSPHATE_PATHWAY
KEGG_PENTOSE_AND_GLUCURONATE_INTERCONVERSIONS
KEGG_FRUCTOSE_AND_MANNOSE_METABOLISM
KEGG_GALACTOSE_METABOLISM
KEGG_ASCORBATE_AND_ALDARATE_METABOLISM
KEGG_FATTY_ACID_METABOLISM
KEGG_STEROID_BIOSYNTHESIS
KEGG_PRIMARY_BILE_ACID_BIOSYNTHESIS
KEGG_STEROID_HORMONE_BIOSYNTHESIS
KEGG_OXIDATIVE_PHOSPHORYLATION
KEGG_PURINE_METABOLISM
KEGG_PYRIMIDINE_METABOLISM
KEGG_ALANINE_ASPARTATE_AND_GLUTAMATE_METABOLISM
KEGG_GLYCINE_SERINE_AND_THREONINE_METABOLISM
KEGG_CYSTEINE_AND_METHIONINE_METABOLISM
KEGG_VALINE_LEUCINE_AND_ISOLEUCINE_DEGRADATION
KEGG_VALINE_LEUCINE_AND_ISOLEUCINE_BIOSYNTHESIS
KEGG_LYSINE_DEGRADATION
KEGG_ARGININE_AND_PROLINE_METABOLISM
KEGG_HISTIDINE_METABOLISM
KEGG_TYROSINE_METABOLISM
KEGG_PHENYLALANINE_METABOLISM
KEGG_TRYPTOPHAN_METABOLISM
KEGG_BETA_ALANINE_METABOLISM
KEGG_TAURINE_AND_HYPOTAURINE_METABOLISM
KEGG_SELENOAMINO_ACID_METABOLISM
KEGG_GLUTATHIONE_METABOLISM
KEGG_STARCH_AND_SUCROSE_METABOLISM
KEGG_N_GLYCAN_BIOSYNTHESIS
KEGG_OTHER_GLYCAN_DEGRADATION
KEGG_O_GLYCAN_BIOSYNTHESIS
KEGG_AMINO_SUGAR_AND_NUCLEOTIDE_SUGAR_METABOLISM
KEGG_GLYCOSAMINOGLYCAN_DEGRADATION
KEGG_GLYCOSAMINOGLYCAN_BIOSYNTHESIS_CHONDROITIN_SULFATE
KEGG_GLYCOSAMINOGLYCAN_BIOSYNTHESIS_KERATAN_SULFATE
KEGG_GLYCOSAMINOGLYCAN_BIOSYNTHESIS_HEPARAN_SULFATE
KEGG_GLYCEROLIPID_METABOLISM
KEGG_INOSITOL_PHOSPHATE_METABOLISM
KEGG_GLYCOSYLPHOSPHATIDYLINOSITOL_GPI_ANCHOR_BIOSYNTHESIS
KEGG_GLYCEROPHOSPHOLIPID_METABOLISM
KEGG_ETHER_LIPID_METABOLISM
KEGG_ARACHIDONIC_ACID_METABOLISM
KEGG_LINOLEIC_ACID_METABOLISM
KEGG_ALPHA_LINOLENIC_ACID_METABOLISM
KEGG_SPHINGOLIPID_METABOLISM
KEGG_GLYCOSPHINGOLIPID_BIOSYNTHESIS_LACTO_AND_NEOLACTO_SERIES
KEGG_GLYCOSPHINGOLIPID_BIOSYNTHESIS_GLOBO_SERIES

KEGG_GLYCOSPHINGOLIPID_BIOSYNTHESIS_GANGLIO_SERIES
KEGG_PYRUVATE_METABOLISM
KEGG_GLYOXYLATE_AND_DICARBOXYLATE_METABOLISM
KEGG_PROPANOATE_METABOLISM
KEGG_BUTANOATE_METABOLISM
KEGG_ONE_CARBON_POOL_BY_FOLATE
KEGG_RIBOFLAVIN_METABOLISM
KEGG_NICOTINATE_AND_NICOTINAMIDE_METABOLISM
KEGG_PANTOTHENATE_AND_COA_BIOSYNTHESIS
KEGG_FOLATE_BIOSYNTHESIS
KEGG_RETINOL_METABOLISM
KEGG_TERPENOID_BACKBONE_BIOSYNTHESIS
KEGG_LIMONENE_AND_PINENE_DEGRADATION
KEGG_NITROGEN_METABOLISM
KEGG_SULFUR_METABOLISM
KEGG_AMINOACYL_TRNA_BIOSYNTHESIS
KEGG_METABOLISM_OF_XENOBIOTICS_BY_CYTOCHROME_P450
KEGG_DRUG_METABOLISM_CYTOCHROME_P450
KEGG_DRUG_METABOLISM_OTHER_ENZYMES
KEGG_BIOSYNTHESIS_OF_UNSATURATED_FATTY_ACIDS
KEGG_ABC_TRANSPORTERS
KEGG_RIBOSOME
KEGG_RNA_DEGRADATION
KEGG_RNA_POLYMERASE
KEGG_BASAL_TRANSCRIPTION_FACTORS
KEGG_DNA_REPLICATION
KEGG_SPLICEOSOME
KEGG_PROTEASOME
KEGG_PROTEIN_EXPORT
KEGG_PPAR_SIGNALING_PATHWAY
KEGG_BASE_EXCISION_REPAIR
KEGG_NUCLEOTIDE_EXCISION_REPAIR
KEGG_MISMATCH_REPAIR
KEGG_HOMOLOGOUS_RECOMBINATION
KEGG_NON_HOMOLOGOUS_END_JOINING
KEGG_MAPK_SIGNALING_PATHWAY
KEGG_ERBB_SIGNALING_PATHWAY
KEGG_CALCIUM_SIGNALING_PATHWAY
KEGG_CYTOKINE_CYTOKINE_RECEPTOR_INTERACTION
KEGG_CHEMOKINE_SIGNALING_PATHWAY
KEGG_PHOSPHATIDYLINOSITOL_SIGNALING_SYSTEM
KEGG_NEUROACTIVE_LIGAND_RECEPTOR_INTERACTION
KEGG_CELL_CYCLE
KEGG_OOCYTE_MEIOSIS
KEGG_P53_SIGNALING_PATHWAY
KEGG_UBIQUITIN_MEDIATED_PROTEOLYSIS
KEGG_SNARE_INTERACTIONS_IN_VESICULAR_TRANSPORT
KEGG_REGULATION_OF_AUTOPHAGY
KEGG_LYSOSOME
KEGG_ENDOCYTOSIS
KEGG_PEROXISOME
KEGG_MTOR_SIGNALING_PATHWAY
KEGG_APOPTOSIS
KEGG_CARDIAC_MUSCLE_CONTRACTION
KEGG_VASCULAR_SMOOTH_MUSCLE_CONTRACTION
KEGG_WNT_SIGNALING_PATHWAY
KEGG_DORSO_VENTRAL_AXIS_FORMATION
KEGG_NOTCH_SIGNALING_PATHWAY
KEGG_HEDGEHOG_SIGNALING_PATHWAY

KEGG_TGF_BETA_SIGNALING_PATHWAY
KEGG_AXON_GUIDANCE
KEGG_VEGF_SIGNALING_PATHWAY
KEGG_FOCAL_ADHESION
KEGG_ECM_RECEPTOR_INTERACTION
KEGG_CELL_ADHESION_MOLECULES_CAMS
KEGG_ADHERENS_JUNCTION
KEGG_TIGHT_JUNCTION
KEGG_GAP_JUNCTION
KEGG_COMPLEMENT_AND_COAGULATION_CASCADES
KEGG_ANTIGEN_PROCESSING_AND_PRESENTATION
KEGG_RENIN_ANGIOTENSIN_SYSTEM
KEGG_TOLL_LIKE_RECEPTOR_SIGNALING_PATHWAY
KEGG_NOD_LIKE_RECEPTOR_SIGNALING_PATHWAY
KEGG_RIG_I_LIKE_RECEPTOR_SIGNALING_PATHWAY
KEGG_CYTOSOLIC_DNA_SENSING_PATHWAY
KEGG_JAK_STAT_SIGNALING_PATHWAY
KEGG_HEMATOPOIETIC_CELL_LINEAGE
KEGG_NATURAL_KILLER_CELL_MEDIATED_CYTOTOXICITY
KEGG_T_CELL_RECEPTOR_SIGNALING_PATHWAY
KEGG_B_CELL_RECEPTOR_SIGNALING_PATHWAY
KEGG_FC_EPSILON_RI_SIGNALING_PATHWAY
KEGG_FC_GAMMA_R_MEDIATED_PHAGOCYTOSIS
KEGG_LEUKOCYTE_TRANSENDOTHELIAL_MIGRATION
KEGG_INSULIN_SIGNALING_PATHWAY
KEGG_GNRH_SIGNALING_PATHWAY
KEGG_NEUROTROPHIN_SIGNALING_PATHWAY
REACTOME_GLYCOGEN_BREAKDOWN_GLYCOGENOLYSIS
REACTOME_TRANSLATION
REACTOME_PYRIMIDINE_CATABOLISM
REACTOME_RNA_POL_III_TRANSCRIPTION_INITIATION_FROM_TYPE_2_PROMOTER
REACTOME_PYRUVATE_METABOLISM_AND_CITRIC_ACID_TCA_CYCLE
REACTOME_PTM_GAMMA_CARBOXYLATION_HYPUSINE_FORMATION_AND_ARYLSULFATASE_ACTIVATION
REACTOME_RNA_POL_I_TRANSCRIPTION_TERMINATION
REACTOME_ACTIVATION_OF_THE_PRE_REPLICATIVE_COMPLEX
REACTOME_PROCESSING_OF_INTRONLESS_PRE_MRNAS
REACTOME_GAP_JUNCTION_DEGRADATION
REACTOME_BILE_ACID_AND_BILE_SALT_METABOLISM
REACTOME_SYNTHESIS_OF_BILE_ACIDS_AND_BILE_SALTS_VIA_7ALPHA_HYDROXYCHOLESTEROL
REACTOME_RECYCLING_OF_BILE_ACIDS_AND_SALTS
REACTOME_METABOLISM_OF_NON_CODING_RNA
REACTOME_SYNTHESIS_OF_BILE_ACIDS_AND_BILE_SALTS_VIA_24_HYDROXYCHOLESTEROL
REACTOME_SYNTHESIS_OF_BILE_ACIDS_AND_BILE_SALTS
REACTOME_METABOLISM_OF_STEROID_HORMONES_AND_VITAMINS_A_AND_D
REACTOME_ANDROGEN_BIOSYNTHESIS
REACTOME_COPI_MEDIATED_TRANSPORT
REACTOME_TCA_CYCLE_AND_RESPIRATORY_ELECTRON_TRANSPORT
REACTOME_GROWTH_HORMONE_RECEPTOR_SIGNALING
REACTOME_CELL_CELL_COMMUNICATION
REACTOME_ABCA_TRANSPORTERS_IN_LIPID_HOMEOSTASIS
REACTOME_ENDOSOMAL_VACUOLAR_PATHWAY
REACTOME_TETRAHYDROBIOPTERIN_BH4_SYNTHESIS_RECYCLING_SALVAGE_AND_REGULATION
REACTOME_ACTIVATED_AMPK_STIMULATES_FATTY_ACID_OXIDATION_IN_MUSCLE
REACTOME_VITAMIN_B5_PANTOTHENATE_METABOLISM
REACTOME_METABOLISM_OF_VITAMINS_AND_COFACTORS
REACTOME_O_LINKED_GLYCOSYLATION_OF_MUCINS
REACTOME_SULFUR_AMINO_ACID_METABOLISM
REACTOME_SPHINGOLIPID_DE_NOVO_BIOSYNTHESIS
REACTOME_TERMINATION_OF_O_GLYCAN_BIOSYNTHESIS

REACTOME_GLYCOSPHINGOLIPID_METABOLISM
REACTOME_PPARA_ACTIVATES_GENE_EXPRESSION
REACTOME_TRIGLYCERIDE_BIOSYNTHESIS
REACTOME_ACYL_CHAIN_REMODELLING_OF_PI
REACTOME_TGF_BETA_RECEPTOR_SIGNALING_IN_EMT_EPITHELIAL_TO_MESENCHYMAL_TRANSITION
REACTOME_DOWNREGULATION_OF_TGF_BETA_RECEPTOR_SIGNALING
REACTOME_ACYL_CHAIN_REMODELLING_OF_PC
REACTOME_TGF_BETA_RECEPTOR_SIGNALING_ACTIVATES_SMADS
REACTOME_PHOSPHOLIPID_METABOLISM
REACTOME_CS_DS_DEGRADATION
REACTOME_SYNTHESIS_OF_PA
REACTOME_OXYGEN_DEPENDENT_PROLINE_HYDROXYLATION_OF_HYPOXIA_INDUCIBLE_FACTOR_ALPHA
REACTOME_SYNTHESIS_OF_PE
REACTOME_CHONDROITIN_SULFATE_BIOSYNTHESIS
REACTOME_HYALURONAN_UPTAKE_AND_DEGRADATION
REACTOME_HYALURONAN_METABOLISM
REACTOME_KERATAN_SULFATE_BIOSYNTHESIS
REACTOME_ALPHA_LINOLENIC_ACID_ALA_METABOLISM
REACTOME_PI_METABOLISM
REACTOME_CHONDROITIN_SULFATE_DERMATAN_SULFATE_METABOLISM
REACTOME_SYNTHESIS_OF_PC
REACTOME_HS_GAG_BIOSYNTHESIS
REACTOME_KERATAN_SULFATE_KERATIN_METABOLISM
REACTOME_KERATAN_SULFATE_DEGRADATION
REACTOME_HEPARAN_SULFATE_HEPARIN_HS_GAG_METABOLISM
REACTOME_GLYCOSAMINOGLYCAN_METABOLISM
REACTOME_ACYL_CHAIN_REMODELLING_OF_PG
REACTOME_ACYL_CHAIN_REMODELLING_OF_PE
REACTOME_ACYL_CHAIN_REMODELLING_OF_PS
REACTOME_GLYCEROPHOSPHOLIPID_BIOSYNTHESIS
REACTOME_PLATELET_ADHESION_TO_EXPOSED_COLLAGEN
REACTOME_REGULATION_OF_PYRUVATE_DEHYDROGENASE_PDH_COMPLEX
REACTOME_METABOLISM_OF_AMINO_ACIDS_AND_DERIVATIVES
REACTOME_RNA_POL_I_TRANSCRIPTION
REACTOME_FATTY_ACYL_COA_BIOSYNTHESIS
REACTOME_INTEGRIN_CELL_SURFACE_INTERACTIONS
REACTOME_REGULATION_OF_ORNITHINE_DECARBOXYLASE_ODC
REACTOME_CYTOCHROME_P450_ARRANGED_BY_SUBSTRATE_TYPE
REACTOME_BASE_FREE_SUGAR_PHOSPHATE_REMOVAL_VIA_THE_SINGLE_NUCLEOTIDE_REPLACEMENT_PATHWAY
REACTOME_HDL_MEDIATED_LIPID_TRANSPORT
REACTOME_RNA_POL_II_TRANSCRIPTION
REACTOME_RNA_POL_III_TRANSCRIPTION
REACTOME_AMINO_ACID_TRANSPORT_ACROSS_THE_PLASMA_MEMBRANE
REACTOME_ENDOGENOUS_STEROLS
REACTOME_GLYCOLYSIS
REACTOME_MITOCHONDRIAL_FATTY_ACID_BETA_OXIDATION
REACTOME_METABOLISM_OF_POLYAMINES
REACTOME_INTEGRATION_OF_ENERGY_METABOLISM
REACTOME_GLUCONEOGENESIS
REACTOME_MITOCHONDRIAL_TRNA_AMINOACYLATION
REACTOME_CYTOSOLIC_TRNA_AMINOACYLATION
REACTOME_ADENYLATE_CYCLASE_ACTIVATING_PATHWAY
REACTOME_ADENYLATE_CYCLASE_INHIBITORY_PATHWAY
REACTOME_PEPTIDE_HORMONE_BIOSYNTHESIS
REACTOME_ACETYLCHOLINE_BINDING_AND_DOWNSTREAM_EVENTS
REACTOME_ABC_FAMILY_PROTEINS_MEDIATED_TRANSPORT
REACTOME_TRNA_AMINOACYLATION
REACTOME_STEROID_HORMONES
REACTOME_AMINE_DERIVED_HORMONES

REACTOME_INSULIN_SYNTHESIS_AND_PROCESSING
REACTOME_GLUCAGON_SIGNALING_IN_METABOLIC_REGULATION
REACTOME_MRNA_PROCESSING
REACTOME_PEROXISOMAL_LIPID_METABOLISM
REACTOME_METABOLISM_OF_NUCLEOTIDES
REACTOME_AMINE_LIGAND_BINDING_RECEPTORS
REACTOME_METABOLISM_OF_PROTEINS
REACTOME_SEROTONIN_RECEPTORS
REACTOME_MRNA_SPLICING
REACTOME_MRNA_SPLICING_MINOR_PATHWAY
REACTOME_3_UTR_MEDIATED_TRANSLATIONAL_REGULATION
REACTOME_PURINE_RIBONUCLEOSIDE_MONOPHOSPHATE_BIOSYNTHESIS
REACTOME_CITRIC_ACID_CYCLE_TCA_CYCLE
REACTOME_REGULATION_OF_INSULIN_SECRETION_BY_GLUCAGON_LIKE_PEPTIDE1
REACTOME_REGULATION_OF_INSULIN_SECRETION
REACTOME_INHIBITION_OF_INSULIN_SECRETION_BY_ADRENALINE_NORADRENALINE
REACTOME_NUCLEOTIDE_LIKE_PURINERGIC_RECEPTORS
REACTOME_ACTIVATION_OF_CHAPERONES_BY_ATF6_ALPHA
REACTOME_EICOSANOID_LIGAND_BINDING_RECEPTORS
REACTOME_UNFOLDED_PROTEIN_RESPONSE
REACTOME_GLUCAGON_TYPE_LIGAND_RECEPTORS
REACTOME_REGULATION_OF_INSULIN_SECRETION_BY_ACETYLCHOLINE
REACTOME_SYNTHESIS_SECRETION_AND_DEACYLATION_OF_GHRELIN
REACTOME_PURINE_SALVAGE
REACTOME_LYSOSOME_VESICLE_BIOGENESIS
REACTOME_TRANSPORT_OF_GLUCOSE_AND_OTHER_SUGARS_BILE_SALTS_AND_ORGANIC_ACIDS_METAL_IONS_AND_AMINE_COMPOUNDS
REACTOME_SPHINGOLIPID_METABOLISM
REACTOME_CELL_CELL_JUNCTION_ORGANIZATION
REACTOME_TRANSPORT_OF_INORGANIC_CATIONS_ANIONS_AND_AMINO_ACIDS_OLIGOPEPTIDES
REACTOME_GOLGI_ASSOCIATED_VESICLE_BIOGENESIS
REACTOME_AMINO_ACID_AND_OLIGOPEPTIDE_SLC_TRANSPORTERS
REACTOME_BRANCHED_CHAIN_AMINO_ACID_CATABOLISM
REACTOME_SYNTHESIS_OF_DNA
REACTOME_DEADENYLATION_OF_MRNA
REACTOME_MRNA_DECAY_BY_5_TO_3_EXORIBONUCLEASE
REACTOME_ZINC_TRANSPORTERS
REACTOME_METAL_ION_SLC_TRANSPORTERS
REACTOME_AUTODEGRADATION_OF_THE_E3_UBIQUITIN_LIGASE_COP1
REACTOME_METABOLISM_OF_MRNA
REACTOME_MRNA_DECAY_BY_3_TO_5_EXORIBONUCLEASE
REACTOME_BILE_SALT_AND_ORGANIC_ANION_SLC_TRANSPORTERS
REACTOME_DEADENYLATION_DEPENDENT_MRNA_DECAY
REACTOME_AMINE_COMPOUND_SLC_TRANSPORTERS
REACTOME_PYRUVATE_METABOLISM
REACTOME_PURINE_CATABOLISM
REACTOME_GLUCOSE_TRANSPORT
REACTOME_METABOLISM_OF_RNA
REACTOME_MITOTIC_G1_G1_S_PHASES
REACTOME_MYOGENESIS
REACTOME_PHOSPHOLIPASE_C_MEDIATED_CASCADE
REACTOME_ACTIVATION_OF_KAINATE_RECEPTORS_UPON_GLUTAMATE_BINDING
REACTOME_SYNTHESIS_AND_INTERCONVERSION_OF_NUCLEOTIDE_DI_AND_TRIPHOSPHATES
REACTOME_RNA_POL_I_RNA_POL_III_AND_MITOCHONDRIAL_TRANSCRIPTION
REACTOME_DNA_REPAIR
REACTOME_EFFECTS_OF_PIP2_HYDROLYSIS
REACTOME_METABOLISM_OF_LIPIDS_AND_LIPOPROTEINS
REACTOME_FATTY_ACID_TRIACYLGLYCEROL_AND_KETONE_BODY_METABOLISM
REACTOME_TRANSPORT_OF_VITAMINS_NUCLEOSIDES_AND_RELATED_MOLECULES
REACTOME_HIGHLY_CALCIUM_PERMEABLE_POSTSYNAPTIC_NICOTINIC_ACETYLCHOLINE_RECEPTORS

REACTOME_ORGANIC_CATION_ANION_ZWITTERION_TRANSPORT
REACTOME_SYNTHESIS_OF_SUBSTRATES_IN_N_GLYCAN_BIOSYTHESIS
REACTOME_RESPIRATORY_ELECTRON_TRANSPORT
REACTOME_ASPARAGINE_N_LINKED_GLYCOSYLATION
REACTOME_BIOSYNTHESIS_OF_THE_N_GLYCAN_PRECURSOR_DOLICHOL_LIPID_LINKED_OLIGOSACCHARIDE_LLO_AND_TRANSFER_TO_A_NASCENT_PROTEIN
REACTOME_AMINO_ACID_SYNTHESIS_AND_INTERCONVERSION_TRANSAMINATION
REACTOME_NITRIC_OXIDE_STIMULATES_GUANYLATE_CYCLASE
REACTOME_N_GLYCAN_TRIMMING_IN_THE_ER_AND_CALNEXIN_CALRETICULIN_CYCLE
REACTOME_PLATELET_SENSITIZATION_BY_LDL
REACTOME_AQUAPORIN_MEDIATED_TRANSPORT
REACTOME_PLATELET_CALCIUM_HOMEOSTASIS
REACTOME_INCRETIN_SYNTHESIS_SECRETION_AND_INACTIVATION
REACTOME_TRANSPORT_OF_ORGANIC_ANIONS
REACTOME_REGULATION_OF_WATER_BALANCE_BY_RENAL_AQUAPORINS
REACTOME_TRANSPORT_TO_THE_GOLGI_AND_SUBSEQUENT_MODIFICATION
REACTOME_IRON_UPTAKE_AND_TRANSPORT
REACTOME_N_GLYCAN_ANTENNAE_ELONGATION
REACTOME_ION_TRANSPORT_BY_P_TYPE_ATPASES
REACTOME_ADVANCED_GLYCOSYLATION_ENDPRODUCT_RECEPTOR_SIGNALING
REACTOME_N_GLYCAN_ANTENNAE_ELONGATION_IN_THE_MEDIAL_TRANS_GOLGI
REACTOME_ION_CHANNEL_TRANSPORT
REACTOME_ETHANOL_OXIDATION
REACTOME_SYNTHESIS_OF_VERY_LONG_CHAIN_FATTY_ACYL_COAS
REACTOME_DNA_REPLICATION
REACTOME_E2F_MEDIATED_REGULATION_OF_DNA_REPLICATION
REACTOME_METABOLISM_OF_CARBOHYDRATES
REACTOME_HORMONE_SENSITIVE_LIPASE_HSL_MEDIATED_TRIACYLGLYCEROL_HYDROLYSIS
REACTOME_PURINE_METABOLISM
REACTOME_LIPID_DIGESTION_MOBILIZATION_AND_TRANSPORT
REACTOME_TRANSPORT_OF_RIBONUCLEOPROTEINS_INTO_THE_HOST_NUCLEUS
REACTOME_RESPIRATORY_ELECTRON_TRANSPORT_ATP_SYNTHESIS_BY_CHEMIOSMOTIC_COUPLING_AND_HEAT_PRODUCTION_BY_UNCOUPLING_PROTEINS
REACTOME_FORMATION_OF_ATP_BY_CHEMIOSMOTIC_COUPLING
REACTOME_GLUCURONIDATION
REACTOME_REGULATION_OF_GLUCOKINASE_BY_GLUCOKINASE_REGULATORY_PROTEIN
REACTOME_LIPOPROTEIN_METABOLISM
REACTOME_CHYLOMICRON_MEDIATED_LIPID_TRANSPORT
REACTOME_SIGNALING_BY_TGF_BETA_RECEPTOR_COMPLEX
REACTOME_GLUTATHIONE_CONJUGATION
REACTOME_GLUCOSE_METABOLISM
REACTOME_VOLTAGE_GATED_POTASSIUM_CHANNELS
REACTOME_POTASSIUM_CHANNELS
REACTOME_NUCLEOTIDE_BINDING_DOMAIN_LEUCINE_RICH_REPEAT_CONTAINING_RECEPTOR_NLR_SIGNALING_PATHWAYS
REACTOME_AMYLOIDS
REACTOME_TELOMERE_MAINTENANCE
REACTOME_TRYPTOPHAN_CATABOLISM
REACTOME_CHOLESTEROL_BIOSYNTHESIS
REACTOME_METABOLISM_OF_PORPHYRINS
REACTOME_DIGESTION_OF_DIETARY_CARBOHYDRATE
REACTOME_SYNTHESIS_OF_GLYCOSYLPHOSPHATIDYLINOSITOL_GPI
REACTOME_PYRIMIDINE_METABOLISM

**Supplementary Table S5. List of percent positive (green) and negative (blue) association rate of the signaling pathways with pyrimidine metabolism along with meta-correlation values and meta-pvalues identified from the activation scores in the TCGA expression profiles of different cancer types.**

| Gene-sets | Positive Rate (%) | Negative Rate (%) | meta-correlation | meta-pvalue |
|---|---|---|---|---|
| DAIRKEE_TERT_TARGETS | 81.3 | 0.0 | 0.57 | 4.2E-93 |
| STEIN_ESRRA_TARGETS | 81.3 | 0.0 | 0.54 | 1.6E-54 |
| CAMP_UP.V1 | 78.1 | 0.0 | 0.58 | 9.5E-63 |
| MTOR_UP.N4.V1 | 75.0 | 0.0 | 0.51 | 4.0E-70 |
| GARCIA_TARGETS_OF_FLI1_AND_DAX1 | 75.0 | 0.0 | 0.51 | 4.4E-112 |
| MOHANKUMAR_HOXA1_TARGETS | 75.0 | 0.0 | 0.49 | 4.5E-80 |
| ROME_INSULIN_TARGETS_IN_MUSCLE | 75.0 | 0.0 | 0.45 | 1.1E-59 |
| PECE_MAMMARY_STEM_CELL | 71.9 | 0.0 | 0.47 | 6.9E-79 |
| LIU_CMYB_TARGETS | 65.6 | 0.0 | 0.41 | 6.8E-42 |
| MOLENAAR_TARGETS_OF_CCND1_AND_CDK4 | 62.5 | 0.0 | 0.43 | 1.5E-49 |
| KRAS.DF.V1 | 62.5 | 0.0 | 0.43 | 2.3E-40 |
| HORIUCHI_WTAP_TARGETS | 59.4 | 0.0 | 0.45 | 5.3E-53 |
| IL15_UP.V1 | 59.4 | 0.0 | 0.40 | 3.6E-42 |
| CROONQUIST_IL6_DEPRIVATION | 56.3 | 0.0 | 0.42 | 1.5E-53 |
| COLLER_MYC_TARGETS | 56.3 | 0.0 | 0.40 | 3.8E-33 |
| SARRIO_EPITHELIAL_MESENCHYMAL_TRANSITION | 56.3 | 0.0 | 0.38 | 2.2E-44 |
| MYC_UP.V1 | 56.3 | 0.0 | 0.37 | 7.5E-24 |
| OXFORD_RALB_TARGETS | 53.1 | 0.0 | 0.38 | 1.3E-37 |
| NEWMAN_ERCC6_TARGETS | 53.1 | 0.0 | 0.36 | 1.4E-37 |
| BAE_BRCA1_TARGETS | 50.0 | 0.0 | 0.36 | 5.1E-27 |
| RADAEVA_RESPONSE_TO_IFNA1 | 50.0 | 0.0 | 0.36 | 1.5E-29 |
| RORIE_TARGETS_OF_EWSR1_FLI1_FUSION | 50.0 | 3.1 | 0.33 | 3.4E-21 |
| PRC2_EED_UP.V1 | 46.9 | 0.0 | 0.38 | 9.9E-28 |
| TANG_SENESCENCE_TP53_TARGETS | 46.9 | 0.0 | 0.36 | 1.2E-31 |
| MACLACHLAN_BRCA1_TARGETS | 46.9 | 0.0 | 0.36 | 3.5E-35 |
| EIF4E | 46.9 | 0.0 | 0.35 | 3.9E-30 |
| KANG_IMMORTALIZED_BY_TERT | 46.9 | 0.0 | 0.33 | 3.9E-21 |
| VETTER_TARGETS_OF_PRKCA_AND_ETS1 | 46.9 | 0.0 | 0.30 | 4.7E-38 |
| ODONNELL_TFRC_TARGETS | 43.8 | 0.0 | 0.37 | 9.6E-38 |
| OLSSON_E2F3_TARGETS | 43.8 | 0.0 | 0.35 | 3.5E-25 |
| SCHUHMACHER_MYC_TARGETS | 43.8 | 0.0 | 0.34 | 1.1E-30 |
| DER_IFN_GAMMA_RESPONSE | 40.6 | 0.0 | 0.35 | 3.0E-30 |
| WANG_CLIM2_TARGETS | 40.6 | 0.0 | 0.31 | 4.2E-17 |
| ODONNELL_TARGETS_OF_MYC_AND_TFRC | 37.5 | 0.0 | 0.34 | 1.5E-26 |
| FUJII_YBX1_TARGETS | 37.5 | 0.0 | 0.33 | 8.3E-42 |
| MARZEC_IL2_SIGNALING | 37.5 | 0.0 | 0.32 | 4.5E-19 |
| NUYTTEN_EZH2_TARGETS | 37.5 | 0.0 | 0.32 | 2.9E-17 |
| DER_IFN_BETA_RESPONSE | 37.5 | 0.0 | 0.32 | 1.3E-21 |
| PDGF_UP.V1 | 37.5 | 0.0 | 0.31 | 2.0E-17 |
| SWEET_KRAS_TARGETS | 37.5 | 0.0 | 0.31 | 8.0E-17 |
| XU_HGF_TARGETS_REPRESSED_BY_AKT1 | 37.5 | 0.0 | 0.30 | 3.1E-18 |
| ITO_PTTG1_TARGETS | 37.5 | 0.0 | 0.29 | 1.2E-17 |
| RAF_UP.V1 | 37.5 | 0.0 | 0.28 | 3.0E-12 |
| SAGIV_CD24_TARGETS | 37.5 | 0.0 | 0.27 | 5.0E-16 |
| HIRSCH_CELLULAR_TRANSFORMATION_SIGNATURE | 37.5 | 0.0 | 0.25 | 1.3E-12 |
| MYC_UP.V1 | 34.4 | 0.0 | 0.32 | 1.4E-15 |
| DER_IFN_ALPHA_RESPONSE | 34.4 | 0.0 | 0.30 | 2.9E-23 |
| RODRIGUES_DCC_TARGETS | 34.4 | 0.0 | 0.29 | 1.7E-24 |

| | | | | |
|---|---|---|---|---|
| PTEN_DN.V1 | 34.4 | 0.0 | 0.25 | 7.2E-14 |
| RUTELLA_RESPONSE_TO_HGF | 34.4 | 3.1 | 0.26 | 8.8E-14 |
| DORSAM_HOXA9_TARGETS | 31.3 | 0.0 | 0.28 | 2.8E-22 |
| PASTURAL_RIZ1_TARGETS | 31.3 | 0.0 | 0.26 | 2.7E-26 |
| BRCA1_DN.V1 | 31.3 | 0.0 | 0.24 | 3.6E-15 |
| PRAMOONJAGO_SOX4_TARGETS | 28.1 | 0.0 | 0.29 | 4.4E-20 |
| PRC2_EZH2_UP.V1 | 28.1 | 0.0 | 0.29 | 4.3E-13 |
| PARENT_MTOR_SIGNALING | 28.1 | 0.0 | 0.29 | 5.7E-17 |
| ALONSO_METASTASIS_EMT | 28.1 | 0.0 | 0.27 | 3.0E-25 |
| GRANDVAUX_IRF3_TARGETS | 28.1 | 0.0 | 0.25 | 1.8E-22 |
| LUCAS_HNF4A_TARGETS | 28.1 | 0.0 | 0.23 | 1.7E-13 |
| ATF2_UP.V1 | 25.0 | 0.0 | 0.26 | 6.9E-17 |
| CYCLIN_D1_UP.V1 | 25.0 | 0.0 | 0.24 | 1.3E-17 |
| SIRNA_EIF4GI | 25.0 | 0.0 | 0.22 | 2.3E-10 |
| SCHRAMM_INHBA_TARGETS | 25.0 | 0.0 | 0.21 | 1.1E-11 |
| JEON_SMAD6_TARGETS | 25.0 | 0.0 | 0.20 | 1.1E-06 |
| YAP1 | 25.0 | 0.0 | 0.19 | 7.3E-08 |
| ONDER_CDH1_TARGETS_2 | 25.0 | 0.0 | 0.13 | 4.4E-04 |
| GU_PDEF_TARGETS | 25.0 | 3.1 | 0.15 | 4.8E-05 |
| BCAT_BILD_ET_AL | 21.9 | 0.0 | 0.22 | 8.2E-18 |
| RUTELLA_RESPONSE_TO_HGF_VS_CSF2RB_AND_IL4 | 21.9 | 0.0 | 0.22 | 5.5E-13 |
| SANA_TNF_SIGNALING | 21.9 | 0.0 | 0.21 | 4.1E-17 |
| PETROVA_PROX1_TARGETS | 21.9 | 0.0 | 0.21 | 7.3E-08 |
| HENDRICKS_SMARCA4_TARGETS | 21.9 | 0.0 | 0.20 | 3.5E-08 |
| PRC2_SUZ12_UP.V1 | 21.9 | 0.0 | 0.18 | 2.0E-06 |
| SMITH_TERT_TARGETS | 21.9 | 0.0 | 0.16 | 3.0E-05 |
| MIYAGAWA_TARGETS_OF_EWSR1_ETS_FUSIONS | 21.9 | 0.0 | 0.13 | 1.3E-03 |
| ZHU_SKIL_TARGETS | 18.8 | 0.0 | 0.27 | 2.2E-13 |
| FEVR_CTNNB1_TARGETS | 18.8 | 0.0 | 0.20 | 2.5E-12 |
| SHIRAISHI_PLZF_TARGETS | 18.8 | 0.0 | 0.19 | 4.4E-10 |
| PRC1_BMI_UP.V1 | 18.8 | 0.0 | 0.19 | 4.2E-10 |
| SENESE_HDAC3_TARGETS | 18.8 | 0.0 | 0.18 | 1.6E-07 |
| IL21_UP.V1 | 18.8 | 0.0 | 0.17 | 1.6E-08 |
| BASSO_CD40_SIGNALING | 18.8 | 0.0 | 0.17 | 3.4E-09 |
| NAGASHIMA_NRG1_SIGNALING | 18.8 | 0.0 | 0.16 | 3.0E-06 |
| JOHNSTONE_PARVB_TARGETS_1 | 15.6 | 0.0 | 0.20 | 2.3E-17 |
| XU_GH1_AUTOCRINE_TARGETS | 15.6 | 0.0 | 0.14 | 1.1E-04 |
| DASU_IL6_SIGNALING_SCAR | 15.6 | 3.1 | 0.15 | 1.3E-05 |
| LIU_TARGETS_OF_VMYB_VS_CMYB | 15.6 | 3.1 | 0.08 | 3.8E-02 |
| HALMOS_CEBPA_TARGETS | 15.6 | 3.1 | 0.07 | 3.7E-02 |
| FORTSCHEGGER_PHF8_TARGETS | 15.6 | 9.4 | 0.12 | 3.3E-03 |
| JOHNSTONE_PARVB_TARGETS_2 | 12.5 | 0.0 | 0.20 | 4.9E-15 |
| ROETH_TERT_TARGETS | 12.5 | 0.0 | 0.19 | 6.7E-12 |
| DASU_IL6_SIGNALING | 12.5 | 0.0 | 0.18 | 9.4E-08 |
| JAZAG_TGFB1_SIGNALING_VIA_SMAD4 | 12.5 | 0.0 | 0.17 | 2.4E-06 |
| ACOSTA_PROLIFERATION_INDEPENDENT_MYC_TARGETS | 12.5 | 0.0 | 0.15 | 1.5E-06 |
| KRASNOSELSKAYA_ILF3_TARGETS | 12.5 | 0.0 | 0.15 | 1.2E-07 |
| KRAS.BREAST_UP.V1 | 12.5 | 0.0 | 0.14 | 2.2E-04 |
| HOXA9_DN.V1 | 12.5 | 0.0 | 0.13 | 7.4E-05 |
| NUYTTEN_NIPP1_TARGETS | 12.5 | 0.0 | 0.08 | 1.9E-02 |
| PIGF_UP.V1 | 12.5 | 0.0 | 0.06 | 2.9E-02 |
| JAZAG_TGFB1_SIGNALING | 12.5 | 3.1 | 0.16 | 2.8E-07 |
| HINATA_NFKB_TARGETS_KERATINOCYTE | 12.5 | 3.1 | 0.12 | 4.7E-04 |
| GOUYER_TATI_TARGETS | 12.5 | 3.1 | 0.05 | 1.2E-01 |
| ONDER_CDH1_TARGETS_1 | 12.5 | 6.3 | 0.07 | 3.8E-02 |

| | | | | |
|---|---|---|---|---|
| LTE2_UP.V1 | 12.5 | 6.3 | 0.05 | 1.6E-01 |
| PTEN_DN.V2 | 9.4 | 0.0 | 0.16 | 1.8E-05 |
| HASINA_NOL7_TARGETS | 9.4 | 0.0 | 0.15 | 9.4E-07 |
| LINDVALL_IMMORTALIZED_BY_TERT | 9.4 | 0.0 | 0.12 | 4.8E-05 |
| KRAS.LUNG.BREAST_UP.V1 | 9.4 | 0.0 | 0.12 | 3.8E-05 |
| SCIAN_INVERSED_TARGETS_OF_TP53_AND_TP73 | 9.4 | 0.0 | 0.12 | 9.0E-05 |
| HOOI_ST7_TARGETS | 9.4 | 0.0 | 0.10 | 3.8E-03 |
| MIKHAYLOVA_OXIDATIVE_STRESS_RESPONSE_VIA_VHL | 9.4 | 0.0 | 0.06 | 5.8E-02 |
| DAVICIONI_TARGETS_OF_PAX_FOXO1_FUSIONS | 9.4 | 3.1 | 0.11 | 5.2E-03 |
| WIERENGA_STAT5A_TARGETS | 9.4 | 3.1 | 0.11 | 6.1E-04 |
| EGFR_UP.V1 | 9.4 | 3.1 | 0.05 | 9.7E-02 |
| CERVERA_SDHB_TARGETS_1 | 9.4 | 6.3 | 0.04 | 1.4E-01 |
| CHARAFE_BREAST_CANCER_LUMINAL_VS_BASAL | 9.4 | 15.6 | -0.06 | 1.6E-01 |
| LIU_IL13_MEMORY_MODEL | 6.3 | 0.0 | 0.15 | 7.4E-10 |
| TSAI_DNAJB4_TARGETS | 6.3 | 0.0 | 0.15 | 8.2E-10 |
| KRAS.600.LUNG.BREAST_UP.V1 | 6.3 | 0.0 | 0.12 | 7.4E-05 |
| SHI_SPARC_TARGETS | 6.3 | 0.0 | 0.11 | 4.2E-05 |
| DOUGLAS_BMI1_TARGETS | 6.3 | 0.0 | 0.08 | 1.8E-03 |
| EBAUER_TARGETS_OF_PAX3_FOXO1_FUSION | 6.3 | 0.0 | 0.06 | 2.6E-02 |
| GRABARCZYK_BCL11B_TARGETS | 6.3 | 0.0 | 0.06 | 4.9E-02 |
| KRAS.LUNG_UP.V1 | 6.3 | 3.1 | 0.09 | 2.0E-04 |
| STK33_NOMO | 6.3 | 3.1 | 0.02 | 3.1E-01 |
| WEINMANN_ADAPTATION_TO_HYPOXIA | 6.3 | 3.1 | -0.01 | 3.7E-01 |
| NGUYEN_NOTCH1_TARGETS | 6.3 | 6.3 | 0.03 | 2.5E-01 |
| SRC_UP.V1 | 6.3 | 6.3 | 0.00 | 4.6E-01 |
| BMI1_DN.V1 | 6.3 | 15.6 | -0.08 | 5.7E-02 |
| MELLMAN_TUT1_TARGETS | 6.3 | 15.6 | -0.12 | 3.1E-03 |
| CHARAFE_BREAST_CANCER_LUMINAL_VS_MESENCHYMAL | 6.3 | 25.0 | -0.11 | 2.5E-02 |
| STK33 | 3.1 | 0.0 | 0.04 | 1.2E-01 |
| MARKS_HDAC_TARGETS | 3.1 | 0.0 | 0.03 | 2.2E-01 |
| RUTELLA_RESPONSE_TO_CSF2RB_AND_IL4 | 3.1 | 0.0 | 0.02 | 3.0E-01 |
| IGARASHI_ATF4_TARGETS | 3.1 | 3.1 | 0.04 | 1.2E-01 |
| LIU_CDX2_TARGETS | 3.1 | 3.1 | -0.05 | 5.3E-02 |
| SANA_RESPONSE_TO_IFNG | 3.1 | 3.1 | -0.06 | 4.3E-02 |
| PHONG_TNF_TARGETS | 3.1 | 6.3 | 0.03 | 1.9E-01 |
| RODRIGUES_NTN1_TARGETS | 3.1 | 6.3 | -0.08 | 9.6E-03 |
| ONDER_CDH1_TARGETS_3 | 3.1 | 9.4 | -0.01 | 4.1E-01 |
| E2F3_UP.V1 | 3.1 | 9.4 | -0.01 | 4.0E-01 |
| BOQUEST_STEM_CELL_CULTURED_VS_FRESH | 3.1 | 9.4 | -0.06 | 4.7E-02 |
| CHUANG_OXIDATIVE_STRESS_RESPONSE | 3.1 | 9.4 | -0.10 | 1.6E-03 |
| KIM_PTEN_TARGETS | 3.1 | 9.4 | -0.10 | 2.2E-03 |
| COULOUARN_TEMPORAL_TGFB1_SIGNATURE | 3.1 | 12.5 | -0.08 | 6.4E-02 |
| OSADA_ASCL1_TARGETS | 3.1 | 12.5 | -0.08 | 1.9E-02 |
| XU_GH1_EXOGENOUS_TARGETS | 3.1 | 12.5 | -0.09 | 5.3E-03 |
| HUANG_FOXA2_TARGETS | 3.1 | 12.5 | -0.11 | 1.7E-03 |
| ROZANOV_MMP14_TARGETS | 3.1 | 15.6 | -0.07 | 7.5E-02 |
| LEF1_UP.V1 | 3.1 | 21.9 | -0.13 | 2.2E-03 |
| PIEPOLI_LGI1_TARGETS | 3.1 | 21.9 | -0.14 | 2.1E-05 |
| HOLLMANN_APOPTOSIS_VIA_CD40 | 3.1 | 21.9 | -0.17 | 1.1E-07 |
| TGFB_UP.V1 | 3.1 | 21.9 | -0.17 | 1.5E-05 |
| GUENTHER_GROWTH_SPHERICAL_VS_ADHERENT | 3.1 | 21.9 | -0.18 | 3.1E-07 |
| MEL18_DN.V1 | 3.1 | 21.9 | -0.21 | 1.6E-06 |
| BMI1_DN_MEL18_DN.V1 | 3.1 | 25.0 | -0.15 | 2.4E-04 |
| NOJIMA_SFRP2_TARGETS | 3.1 | 25.0 | -0.18 | 5.5E-08 |
| MIZUKAMI_HYPOXIA | 0.0 | 0.0 | 0.07 | 1.2E-02 |

| | | | | |
|---|---|---|---|---|
| KANNAN_TP53_TARGETS | 0.0 | 0.0 | 0.05 | 3.0E-02 |
| RELA_DN.V1 | 0.0 | 0.0 | 0.02 | 2.1E-01 |
| KRAS.PROSTATE_UP.V1 | 0.0 | 0.0 | 0.02 | 2.0E-01 |
| HAN_SATB1_TARGETS | 0.0 | 0.0 | 0.01 | 3.5E-01 |
| BUSA_SAM68_TARGETS | 0.0 | 0.0 | 0.01 | 4.0E-01 |
| HOEGERKORP_CD44_TARGETS_TEMPORAL | 0.0 | 0.0 | -0.07 | 6.8E-03 |
| DE_YY1_TARGETS | 0.0 | 3.1 | 0.01 | 3.5E-01 |
| PURBEY_TARGETS_OF_CTBP1_AND_SATB1 | 0.0 | 3.1 | -0.01 | 3.4E-01 |
| CEBALLOS_TARGETS_OF_TP53_AND_MYC | 0.0 | 3.1 | -0.02 | 2.7E-01 |
| STK33_SKM | 0.0 | 3.1 | -0.05 | 8.6E-02 |
| PDGF_ERK_DN.V1 | 0.0 | 3.1 | -0.10 | 3.9E-04 |
| HOEGERKORP_CD44_TARGETS_DIRECT | 0.0 | 6.3 | -0.01 | 4.4E-01 |
| ERB2_UP.V1 | 0.0 | 6.3 | -0.02 | 2.3E-01 |
| AZARE_NEOPLASTIC_TRANSFORMATION_BY_STAT3 | 0.0 | 6.3 | -0.02 | 2.7E-01 |
| SCHURINGA_STAT5A_TARGETS | 0.0 | 6.3 | -0.04 | 1.2E-01 |
| CTIP_DN.V1 | 0.0 | 6.3 | -0.05 | 9.6E-02 |
| XU_HGF_SIGNALING_NOT_VIA_AKT1_48HR | 0.0 | 6.3 | -0.05 | 1.0E-01 |
| P53_DN.V2 | 0.0 | 6.3 | -0.05 | 5.9E-02 |
| MAHAJAN_RESPONSE_TO_IL1A | 0.0 | 6.3 | -0.06 | 1.8E-02 |
| MEK_UP.V1 | 0.0 | 6.3 | -0.09 | 2.8E-03 |
| KOYAMA_SEMA3B_TARGETS | 0.0 | 6.3 | -0.10 | 5.9E-04 |
| CHEN_HOXA5_TARGETS_6HR | 0.0 | 9.4 | 0.00 | 4.6E-01 |
| JIANG_TIP30_TARGETS | 0.0 | 9.4 | -0.02 | 3.3E-01 |
| JAK2_DN.V1 | 0.0 | 9.4 | -0.04 | 1.1E-01 |
| AIYAR_COBRA1_TARGETS | 0.0 | 9.4 | -0.06 | 2.4E-02 |
| KRAS.50_UP.V1 | 0.0 | 9.4 | -0.09 | 2.5E-03 |
| TAVOR_CEBPA_TARGETS | 0.0 | 9.4 | -0.12 | 4.9E-04 |
| WANG_LMO4_TARGETS | 0.0 | 9.4 | -0.12 | 4.5E-05 |
| KRAS.AMP.LUNG_UP.V1 | 0.0 | 9.4 | -0.13 | 3.8E-06 |
| KRAS.600_UP.V1 | 0.0 | 9.4 | -0.14 | 1.1E-05 |
| PURBEY_TARGETS_OF_CTBP1_NOT_SATB1 | 0.0 | 9.4 | -0.14 | 3.7E-07 |
| KRAS.300_UP.V1 | 0.0 | 9.4 | -0.15 | 5.0E-07 |
| LEE_NEURAL_CREST_STEM_CELL | 0.0 | 12.5 | -0.08 | 1.3E-02 |
| JAATINEN_HEMATOPOIETIC_STEM_CELL | 0.0 | 12.5 | -0.09 | 3.7E-04 |
| XU_HGF_TARGETS_INDUCED_BY_AKT1_48HR | 0.0 | 12.5 | -0.12 | 7.2E-05 |
| DAUER_STAT3_TARGETS | 0.0 | 12.5 | -0.16 | 1.5E-07 |
| DUNNE_TARGETS_OF_AML1_MTG8_FUSION | 0.0 | 12.5 | -0.19 | 6.2E-11 |
| ELVIDGE_HIF1A_TARGETS | 0.0 | 15.6 | -0.12 | 4.2E-04 |
| CHOW_RASSF1_TARGETS | 0.0 | 15.6 | -0.15 | 2.5E-06 |
| BEIER_GLIOMA_STEM_CELL | 0.0 | 15.6 | -0.16 | 1.1E-09 |
| ELVIDGE_HIF1A_AND_HIF2A_TARGETS | 0.0 | 15.6 | -0.17 | 3.8E-05 |
| STEIN_ESRRA_TARGETS_RESPONSIVE_TO_ESTROGEN | 0.0 | 15.6 | -0.21 | 1.1E-14 |
| ELVIDGE_HYPOXIA_BY_DMOG | 0.0 | 18.8 | -0.12 | 4.7E-03 |
| ATM_DN.V1 | 0.0 | 18.8 | -0.15 | 3.1E-07 |
| BILBAN_B_CLL_LPL | 0.0 | 18.8 | -0.15 | 2.1E-06 |
| JOHNSTONE_PARVB_TARGETS_3 | 0.0 | 18.8 | -0.17 | 2.3E-08 |
| PEART_HDAC_PROLIFERATION_CLUSTER | 0.0 | 18.8 | -0.19 | 8.0E-07 |
| CHEN_HOXA5_TARGETS_9HR | 0.0 | 18.8 | -0.22 | 8.4E-11 |
| CHARAFE_BREAST_CANCER_BASAL_VS_MESENCHYMAL | 0.0 | 21.9 | -0.16 | 2.8E-06 |
| ALK_DN.V1 | 0.0 | 21.9 | -0.23 | 1.3E-12 |
| VEGF_A_UP.V1 | 0.0 | 25.0 | -0.23 | 1.5E-14 |
| STREICHER_LSM1_TARGETS | 0.0 | 25.0 | -0.23 | 6.7E-07 |
| GARY_CD5_TARGETS | 0.0 | 25.0 | -0.24 | 1.2E-14 |
| LIANG_SILENCED_BY_METHYLATION | 0.0 | 25.0 | -0.24 | 3.3E-12 |
| CREIGHTON_AKT1_SIGNALING_VIA_MTOR | 0.0 | 31.3 | -0.26 | 2.7E-13 |

| | | | | |
|---|---|---|---|---|
| FURUKAWA_DUSP6_TARGETS_PCI35 | 0.0 | 31.3 | -0.27 | 1.0E-19 |
| ELVIDGE_HYPOXIA | 0.0 | 31.3 | -0.29 | 1.8E-11 |
| SENESE_HDAC1_TARGETS | 0.0 | 34.4 | -0.22 | 1.3E-10 |
| CHANG_POU5F1_TARGETS | 0.0 | 34.4 | -0.23 | 2.5E-12 |
| HOELZEL_NF1_TARGETS | 0.0 | 34.4 | -0.28 | 4.9E-23 |
| GOZGIT_ESR1_TARGETS | 0.0 | 34.4 | -0.28 | 8.8E-13 |
| BOQUEST_STEM_CELL | 0.0 | 34.4 | -0.30 | 1.9E-17 |
| DITTMER_PTHLH_TARGETS | 0.0 | 34.4 | -0.31 | 3.4E-17 |
| DELPUECH_FOXO3_TARGETS | 0.0 | 34.4 | -0.32 | 7.1E-20 |
| MAINA_VHL_TARGETS | 0.0 | 37.5 | -0.24 | 1.3E-13 |
| BEGUM_TARGETS_OF_PAX3_FOXO1_FUSION | 0.0 | 37.5 | -0.31 | 1.4E-21 |
| GAL_LEUKEMIC_STEM_CELL | 0.0 | 37.5 | -0.31 | 7.2E-23 |
| WELCSH_BRCA1_TARGETS | 0.0 | 40.6 | -0.23 | 2.5E-08 |
| SENESE_HDAC1_AND_HDAC2_TARGETS | 0.0 | 40.6 | -0.29 | 2.4E-25 |
| SCIBETTA_KDM5B_TARGETS | 0.0 | 40.6 | -0.31 | 3.3E-26 |
| VANTVEER_BREAST_CANCER_ESR1 | 0.0 | 40.6 | -0.33 | 1.1E-10 |
| OXFORD_RALA_AND_RALB_TARGETS | 0.0 | 43.8 | -0.28 | 3.1E-13 |
| SENESE_HDAC2_TARGETS | 0.0 | 43.8 | -0.29 | 1.2E-20 |
| GHO_ATF5_TARGETS | 0.0 | 50.0 | -0.35 | 9.8E-19 |
| KREPPEL_CD99_TARGETS | 0.0 | 53.1 | -0.34 | 2.2E-33 |
| OXFORD_RALA_TARGETS | 0.0 | 59.4 | -0.38 | 4.0E-33 |
| KRAS.KIDNEY_UP.V1 | 0.0 | 62.5 | -0.40 | 2.9E-39 |
| HOEBEKE_LYMPHOID_STEM_CELL | 0.0 | 62.5 | -0.44 | 1.1E-45 |
| SCIAN_CELL_CYCLE_TARGETS_OF_TP53_AND_TP73 | 0.0 | 65.6 | -0.39 | 7.3E-47 |
| KINSEY_TARGETS_OF_EWSR1_FLII_FUSION | 0.0 | 65.6 | -0.43 | 1.1E-63 |
| UDAYAKUMAR_MED1_TARGETS | 0.0 | 65.6 | -0.45 | 7.5E-56 |
| NATSUME_RESPONSE_TO_INTERFERON_BETA | 0.0 | 65.6 | -0.46 | 1.1E-61 |
| LIU_SOX4_TARGETS | 0.0 | 65.6 | -0.48 | 7.7E-49 |
| LU_EZH2_TARGETS | 0.0 | 68.8 | -0.40 | 3.9E-73 |
| LAU_APOPTOSIS_CDKN2A | 0.0 | 68.8 | -0.45 | 5.6E-49 |
| WANG_SMARCE1_TARGETS | 0.0 | 71.9 | -0.51 | 3.2E-50 |
| MANALO_HYPOXIA | 0.0 | 75.0 | -0.50 | 2.8E-68 |
| BASAKI_YBX1_TARGETS | 0.0 | 75.0 | -0.49 | 1.0E-46 |
| RUIZ_TNC_TARGETS | 0.0 | 78.1 | -0.52 | 4.2E-79 |

**Supplementary Table S6. List of percent positive (green) and negative (blue) association rate of the metabolic processes with pyrimidine metabolism along with meta-correlation values and meta-pvalues identified from the activation scores in the TCGA expression profiles of different cancer types.**

| Gene-sets | Positive Rate (%) | Negative Rate (%) | meta-correlation | meta-pvalue |
|---|---|---|---|---|
| KEGG_PYRIMIDINE_METABOLISM | 100.0 | 0.0 | 0.86 | 0E+00 |
| REACTOME_METABOLISM_OF_NUCLEOTIDES | 100.0 | 0.0 | 0.83 | 0E+00 |
| REACTOME_PYRIMIDINE_METABOLISM | 100.0 | 0.0 | 0.79 | 5E-131 |
| REACTOME_SYNTHESIS_AND_INTERCONVERSION_OF_NUCLEOTIDE_DI_AND_TRIPHOSPHATES | 93.8 | 0.0 | 0.72 | 0E+00 |
| REACTOME_REGULATION_OF_ORNITHINE_DECARBOXYLASE_ODC | 90.6 | 0.0 | 0.68 | 1E-161 |
| REACTOME_SYNTHESIS_OF_DNA | 90.6 | 0.0 | 0.67 | 1E-154 |
| KEGG_PROTEASOME | 90.6 | 0.0 | 0.65 | 2E-134 |
| REACTOME_MITOTIC_G1_G1_S_PHASES | 90.6 | 0.0 | 0.62 | 6E-121 |
| REACTOME_SYNTHESIS_OF_SUBSTRATES_IN_N_GLYCAN_BIOSYTHESIS | 90.6 | 0.0 | 0.61 | 5E-161 |
| REACTOME_AUTODEGRADATION_OF_THE_E3_UBIQUITIN_LIGASE_COP1 | 87.5 | 0.0 | 0.66 | 4E-144 |
| KEGG_DNA_REPLICATION | 87.5 | 0.0 | 0.54 | 1E-102 |
| KEGG_GLYOXYLATE_AND_DICARBOXYLATE_METABOLISM | 84.4 | 0.0 | 0.64 | 7E-95 |
| REACTOME_BIOSYNTHESIS_OF_THE_N_GLYCAN_PRECURSOR_DOLICHOL_LIPID_LINKED_OLIGOSACCHARIDE_LLO_AND_TRANSFER_TO_A_NASCENT_PROTEIN | 84.4 | 0.0 | 0.61 | 9E-159 |
| REACTOME_TCA_CYCLE_AND_RESPIRATORY_ELECTRON_TRANSPORT | 84.4 | 0.0 | 0.60 | 4E-84 |
| REACTOME_RESPIRATORY_ELECTRON_TRANSPORT | 84.4 | 0.0 | 0.59 | 6E-91 |
| REACTOME_RESPIRATORY_ELECTRON_TRANSPORT_ATP_SYNTHESIS_BY_CHEMIOSMOTIC_COUPLING_AND_HEAT_PRODUCTION_BY_UNCOUPLING_PROTEINS_ | 84.4 | 0.0 | 0.59 | 1E-91 |
| REACTOME_METABOLISM_OF_AMINO_ACIDS_AND_DERIVATIVES | 84.4 | 0.0 | 0.58 | 1E-72 |
| KEGG_AMINO_SUGAR_AND_NUCLEOTIDE_SUGAR_METABOLISM | 84.4 | 0.0 | 0.55 | 2E-128 |
| REACTOME_CYTOSOLIC_TRNA_AMINOACYLATION | 84.4 | 0.0 | 0.55 | 3E-75 |
| REACTOME_DNA_REPLICATION | 84.4 | 0.0 | 0.54 | 1E-94 |
| REACTOME_TRNA_AMINOACYLATION | 84.4 | 0.0 | 0.54 | 2E-80 |
| KEGG_BASE_EXCISION_REPAIR | 81.3 | 0.0 | 0.61 | 3E-128 |
| REACTOME_PURINE_METABOLISM | 81.3 | 0.0 | 0.59 | 3E-94 |
| REACTOME_MRNA_SPLICING_MINOR_PATHWAY | 81.3 | 0.0 | 0.58 | 5E-99 |
| REACTOME_METABOLISM_OF_PROTEINS | 81.3 | 0.0 | 0.58 | 8E-106 |
| REACTOME_FORMATION_OF_ATP_BY_CHEMIOSMOTIC_COUPLING | 81.3 | 0.0 | 0.57 | 5E-93 |
| KEGG_OXIDATIVE_PHOSPHORYLATION | 81.3 | 0.0 | 0.57 | 3E-84 |
| KEGG_SPLICEOSOME | 81.3 | 0.0 | 0.51 | 4E-59 |
| REACTOME_MRNA_SPLICING | 81.3 | 0.0 | 0.51 | 2E-62 |
| KEGG_PURINE_METABOLISM | 78.1 | 0.0 | 0.61 | 3E-108 |
| KEGG_NUCLEOTIDE_EXCISION_REPAIR | 78.1 | 0.0 | 0.57 | 3E-125 |
| REACTOME_MRNA_DECAY_BY_3_TO_5_EXORIBONUCLEASE | 78.1 | 0.0 | 0.54 | 3E-78 |
| REACTOME_DNA_REPAIR | 78.1 | 0.0 | 0.53 | 3E-95 |
| KEGG_RNA_POLYMERASE | 78.1 | 0.0 | 0.52 | 8E-87 |
| KEGG_MISMATCH_REPAIR | 78.1 | 0.0 | 0.52 | 3E-88 |
| REACTOME_METABOLISM_OF_VITAMINS_AND_COFACTORS | 78.1 | 0.0 | 0.52 | 2E-98 |
| REACTOME_CITRIC_ACID_CYCLE_TCA_CYCLE | 78.1 | 0.0 | 0.51 | 6E-66 |
| REACTOME_DEADENYLATION_DEPENDENT_MRNA_DECAY | 78.1 | 0.0 | 0.49 | 4E-58 |
| REACTOME_MRNA_PROCESSING | 78.1 | 0.0 | 0.48 | 4E-64 |
| KEGG_RNA_DEGRADATION | 78.1 | 0.0 | 0.48 | 1E-68 |
| REACTOME_METABOLISM_OF_POLYAMINES | 75.0 | 0.0 | 0.54 | 4E-57 |
| KEGG_PROTEIN_EXPORT | 75.0 | 0.0 | 0.53 | 1E-111 |
| REACTOME_METABOLISM_OF_RNA | 75.0 | 0.0 | 0.53 | 5E-71 |
| REACTOME_METABOLISM_OF_MRNA | 75.0 | 0.0 | 0.51 | 5E-66 |
| KEGG_AMINOACYL_TRNA_BIOSYNTHESIS | 75.0 | 0.0 | 0.50 | 1E-69 |
| REACTOME_RNA_POL_II_TRANSCRIPTION | 75.0 | 0.0 | 0.50 | 2E-64 |
| KEGG_FRUCTOSE_AND_MANNOSE_METABOLISM | 75.0 | 0.0 | 0.49 | 1E-68 |
| REACTOME_UNFOLDED_PROTEIN_RESPONSE | 75.0 | 0.0 | 0.49 | 6E-105 |
| REACTOME_RNA_POL_III_TRANSCRIPTION_INITIATION_FROM_TYPE_2_PROMOTER | 75.0 | 0.0 | 0.49 | 1E-49 |
| REACTOME_GLUCOSE_METABOLISM | 75.0 | 0.0 | 0.48 | 3E-44 |
| KEGG_CITRATE_CYCLE_TCA_CYCLE | 75.0 | 0.0 | 0.47 | 5E-57 |

| | | | | |
|---|---|---|---|---|
| REACTOME_E2F_MEDIATED_REGULATION_OF_DNA_REPLICATION | 71.9 | 0.0 | 0.50 | 3E-83 |
| REACTOME_BASE_FREE_SUGAR_PHOSPHATE_REMOVAL_VIA_THE_SINGLE_NUCLEOTIDE_REPLACEMENT_PATHWAY | 71.9 | 0.0 | 0.48 | 3E-70 |
| KEGG_PEROXISOME | 71.9 | 0.0 | 0.46 | 4E-39 |
| REACTOME_RNA_POL_III_TRANSCRIPTION | 71.9 | 0.0 | 0.45 | 3E-49 |
| REACTOME_PYRIMIDINE_CATABOLISM | 68.8 | 0.0 | 0.49 | 5E-36 |
| REACTOME_PURINE_RIBONUCLEOSIDE_MONOPHOSPHATE_BIOSYNTHESIS | 68.8 | 0.0 | 0.45 | 1E-75 |
| KEGG_PYRUVATE_METABOLISM | 68.8 | 0.0 | 0.44 | 5E-38 |
| REACTOME_METABOLISM_OF_NON_CODING_RNA | 68.8 | 0.0 | 0.44 | 4E-73 |
| REACTOME_PYRUVATE_METABOLISM_AND_CITRIC_ACID_TCA_CYCLE | 68.8 | 0.0 | 0.44 | 1E-41 |
| KEGG_SNARE_INTERACTIONS_IN_VESICULAR_TRANSPORT | 68.8 | 0.0 | 0.39 | 8E-34 |
| REACTOME_GLUCONEOGENESIS | 65.6 | 0.0 | 0.48 | 3E-39 |
| REACTOME_METABOLISM_OF_PORPHYRINS | 65.6 | 0.0 | 0.48 | 1E-41 |
| KEGG_GALACTOSE_METABOLISM | 65.6 | 0.0 | 0.47 | 2E-58 |
| REACTOME_MITOCHONDRIAL_TRNA_AMINOACYLATION | 65.6 | 0.0 | 0.47 | 4E-54 |
| REACTOME_PURINE_SALVAGE | 65.6 | 0.0 | 0.45 | 1E-44 |
| KEGG_HOMOLOGOUS_RECOMBINATION | 65.6 | 0.0 | 0.45 | 2E-51 |
| KEGG_CELL_CYCLE | 65.6 | 0.0 | 0.44 | 4E-51 |
| REACTOME_TRANSLATION | 65.6 | 0.0 | 0.44 | 9E-53 |
| KEGG_PENTOSE_PHOSPHATE_PATHWAY | 62.5 | 0.0 | 0.47 | 1E-70 |
| REACTOME_TELOMERE_MAINTENANCE | 62.5 | 0.0 | 0.42 | 4E-58 |
| REACTOME_GLYCOLYSIS | 62.5 | 0.0 | 0.42 | 3E-35 |
| REACTOME_MRNA_DECAY_BY_5_TO_3_EXORIBONUCLEASE | 62.5 | 0.0 | 0.42 | 7E-46 |
| REACTOME_3_UTR_MEDIATED_TRANSLATIONAL_REGULATION | 62.5 | 0.0 | 0.39 | 3E-40 |
| KEGG_RIBOSOME | 62.5 | 0.0 | 0.39 | 2E-41 |
| REACTOME_ACTIVATION_OF_THE_PRE_REPLICATIVE_COMPLEX | 59.4 | 0.0 | 0.43 | 1E-56 |
| KEGG_ONE_CARBON_POOL_BY_FOLATE | 59.4 | 0.0 | 0.42 | 5E-37 |
| REACTOME_TETRAHYDROBIOPTERIN_BH4_SYNTHESIS_RECYCLING_SALVAGE_AND_REGULATION | 59.4 | 0.0 | 0.42 | 5E-28 |
| REACTOME_LYSOSOME_VESICLE_BIOGENESIS | 59.4 | 0.0 | 0.41 | 3E-45 |
| KEGG_VALINE_LEUCINE_AND_ISOLEUCINE_BIOSYNTHESIS | 59.4 | 0.0 | 0.41 | 2E-35 |
| KEGG_TERPENOID_BACKBONE_BIOSYNTHESIS | 59.4 | 0.0 | 0.40 | 3E-27 |
| KEGG_GLYCOSYLPHOSPHATIDYLINOSITOL_GPI_ANCHOR_BIOSYNTHESIS | 59.4 | 0.0 | 0.39 | 5E-44 |
| REACTOME_PEROXISOMAL_LIPID_METABOLISM | 56.3 | 0.0 | 0.39 | 1E-24 |
| REACTOME_COPI_MEDIATED_TRANSPORT | 56.3 | 0.0 | 0.38 | 4E-39 |
| REACTOME_VITAMIN_B5_PANTOTHENATE_METABOLISM | 53.1 | 0.0 | 0.41 | 2E-36 |
| KEGG_RIBOFLAVIN_METABOLISM | 53.1 | 0.0 | 0.39 | 3E-32 |
| KEGG_GLUTATHIONE_METABOLISM | 53.1 | 0.0 | 0.38 | 2E-23 |
| REACTOME_RNA_POL_I_RNA_POL_III_AND_MITOCHONDRIAL_TRANSCRIPTION | 53.1 | 0.0 | 0.36 | 6E-35 |
| KEGG_UBIQUITIN_MEDIATED_PROTEOLYSIS | 53.1 | 0.0 | 0.36 | 8E-41 |
| KEGG_FOLATE_BIOSYNTHESIS | 50.0 | 0.0 | 0.39 | 1E-28 |
| REACTOME_CHOLESTEROL_BIOSYNTHESIS | 50.0 | 0.0 | 0.37 | 3E-29 |
| REACTOME_GOLGI_ASSOCIATED_VESICLE_BIOGENESIS | 50.0 | 0.0 | 0.35 | 2E-30 |
| KEGG_REGULATION_OF_AUTOPHAGY | 50.0 | 0.0 | 0.35 | 3E-28 |
| REACTOME_ASPARAGINE_N_LINKED_GLYCOSYLATION | 50.0 | 0.0 | 0.34 | 5E-38 |
| REACTOME_PURINE_CATABOLISM | 50.0 | 3.1 | 0.32 | 4E-14 |
| REACTOME_RNA_POL_I_TRANSCRIPTION_TERMINATION | 46.9 | 0.0 | 0.37 | 9E-42 |
| KEGG_P53_SIGNALING_PATHWAY | 46.9 | 0.0 | 0.36 | 3E-23 |
| REACTOME_SULFUR_AMINO_ACID_METABOLISM | 46.9 | 0.0 | 0.35 | 7E-18 |
| REACTOME_SYNTHESIS_OF_GLYCOSYLPHOSPHATIDYLINOSITOL_GPI | 46.9 | 0.0 | 0.35 | 2E-26 |
| REACTOME_N_GLYCAN_TRIMMING_IN_THE_ER_AND_CALNEXIN_CALRETICULIN_CYCLE | 46.9 | 0.0 | 0.33 | 6E-27 |
| KEGG_DRUG_METABOLISM_OTHER_ENZYMES | 43.8 | 0.0 | 0.36 | 2E-15 |
| REACTOME_MITOCHONDRIAL_FATTY_ACID_BETA_OXIDATION | 43.8 | 0.0 | 0.34 | 1E-15 |
| REACTOME_RNA_POL_I_TRANSCRIPTION | 43.8 | 0.0 | 0.32 | 2E-29 |
| KEGG_RIG_I_LIKE_RECEPTOR_SIGNALING_PATHWAY | 43.8 | 0.0 | 0.32 | 6E-31 |
| REACTOME_OXYGEN_DEPENDENT_PROLINE_HYDROXYLATION_OF_HYPOXIA_INDUCIBLE_FACTOR_ALPHA | 43.8 | 0.0 | 0.30 | 2E-21 |
| KEGG_N_GLYCAN_BIOSYNTHESIS | 43.8 | 0.0 | 0.30 | 4E-24 |
| KEGG_LYSOSOME | 40.6 | 0.0 | 0.34 | 5E-36 |
| REACTOME_ACTIVATION_OF_CHAPERONES_BY_ATF6_ALPHA | 40.6 | 0.0 | 0.32 | 1E-26 |
| REACTOME_AMINO_ACID_SYNTHESIS_AND_INTERCONVERSION_TRANSAMINATION | 40.6 | 3.1 | 0.31 | 2E-11 |
| KEGG_NICOTINATE_AND_NICOTINAMIDE_METABOLISM | 37.5 | 0.0 | 0.32 | 9E-16 |

| | | | | |
|---|---|---|---|---|
| KEGG_CYSTEINE_AND_METHIONINE_METABOLISM | 37.5 | 0.0 | 0.31 | 1E-19 |
| KEGG_PANTOTHENATE_AND_COA_BIOSYNTHESIS | 37.5 | 0.0 | 0.29 | 1E-19 |
| KEGG_CYTOSOLIC_DNA_SENSING_PATHWAY | 37.5 | 0.0 | 0.29 | 2E-15 |
| KEGG_VALINE_LEUCINE_AND_ISOLEUCINE_DEGRADATION | 37.5 | 0.0 | 0.29 | 9E-16 |
| KEGG_LIMONENE_AND_PINENE_DEGRADATION | 37.5 | 0.0 | 0.29 | 3E-14 |
| REACTOME_PYRUVATE_METABOLISM | 37.5 | 0.0 | 0.24 | 1E-11 |
| KEGG_OTHER_GLYCAN_DEGRADATION | 37.5 | 3.1 | 0.28 | 9E-22 |
| KEGG_GLYCOLYSIS_GLUCONEOGENESIS | 34.4 | 0.0 | 0.31 | 5E-18 |
| KEGG_SELENOAMINO_ACID_METABOLISM | 34.4 | 0.0 | 0.30 | 2E-20 |
| REACTOME_GLUCOSE_TRANSPORT | 34.4 | 0.0 | 0.29 | 3E-24 |
| KEGG_OOCYTE_MEIOSIS | 34.4 | 0.0 | 0.29 | 3E-21 |
| REACTOME_PROCESSING_OF_INTRONLESS_PRE_MRNAS | 34.4 | 0.0 | 0.29 | 2E-18 |
| KEGG_STEROID_BIOSYNTHESIS | 34.4 | 0.0 | 0.28 | 1E-24 |
| REACTOME_DEADENYLATION_OF_MRNA | 31.3 | 0.0 | 0.31 | 6E-24 |
| KEGG_ARGININE_AND_PROLINE_METABOLISM | 31.3 | 0.0 | 0.28 | 7E-14 |
| REACTOME_AMYLOIDS | 31.3 | 0.0 | 0.27 | 1E-25 |
| KEGG_BIOSYNTHESIS_OF_UNSATURATED_FATTY_ACIDS | 31.3 | 0.0 | 0.27 | 8E-19 |
| KEGG_ALANINE_ASPARTATE_AND_GLUTAMATE_METABOLISM | 31.3 | 3.1 | 0.26 | 2E-07 |
| KEGG_BUTANOATE_METABOLISM | 28.1 | 0.0 | 0.27 | 1E-08 |
| KEGG_NON_HOMOLOGOUS_END_JOINING | 28.1 | 0.0 | 0.27 | 2E-21 |
| REACTOME_TGF_BETA_RECEPTOR_SIGNALING_IN_EMT_EPITHELIAL_TO_MESENCHYMAL_TRANSITION | 28.1 | 0.0 | 0.25 | 2E-17 |
| REACTOME_SYNTHESIS_OF_BILE_ACIDS_AND_BILE_SALTS_VIA_7ALPHA_HYDROXYCHOLESTEROL | 28.1 | 0.0 | 0.21 | 4E-09 |
| REACTOME_TGF_BETA_RECEPTOR_SIGNALING_ACTIVATES_SMADS | 28.1 | 0.0 | 0.20 | 2E-08 |
| REACTOME_DOWNREGULATION_OF_TGF_BETA_RECEPTOR_SIGNALING | 28.1 | 0.0 | 0.18 | 9E-07 |
| KEGG_PENTOSE_AND_GLUCURONATE_INTERCONVERSIONS | 28.1 | 3.1 | 0.24 | 2E-09 |
| REACTOME_BRANCHED_CHAIN_AMINO_ACID_CATABOLISM | 25.0 | 0.0 | 0.26 | 2E-12 |
| REACTOME_ADVANCED_GLYCOSYLATION_ENDPRODUCT_RECEPTOR_SIGNALING | 25.0 | 0.0 | 0.23 | 3E-09 |
| KEGG_SULFUR_METABOLISM | 25.0 | 0.0 | 0.22 | 2E-10 |
| REACTOME_TRIGLYCERIDE_BIOSYNTHESIS | 25.0 | 0.0 | 0.18 | 4E-07 |
| REACTOME_IRON_UPTAKE_AND_TRANSPORT | 21.9 | 0.0 | 0.26 | 1E-18 |
| REACTOME_GLYCOSPHINGOLIPID_METABOLISM | 21.9 | 0.0 | 0.24 | 3E-13 |
| REACTOME_GLUTATHIONE_CONJUGATION | 21.9 | 0.0 | 0.23 | 1E-12 |
| REACTOME_REGULATION_OF_GLUCOKINASE_BY_GLUCOKINASE_REGULATORY_PROTEIN | 21.9 | 0.0 | 0.22 | 1E-13 |
| REACTOME_TRYPTOPHAN_CATABOLISM | 21.9 | 0.0 | 0.21 | 4E-11 |
| REACTOME_PTM_GAMMA_CARBOXYLATION_HYPUSINE_FORMATION_AND_ARYLSULFATASE_ACTIVATION | 21.9 | 0.0 | 0.21 | 2E-10 |
| REACTOME_METABOLISM_OF_LIPIDS_AND_LIPOPROTEINS | 21.9 | 0.0 | 0.20 | 2E-07 |
| KEGG_PROPANOATE_METABOLISM | 21.9 | 0.0 | 0.18 | 6E-08 |
| KEGG_STARCH_AND_SUCROSE_METABOLISM | 21.9 | 0.0 | 0.18 | 5E-06 |
| KEGG_BETA_ALANINE_METABOLISM | 21.9 | 0.0 | 0.17 | 2E-04 |
| REACTOME_FATTY_ACID_TRIACYLGLYCEROL_AND_KETONE_BODY_METABOLISM | 21.9 | 0.0 | 0.16 | 1E-05 |
| REACTOME_TRANSPORT_OF_RIBONUCLEOPROTEINS_INTO_THE_HOST_NUCLEUS | 18.8 | 0.0 | 0.24 | 1E-19 |
| KEGG_LYSINE_DEGRADATION | 18.8 | 0.0 | 0.22 | 2E-14 |
| KEGG_GLYCEROPHOSPHOLIPID_METABOLISM | 18.8 | 0.0 | 0.14 | 2E-04 |
| KEGG_FATTY_ACID_METABOLISM | 18.8 | 3.1 | 0.11 | 1E-02 |
| REACTOME_METABOLISM_OF_CARBOHYDRATES | 15.6 | 0.0 | 0.19 | 4E-10 |
| REACTOME_HYALURONAN_UPTAKE_AND_DEGRADATION | 15.6 | 0.0 | 0.19 | 7E-09 |
| REACTOME_PLATELET_SENSITIZATION_BY_LDL | 15.6 | 0.0 | 0.19 | 6E-10 |
| KEGG_BASAL_TRANSCRIPTION_FACTORS | 15.6 | 0.0 | 0.15 | 2E-06 |
| REACTOME_GLYCEROPHOSPHOLIPID_BIOSYNTHESIS | 15.6 | 0.0 | 0.14 | 2E-04 |
| REACTOME_SPHINGOLIPID_METABOLISM | 15.6 | 3.1 | 0.17 | 5E-06 |
| KEGG_STEROID_HORMONE_BIOSYNTHESIS | 15.6 | 3.1 | 0.08 | 2E-02 |
| REACTOME_ENDOSOMAL_VACUOLAR_PATHWAY | 12.5 | 0.0 | 0.19 | 7E-11 |
| KEGG_ANTIGEN_PROCESSING_AND_PRESENTATION | 12.5 | 0.0 | 0.18 | 3E-10 |
| KEGG_GLYCINE_SERINE_AND_THREONINE_METABOLISM | 12.5 | 0.0 | 0.18 | 2E-04 |
| KEGG_GLYCEROLIPID_METABOLISM | 12.5 | 0.0 | 0.15 | 3E-05 |
| REACTOME_ALPHA_LINOLENIC_ACID_ALA_METABOLISM | 12.5 | 0.0 | 0.15 | 3E-05 |
| REACTOME_CHYLOMICRON_MEDIATED_LIPID_TRANSPORT | 12.5 | 0.0 | 0.13 | 1E-04 |
| KEGG_TRYPTOPHAN_METABOLISM | 12.5 | 0.0 | 0.12 | 3E-03 |
| REACTOME_SYNTHESIS_OF_BILE_ACIDS_AND_BILE_SALTS_VIA_24_HYDROXYCHOLESTEROL | 12.5 | 0.0 | 0.09 | 5E-03 |
| REACTOME_PHOSPHOLIPID_METABOLISM | 12.5 | 0.0 | 0.07 | 3E-02 |

| | | | | |
|---|---|---|---|---|
| KEGG_NOD_LIKE_RECEPTOR_SIGNALING_PATHWAY | 12.5 | 3.1 | 0.08 | 1E-02 |
| KEGG_SPHINGOLIPID_METABOLISM | 12.5 | 3.1 | 0.08 | 3E-02 |
| REACTOME_LIPOPROTEIN_METABOLISM | 12.5 | 3.1 | 0.07 | 6E-02 |
| REACTOME_HDL_MEDIATED_LIPID_TRANSPORT | 12.5 | 3.1 | 0.07 | 6E-02 |
| REACTOME_ACTIVATED_AMPK_STIMULATES_FATTY_ACID_OXIDATION_IN_MUSCLE | 12.5 | 3.1 | 0.03 | 3E-01 |
| REACTOME_ZINC_TRANSPORTERS | 9.4 | 0.0 | 0.16 | 2E-08 |
| REACTOME_ANDROGEN_BIOSYNTHESIS | 9.4 | 0.0 | 0.16 | 1E-09 |
| REACTOME_GLYCOGEN_BREAKDOWN_GLYCOGENOLYSIS | 9.4 | 0.0 | 0.14 | 4E-07 |
| KEGG_PHENYLALANINE_METABOLISM | 9.4 | 0.0 | 0.11 | 3E-03 |
| REACTOME_METAL_ION_SLC_TRANSPORTERS | 9.4 | 0.0 | 0.09 | 8E-04 |
| REACTOME_METABOLISM_OF_STEROID_HORMONES_AND_VITAMINS_A_AND_D | 9.4 | 0.0 | 0.09 | 7E-03 |
| REACTOME_SIGNALING_BY_TGF_BETA_RECEPTOR_COMPLEX | 9.4 | 0.0 | 0.09 | 1E-02 |
| KEGG_NATURAL_KILLER_CELL_MEDIATED_CYTOTOXICITY | 9.4 | 0.0 | 0.08 | 7E-03 |
| KEGG_TOLL_LIKE_RECEPTOR_SIGNALING_PATHWAY | 9.4 | 0.0 | 0.08 | 8E-03 |
| REACTOME_ORGANIC_CATION_ANION_ZWITTERION_TRANSPORT | 9.4 | 0.0 | 0.08 | 2E-02 |
| KEGG_HISTIDINE_METABOLISM | 9.4 | 0.0 | 0.07 | 3E-02 |
| REACTOME_LIPID_DIGESTION_MOBILIZATION_AND_TRANSPORT | 9.4 | 0.0 | 0.07 | 5E-02 |
| KEGG_PPAR_SIGNALING_PATHWAY | 9.4 | 0.0 | 0.05 | 1E-01 |
| REACTOME_BILE_ACID_AND_BILE_SALT_METABOLISM | 9.4 | 0.0 | 0.04 | 1E-01 |
| REACTOME_SYNTHESIS_OF_BILE_ACIDS_AND_BILE_SALTS | 9.4 | 0.0 | 0.03 | 2E-01 |
| REACTOME_GLUCURONIDATION | 9.4 | 3.1 | 0.15 | 1E-05 |
| KEGG_ASCORBATE_AND_ALDARATE_METABOLISM | 9.4 | 3.1 | 0.14 | 6E-05 |
| KEGG_METABOLISM_OF_XENOBIOTICS_BY_CYTOCHROME_P450 | 9.4 | 3.1 | 0.07 | 4E-02 |
| REACTOME_NUCLEOTIDE_BINDING_DOMAIN_LEUCINE_RICH_REPEAT_CONTAINING_RECEPTOR_NLR_SIGNALING_PATHWAYS | 9.4 | 3.1 | 0.06 | 5E-02 |
| KEGG_RETINOL_METABOLISM | 9.4 | 3.1 | 0.02 | 3E-01 |
| KEGG_DRUG_METABOLISM_CYTOCHROME_P450 | 9.4 | 3.1 | 0.01 | 4E-01 |
| KEGG_TYROSINE_METABOLISM | 9.4 | 6.3 | 0.02 | 3E-01 |
| KEGG_ALPHA_LINOLENIC_ACID_METABOLISM | 9.4 | 9.4 | 0.02 | 3E-01 |
| REACTOME_FATTY_ACYL_COA_BIOSYNTHESIS | 6.3 | 0.0 | 0.13 | 4E-07 |
| KEGG_APOPTOSIS | 6.3 | 0.0 | 0.12 | 3E-04 |
| REACTOME_STEROID_HORMONES | 6.3 | 0.0 | 0.09 | 7E-03 |
| REACTOME_GAP_JUNCTION_DEGRADATION | 6.3 | 0.0 | 0.09 | 3E-03 |
| REACTOME_HYALURONAN_METABOLISM | 6.3 | 0.0 | 0.05 | 9E-02 |
| REACTOME_GROWTH_HORMONE_RECEPTOR_SIGNALING | 6.3 | 0.0 | -0.05 | 7E-02 |
| KEGG_GLYCOSAMINOGLYCAN_DEGRADATION | 6.3 | 3.1 | 0.14 | 5E-06 |
| REACTOME_PPARA_ACTIVATES_GENE_EXPRESSION | 6.3 | 3.1 | 0.05 | 8E-02 |
| REACTOME_SYNTHESIS_OF_PA | 6.3 | 3.1 | 0.04 | 1E-01 |
| KEGG_CARDIAC_MUSCLE_CONTRACTION | 6.3 | 3.1 | 0.01 | 4E-01 |
| REACTOME_AMINO_ACID_TRANSPORT_ACROSS_THE_PLASMA_MEMBRANE | 6.3 | 3.1 | -0.04 | 1E-01 |
| KEGG_PRIMARY_BILE_ACID_BIOSYNTHESIS | 6.3 | 6.3 | -0.02 | 3E-01 |
| REACTOME_AMINO_ACID_AND_OLIGOPEPTIDE_SLC_TRANSPORTERS | 6.3 | 6.3 | -0.10 | 8E-03 |
| KEGG_ARACHIDONIC_ACID_METABOLISM | 6.3 | 9.4 | 0.00 | 5E-01 |
| KEGG_LINOLEIC_ACID_METABOLISM | 6.3 | 9.4 | -0.05 | 1E-01 |
| KEGG_ETHER_LIPID_METABOLISM | 6.3 | 12.5 | -0.03 | 3E-01 |
| REACTOME_SYNTHESIS_OF_PE | 3.1 | 0.0 | 0.12 | 1E-05 |
| REACTOME_SYNTHESIS_OF_VERY_LONG_CHAIN_FATTY_ACYL_COAS | 3.1 | 0.0 | 0.06 | 3E-02 |
| REACTOME_PEPTIDE_HORMONE_BIOSYNTHESIS | 3.1 | 0.0 | 0.04 | 7E-02 |
| REACTOME_DIGESTION_OF_DIETARY_CARBOHYDRATE | 3.1 | 0.0 | -0.06 | 3E-02 |
| REACTOME_SYNTHESIS_SECRETION_AND_DEACYLATION_OF_GHRELIN | 3.1 | 3.1 | 0.09 | 1E-03 |
| REACTOME_REGULATION_OF_PYRUVATE_DEHYDROGENASE_PDH_COMPLEX | 3.1 | 3.1 | 0.04 | 1E-01 |
| REACTOME_ACYL_CHAIN_REMODELLING_OF_PG | 3.1 | 3.1 | 0.00 | 5E-01 |
| REACTOME_ACYL_CHAIN_REMODELLING_OF_PS | 3.1 | 3.1 | 0.00 | 5E-01 |
| REACTOME_HORMONE_SENSITIVE_LIPASE_HSL_MEDIATED_TRIACYLGLYCEROL_HYDROLYSIS | 3.1 | 3.1 | 0.00 | 5E-01 |
| REACTOME_BILE_SALT_AND_ORGANIC_ANION_SLC_TRANSPORTERS | 3.1 | 3.1 | -0.02 | 3E-01 |
| REACTOME_SYNTHESIS_OF_PC | 3.1 | 3.1 | -0.05 | 4E-02 |
| KEGG_RENIN_ANGIOTENSIN_SYSTEM | 3.1 | 6.3 | -0.01 | 4E-01 |
| REACTOME_SPHINGOLIPID_DE_NOVO_BIOSYNTHESIS | 3.1 | 6.3 | -0.02 | 4E-01 |
| KEGG_FC_GAMMA_R_MEDIATED_PHAGOCYTOSIS | 3.1 | 6.3 | -0.06 | 6E-02 |

| | | | | |
|---|---|---|---|---|
| REACTOME_TRANSPORT_OF_GLUCOSE_AND_OTHER_SUGARS_BILE_SALTS_AND_ORGANIC_ACIDS_METAL_IONS_AND_AMINE_COMPOUNDS | 3.1 | 6.3 | -0.07 | 4E-02 |
| KEGG_FC_EPSILON_RI_SIGNALING_PATHWAY | 3.1 | 9.4 | -0.04 | 1E-01 |
| KEGG_T_CELL_RECEPTOR_SIGNALING_PATHWAY | 3.1 | 9.4 | -0.06 | 4E-02 |
| KEGG_COMPLEMENT_AND_COAGULATION_CASCADES | 3.1 | 15.6 | -0.05 | 1E-01 |
| KEGG_JAK_STAT_SIGNALING_PATHWAY | 3.1 | 15.6 | -0.09 | 2E-02 |
| REACTOME_HIGHLY_CALCIUM_PERMEABLE_POSTSYNAPTIC_NICOTINIC_ACETYLCHOLINE_RECEPTORS | 3.1 | 15.6 | -0.15 | 2E-06 |
| REACTOME_PLATELET_ADHESION_TO_EXPOSED_COLLAGEN | 3.1 | 15.6 | -0.16 | 2E-04 |
| REACTOME_AMINE_DERIVED_HORMONES | 3.1 | 18.8 | -0.13 | 5E-04 |
| REACTOME_ABCA_TRANSPORTERS_IN_LIPID_HOMEOSTASIS | 3.1 | 18.8 | -0.21 | 1E-09 |
| REACTOME_ETHANOL_OXIDATION | 3.1 | 21.9 | -0.12 | 3E-03 |
| REACTOME_CHONDROITIN_SULFATE_DERMATAN_SULFATE_METABOLISM | 3.1 | 21.9 | -0.16 | 5E-05 |
| REACTOME_ENDOGENOUS_STEROLS | 3.1 | 21.9 | -0.17 | 9E-05 |
| REACTOME_INTEGRIN_CELL_SURFACE_INTERACTIONS | 3.1 | 21.9 | -0.20 | 1E-05 |
| REACTOME_CYTOCHROME_P450_ARRANGED_BY_SUBSTRATE_TYPE | 3.1 | 25.0 | -0.18 | 4E-06 |
| REACTOME_TRANSPORT_OF_INORGANIC_CATIONS_ANIONS_AND_AMINO_ACIDS_OLIGOPEPTIDES | 3.1 | 34.4 | -0.27 | 2E-13 |
| KEGG_PHOSPHATIDYLINOSITOL_SIGNALING_SYSTEM | 3.1 | 34.4 | -0.28 | 2E-11 |
| KEGG_ECM_RECEPTOR_INTERACTION | 3.1 | 37.5 | -0.26 | 4E-08 |
| REACTOME_EFFECTS_OF_PIP2_HYDROLYSIS | 3.1 | 50.0 | -0.32 | 4E-21 |
| KEGG_MAPK_SIGNALING_PATHWAY | 3.1 | 50.0 | -0.33 | 5E-22 |
| REACTOME_TRANSPORT_OF_VITAMINS_NUCLEOSIDES_AND_RELATED_MOLECULES | 0.0 | 0.0 | 0.07 | 1E-02 |
| REACTOME_RECYCLING_OF_BILE_ACIDS_AND_SALTS | 0.0 | 0.0 | 0.06 | 3E-03 |
| KEGG_GLYCOSAMINOGLYCAN_BIOSYNTHESIS_KERATAN_SULFATE | 0.0 | 3.1 | 0.07 | 4E-03 |
| REACTOME_INCRETIN_SYNTHESIS_SECRETION_AND_INACTIVATION | 0.0 | 3.1 | 0.06 | 9E-03 |
| KEGG_NITROGEN_METABOLISM | 0.0 | 3.1 | 0.02 | 3E-01 |
| KEGG_VEGF_SIGNALING_PATHWAY | 0.0 | 3.1 | 0.00 | 5E-01 |
| REACTOME_ACYL_CHAIN_REMODELLING_OF_PC | 0.0 | 3.1 | -0.01 | 4E-01 |
| REACTOME_ACYL_CHAIN_REMODELLING_OF_PI | 0.0 | 3.1 | -0.01 | 4E-01 |
| REACTOME_TRANSPORT_TO_THE_GOLGI_AND_SUBSEQUENT_MODIFICATION | 0.0 | 3.1 | -0.08 | 1E-03 |
| REACTOME_TERMINATION_OF_O_GLYCAN_BIOSYNTHESIS | 0.0 | 3.1 | -0.09 | 4E-04 |
| KEGG_TAURINE_AND_HYPOTAURINE_METABOLISM | 0.0 | 3.1 | -0.14 | 7E-07 |
| REACTOME_ACYL_CHAIN_REMODELLING_OF_PE | 0.0 | 6.3 | -0.01 | 4E-01 |
| KEGG_ENDOCYTOSIS | 0.0 | 6.3 | -0.02 | 3E-01 |
| KEGG_INSULIN_SIGNALING_PATHWAY | 0.0 | 6.3 | -0.06 | 1E-02 |
| REACTOME_TRANSPORT_OF_ORGANIC_ANIONS | 0.0 | 6.3 | -0.07 | 3E-02 |
| KEGG_B_CELL_RECEPTOR_SIGNALING_PATHWAY | 0.0 | 6.3 | -0.07 | 1E-02 |
| KEGG_MTOR_SIGNALING_PATHWAY | 0.0 | 6.3 | -0.09 | 4E-03 |
| PENG_GLUCOSE_DEPRIVATION | 0.0 | 6.3 | -0.12 | 4E-07 |
| KEGG_GLYCOSPHINGOLIPID_BIOSYNTHESIS_LACTO_AND_NEOLACTO_SERIES | 0.0 | 9.4 | -0.03 | 3E-01 |
| KEGG_GLYCOSPHINGOLIPID_BIOSYNTHESIS_GLOBO_SERIES | 0.0 | 9.4 | -0.04 | 2E-01 |
| KEGG_CHEMOKINE_SIGNALING_PATHWAY | 0.0 | 9.4 | -0.06 | 4E-02 |
| KEGG_GLYCOSPHINGOLIPID_BIOSYNTHESIS_GANGLIO_SERIES | 0.0 | 9.4 | -0.06 | 3E-02 |
| REACTOME_CS_DS_DEGRADATION | 0.0 | 9.4 | -0.11 | 2E-04 |
| REACTOME_N_GLYCAN_ANTENNAE_ELONGATION | 0.0 | 9.4 | -0.12 | 2E-05 |
| KEGG_CYTOKINE_CYTOKINE_RECEPTOR_INTERACTION | 0.0 | 12.5 | -0.06 | 5E-02 |
| KEGG_GLYCOSAMINOGLYCAN_BIOSYNTHESIS_CHONDROITIN_SULFATE | 0.0 | 12.5 | -0.08 | 2E-02 |
| KEGG_HEMATOPOIETIC_CELL_LINEAGE | 0.0 | 12.5 | -0.08 | 1E-02 |
| REACTOME_INSULIN_SYNTHESIS_AND_PROCESSING | 0.0 | 12.5 | -0.11 | 2E-04 |
| REACTOME_O_LINKED_GLYCOSYLATION_OF_MUCINS | 0.0 | 12.5 | -0.12 | 5E-04 |
| KEGG_LEUKOCYTE_TRANSENDOTHELIAL_MIGRATION | 0.0 | 12.5 | -0.12 | 4E-04 |
| KEGG_NEUROTROPHIN_SIGNALING_PATHWAY | 0.0 | 12.5 | -0.15 | 3E-06 |
| REACTOME_KERATAN_SULFATE_KERATIN_METABOLISM | 0.0 | 15.6 | -0.11 | 3E-04 |
| KEGG_NOTCH_SIGNALING_PATHWAY | 0.0 | 15.6 | -0.12 | 2E-03 |
| REACTOME_ACETYLCHOLINE_BINDING_AND_DOWNSTREAM_EVENTS | 0.0 | 15.6 | -0.16 | 2E-07 |
| REACTOME_N_GLYCAN_ANTENNAE_ELONGATION_IN_THE_MEDIAL_TRANS_GOLGI | 0.0 | 15.6 | -0.17 | 7E-09 |
| KEGG_O_GLYCAN_BIOSYNTHESIS | 0.0 | 15.6 | -0.18 | 6E-08 |
| KEGG_ERBB_SIGNALING_PATHWAY | 0.0 | 15.6 | -0.22 | 4E-11 |
| REACTOME_KERATAN_SULFATE_BIOSYNTHESIS | 0.0 | 18.8 | -0.16 | 4E-08 |

| | | | | |
|---|---|---|---|---|
| REACTOME_ACTIVATION_OF_KAINATE_RECEPTORS_UPON_GLUTAMATE_BINDING | 0.0 | 18.8 | -0.19 | 2E-11 |
| REACTOME_SEROTONIN_RECEPTORS | 0.0 | 18.8 | -0.20 | 2E-10 |
| REACTOME_AMINE_COMPOUND_SLC_TRANSPORTERS | 0.0 | 18.8 | -0.21 | 1E-16 |
| REACTOME_GLUCAGON_TYPE_LIGAND_RECEPTORS | 0.0 | 18.8 | -0.22 | 1E-10 |
| KEGG_GLYCOSAMINOGLYCAN_BIOSYNTHESIS_HEPARAN_SULFATE | 0.0 | 18.8 | -0.23 | 4E-26 |
| REACTOME_CHONDROITIN_SULFATE_BIOSYNTHESIS | 0.0 | 21.9 | -0.16 | 2E-05 |
| KEGG_INOSITOL_PHOSPHATE_METABOLISM | 0.0 | 21.9 | -0.19 | 1E-07 |
| REACTOME_REGULATION_OF_INSULIN_SECRETION | 0.0 | 21.9 | -0.23 | 4E-13 |
| REACTOME_REGULATION_OF_INSULIN_SECRETION_BY_ACETYLCHOLINE | 0.0 | 21.9 | -0.23 | 5E-14 |
| REACTOME_PI_METABOLISM | 0.0 | 25.0 | -0.18 | 5E-08 |
| REACTOME_GLYCOSAMINOGLYCAN_METABOLISM | 0.0 | 25.0 | -0.20 | 5E-09 |
| REACTOME_INTEGRATION_OF_ENERGY_METABOLISM | 0.0 | 25.0 | -0.20 | 4E-09 |
| REACTOME_CELL_CELL_JUNCTION_ORGANIZATION | 0.0 | 25.0 | -0.21 | 1E-10 |
| KEGG_ADHERENS_JUNCTION | 0.0 | 25.0 | -0.21 | 2E-08 |
| REACTOME_CELL_CELL_COMMUNICATION | 0.0 | 25.0 | -0.21 | 4E-08 |
| REACTOME_HS_GAG_BIOSYNTHESIS | 0.0 | 25.0 | -0.24 | 8E-19 |
| REACTOME_NUCLEOTIDE_LIKE_PURINERGIC_RECEPTORS | 0.0 | 28.1 | -0.18 | 2E-06 |
| KEGG_CELL_ADHESION_MOLECULES_CAMS | 0.0 | 28.1 | -0.19 | 4E-07 |
| REACTOME_KERATAN_SULFATE_DEGRADATION | 0.0 | 28.1 | -0.20 | 2E-09 |
| KEGG_TIGHT_JUNCTION | 0.0 | 28.1 | -0.24 | 2E-14 |
| REACTOME_AQUAPORIN_MEDIATED_TRANSPORT | 0.0 | 28.1 | -0.24 | 3E-17 |
| REACTOME_MYOGENESIS | 0.0 | 28.1 | -0.25 | 9E-19 |
| REACTOME_INHIBITION_OF_INSULIN_SECRETION_BY_ADRENALINE_NORADRENALINE | 0.0 | 28.1 | -0.26 | 3E-18 |
| REACTOME_HEPARAN_SULFATE_HEPARIN_HS_GAG_METABOLISM | 0.0 | 31.3 | -0.23 | 6E-18 |
| REACTOME_REGULATION_OF_INSULIN_SECRETION_BY_GLUCAGON_LIKE_PEPTIDE1 | 0.0 | 31.3 | -0.26 | 9E-13 |
| KEGG_FOCAL_ADHESION | 0.0 | 31.3 | -0.26 | 3E-08 |
| REACTOME_GLUCAGON_SIGNALING_IN_METABOLIC_REGULATION | 0.0 | 34.4 | -0.28 | 9E-18 |
| KEGG_ABC_TRANSPORTERS | 0.0 | 37.5 | -0.27 | 3E-15 |
| REACTOME_EICOSANOID_LIGAND_BINDING_RECEPTORS | 0.0 | 37.5 | -0.29 | 9E-15 |
| KEGG_GNRH_SIGNALING_PATHWAY | 0.0 | 40.6 | -0.29 | 2E-13 |
| REACTOME_REGULATION_OF_WATER_BALANCE_BY_RENAL_AQUAPORINS | 0.0 | 40.6 | -0.30 | 6E-25 |
| KEGG_DORSO_VENTRAL_AXIS_FORMATION | 0.0 | 40.6 | -0.32 | 3E-25 |
| KEGG_TGF_BETA_SIGNALING_PATHWAY | 0.0 | 40.6 | -0.33 | 8E-18 |
| KEGG_GAP_JUNCTION | 0.0 | 43.8 | -0.34 | 1E-21 |
| REACTOME_PLATELET_CALCIUM_HOMEOSTASIS | 0.0 | 46.9 | -0.34 | 6E-28 |
| REACTOME_ABC_FAMILY_PROTEINS_MEDIATED_TRANSPORT | 0.0 | 50.0 | -0.29 | 7E-16 |
| REACTOME_AMINE_LIGAND_BINDING_RECEPTORS | 0.0 | 50.0 | -0.32 | 4E-19 |
| KEGG_WNT_SIGNALING_PATHWAY | 0.0 | 50.0 | -0.40 | 3E-35 |
| KEGG_VASCULAR_SMOOTH_MUSCLE_CONTRACTION | 0.0 | 53.1 | -0.37 | 2E-25 |
| REACTOME_ADENYLATE_CYCLASE_INHIBITORY_PATHWAY | 0.0 | 56.3 | -0.39 | 8E-24 |
| REACTOME_PHOSPHOLIPASE_C_MEDIATED_CASCADE | 0.0 | 59.4 | -0.41 | 2E-28 |
| REACTOME_ION_CHANNEL_TRANSPORT | 0.0 | 59.4 | -0.41 | 9E-55 |
| REACTOME_ADENYLATE_CYCLASE_ACTIVATING_PATHWAY | 0.0 | 59.4 | -0.42 | 5E-25 |
| KEGG_HEDGEHOG_SIGNALING_PATHWAY | 0.0 | 62.5 | -0.43 | 4E-46 |
| KEGG_AXON_GUIDANCE | 0.0 | 62.5 | -0.44 | 6E-46 |
| REACTOME_POTASSIUM_CHANNELS | 0.0 | 62.5 | -0.45 | 2E-50 |
| REACTOME_ION_TRANSPORT_BY_P_TYPE_ATPASES | 0.0 | 65.6 | -0.42 | 5E-50 |
| KEGG_NEUROACTIVE_LIGAND_RECEPTOR_INTERACTION | 0.0 | 65.6 | -0.46 | 1E-41 |
| REACTOME_VOLTAGE_GATED_POTASSIUM_CHANNELS | 0.0 | 65.6 | -0.47 | 6E-48 |
| REACTOME_NITRIC_OXIDE_STIMULATES_GUANYLATE_CYCLASE | 0.0 | 68.8 | -0.42 | 2E-43 |
| KEGG_CALCIUM_SIGNALING_PATHWAY | 0.0 | 68.8 | -0.50 | 8E-51 |
| PENG_LEUCINE_DEPRIVATION | 0.0 | 81.3 | -0.54 | 6E-122 |
| PENG_GLUTAMINE_DEPRIVATION | 0.0 | 87.5 | -0.62 | 3E-115 |

**Supplementary Table S7. List of correlation with p-values between KEGG Pyrimidine or REACTOME Pyrimidine metabolism with the top positively and negatively associated pathways in the TCGA expression profiles of BRCA (N = 1082), COAD (N = 592) and UCEC (N = 527) cancer types.**

| **BRCA** | **KEGG PYRIMIDINE** | | **REACTOME PYRIMIDINE** | |
|---|---|---|---|---|
| **Gene sets** | **Correlation** | **p-value** | **Correlation** | **p-value** |
| PYRIMIDINE | 0.86 | 0E+00 | 0.63 | 3E-119 |
| KEGG PYRIMIDINE | 1.00 | NA | 0.70 | 2E-159 |
| REACTOME PYRIMIDINE | 0.70 | 2E-159 | 1.00 | NA |
| DAIRKEE TERT | 0.66 | 1E-137 | 0.68 | 1E-149 |
| STEIN ESRRA | 0.71 | 3E-164 | 0.55 | 1E-85 |
| CAMP | 0.77 | 4E-217 | 0.48 | 2E-63 |
| MTOR | 0.61 | 6E-110 | 0.59 | 2E-100 |
| GARCIA FLI1 AND DAX1 | 0.74 | 6E-190 | 0.55 | 3E-85 |
| MOHANKUMAR HOXA1 | 0.76 | 2E-200 | 0.42 | 9E-47 |
| ROME INSULIN | 0.56 | 7E-91 | 0.18 | 6E-09 |
| PECE STEM CELL | 0.48 | 4E-63 | 0.56 | 4E-90 |
| LIU CMYB | 0.44 | 6E-52 | 0.35 | 9E-32 |
| SCIAN TP53 AND TP73 | -0.66 | 3E-136 | -0.48 | 1E-63 |
| KINSEY EWSR1-FLI1 | -0.73 | 5E-178 | -0.39 | 2E-40 |
| UDAYAKUMAR MED1 | -0.76 | 1E-205 | -0.40 | 2E-43 |
| NATSUME IFN BETA | -0.55 | 9E-87 | -0.45 | 1E-54 |
| LIU SOX4 | -0.77 | 3E-215 | -0.46 | 5E-57 |
| LU EZH2 | -0.45 | 3E-54 | -0.53 | 2E-78 |
| LAU APOPTOSIS CDKN2A | -0.54 | 3E-84 | -0.43 | 9E-50 |
| WANG SMARCE1 | -0.77 | 8E-217 | -0.58 | 3E-96 |
| MANALO HYPOXIA | -0.85 | 2E-298 | -0.48 | 3E-64 |
| BASAKI YBX1 | -0.73 | 4E-178 | -0.68 | 2E-145 |
| RUIZ TNC | -0.75 | 1E-199 | -0.56 | 7E-91 |

| **COAD** | **KEGG PYRIMIDINE** | | **REACTOME PYRIMIDINE** | |
|---|---|---|---|---|
| **Gene-sets** | **Correlation** | **p-value** | **Correlation** | **p-value** |
| PYRIMIDINE | 0.81 | 6E-140 | 0.65 | 1E-71 |
| KEGG PYRIMIDINE | 1.00 | NA | 0.67 | 2E-79 |
| REACTOME PYRIMIDINE | 0.67 | 2E-79 | 1.00 | NA |
| DAIRKEE TERT | 0.78 | 7E-124 | 0.52 | 4E-42 |
| STEIN ESRRA | 0.67 | 7E-78 | 0.44 | 3E-29 |
| CAMP | 0.78 | 1E-124 | 0.39 | 2E-22 |
| MTOR | 0.64 | 1E-70 | 0.54 | 2E-45 |
| GARCIA FLI1 AND DAX1 | 0.81 | 6E-136 | 0.40 | 2E-24 |
| MOHANKUMAR HOXA1 | 0.76 | 1E-111 | 0.41 | 3E-25 |
| ROME INSULIN | 0.56 | 1E-49 | 0.26 | 7E-11 |
| PECE STEM CELL | 0.65 | 7E-73 | 0.44 | 1E-29 |
| LIU CMYB | 0.40 | 9E-24 | 0.59 | 8E-56 |
| SCIAN TP53 AND TP73 | -0.65 | 1E-72 | -0.28 | 4E-12 |
| KINSEY EWSR1-FLI1 | -0.74 | 1E-103 | -0.34 | 7E-18 |
| UDAYAKUMAR MED1 | -0.80 | 2E-132 | -0.40 | 4E-24 |
| NATSUME IFN BETA | -0.55 | 5E-48 | -0.30 | 2E-13 |
| LIU SOX4 | -0.78 | 1E-122 | -0.43 | 1E-27 |
| LU EZH2 | -0.44 | 1E-29 | -0.41 | 2E-25 |
| LAU APOPTOSIS CDKN2A | -0.66 | 3E-76 | -0.33 | 5E-16 |
| WANG SMARCE1 | -0.80 | 3E-135 | -0.39 | 2E-22 |
| MANALO HYPOXIA | -0.83 | 3E-154 | -0.37 | 8E-21 |
| BASAKI YBX1 | -0.63 | 1E-67 | -0.46 | 4E-32 |
| RUIZ TNC | -0.84 | 1E-159 | -0.46 | 6E-33 |

| UCEC | KEGG PYRIMIDINE | | REACTOME PYRIMIDINE | |
|---|---|---|---|---|
| **Gene sets** | **Correlation** | **p-value** | **Correlation** | **p-value** |
| PYRIMIDINE | 0.74 | 7E-94 | 0.70 | 3E-79 |
| KEGG PYRIMIDINE | 1.00 | NA | 0.72 | 8E-87 |
| REACTOME PYRIMIDINE | 0.72 | 8E-87 | 1.00 | NA |
| DAIRKEE TERT | 0.67 | 1E-69 | 0.53 | 4E-39 |
| STEIN ESRRA | 0.61 | 1E-54 | 0.36 | 1E-17 |
| CAMP | 0.66 | 6E-67 | 0.43 | 3E-25 |
| MTOR | 0.66 | 2E-68 | 0.51 | 1E-35 |
| GARCIA FLI1 AND DAX1 | 0.74 | 6E-93 | 0.45 | 4E-27 |
| MOHANKUMAR HOXA1 | 0.67 | 6E-70 | 0.33 | 8E-15 |
| ROME INSULIN | 0.54 | 5E-42 | 0.25 | 3E-09 |
| PECE STEM CELL | 0.48 | 5E-32 | 0.37 | 5E-19 |
| LIU CMYB | 0.45 | 1E-27 | 0.37 | 8E-19 |
| SCIAN TP53 AND TP73 | -0.49 | 7E-34 | -0.29 | 1E-11 |
| KINSEY EWSR1-FLI1 | -0.64 | 4E-61 | -0.31 | 2E-13 |
| UDAYAKUMAR MED1 | -0.66 | 4E-67 | -0.27 | 2E-10 |
| NATSUME IFN BETA | -0.43 | 4E-25 | -0.23 | 9E-08 |
| LIU SOX4 | -0.75 | 2E-95 | -0.45 | 2E-27 |
| LU EZH2 | -0.34 | 1E-15 | -0.37 | 3E-18 |
| LAU APOPTOSIS CDKN2A | -0.60 | 2E-52 | -0.47 | 3E-30 |
| WANG SMARCE1 | -0.66 | 1E-66 | -0.39 | 3E-20 |
| MANALO HYPOXIA | -0.77 | 7E-106 | -0.42 | 6E-24 |
| BASAKI YBX1 | -0.59 | 3E-50 | -0.39 | 1E-20 |
| RUIZ TNC | -0.74 | 7E-94 | -0.51 | 1E-35 |

**Supplementary Table S8. List of differentialy expressed genes identified from RNA-seq expression profiling of $TS^{High}$ and $TS^{Low}$ CALU-1 cells stably expressing mCherry gene under the control of *TYMS promoter*.**

**Up-regulated Genes**

| | | | | | | | | |
|---|---|---|---|---|---|---|---|---|
| SCARB1 | TST | CTSL | C9orf3 | RAB26 | AC068580.4 | MEGF9 | TENM3 | C11orf86 |
| RAB27B | MTFMT | SVEP1 | GTF2IRD1 | CA12 | POR | AP1S3 | DMTN | VCAN |
| MFF | GPR157 | IER5L | RHOB | LMO7 | PDLIM5 | KRT86 | LGSN | SAMD11 |
| CLIP2 | RAP1GAP2 | RHBDD2 | C1R | SLC29A4 | B4GALT4 | CFD | PPFIBP2 | CLDN2 |
| FSBP | METTL7B | PDCD4 | PLCH1 | BMPER | OSGIN1 | MYRF | ERRFI1 | CEACAM6 |
| TRAF4 | ASNS | C1S | TWIST2 | FZD5 | PIK3CD-AS2 | FA2H | C1QTNF6 | AKR1B10 |
| TP53TG1 | FTCDNL1 | RETREG1 | SAPCD2 | TPD52 | FCHO1 | TFPI2 | CSGALNACT1 | FGG |
| CARMIL1 | CHEK2 | PARVB | DENND3 | SPAG1 | TM4SF1 | DIRAS3 | OPLAH | VTN |
| LHX4 | LINC01503 | CHP1 | COL4A5 | CCNYL1 | IGFBP1 | LINC01137 | FAM167A | NOTCH3 |
| ALDH3B1 | HSPB8 | EFNA4 | AMDHD1 | EPAS1 | RAB31 | KIAA0319 | KAZALD1 | FST |
| PDXK | NEDD9 | TMEM168 | CDKL5 | SLFN11 | KIFC2 | MFSD12 | KRT87P | FGL1 |
| UBAC1 | PXN-AS1 | AZIN1-AS1 | AC009779.2 | AC009118.2 | THEM6 | TMEM184A | SYT12 | PTPRD |
| PPT2 | MDH2 | ANKS6 | PKP3 | DERL1 | DSCC1 | ADAMTS9 | DHRS2 | UGT1A9 |
| RPL30 | LRSAM1 | CELSR2 | SLC16A5 | SLCO4A1 | ABCB6 | C2orf82 | SELENBP1 | MUC5B |
| CD24P4 | MISP | INTS10 | GINS4 | RGS20 | NQO2 | STOM | SH3TC1 | CPLX2 |
| OR7E14P | SUCLG2 | MROH6 | NECAB3 | ACOX2 | LLGL2 | CDA | SLC7A2 | PPARGC1A |
| SGK2 | ESRRA | SPSB2 | RAB15 | CYC1 | GOT1 | CYP2S1 | HMOX1 | AC105999.2 |
| NINJ1 | FNBP1 | LRIG3 | S100A4 | GSTM4 | MFAP3L | RHOBTB1 | GDF15 | DEFB1 |
| KLHDC10 | MTMR11 | AC007240.1 | FDX1 | SFI1 | GNG11 | RAET1G | ALDH3A1 | UGT1A6 |
| EEF1A2 | PHF7 | PGD | NR0B1 | AVPI1 | SLC22A18 | ARRB1 | RAP1GAP | AGR2 |
| ZHX1 | PPP3CC | DECR1 | ZP3 | PBK | IGFBP3 | PLA2G4A | PEG10 | GPX2 |
| FZD7 | UAP1L1 | ESCO2 | LINGO1 | AGPAT2 | FOXC1 | ASB4 | CD55 | S100P |
| GNE | CASP10 | FBXO44 | DUSP16 | IFITM3 | TBC1D2 | SCARA3 | FSTL4 | PDK4 |
| SORT1 | LTBP1 | CBLB | LINC00963 | AK1 | SLC6A9 | NUP50-AS1 | POF1B | FAM83A |
| ADAM15 | NDUFB9 | RELL2 | SPP1 | ACSS2 | SEC14L4 | RHCG | AKR1C2 | FXYD2 |
| ACAT2 | IMPA2 | SLC25A4 | CRAT | CHCHD10 | MAFK | AC008443.5 | DOK4 | FGA |
| E2F2 | ALAS1 | PIR | XPR1 | TESC | PDE3A | CYP1B1 | NME1-NME2 | FGB |
| LAMA5 | INO80C | PFKFB2 | GSR | DLK2 | ABCA2 | FLVCR2 | RNF43 | RTN4RL2 |
| APTR | DHRS13 | MSI2 | BAG1 | SLC23A2 | MREG | AL162411.1 | CDH1 | |
| CNTNAP3B | GPRC5A | DHCR7 | WWP1 | TOB1 | RAB3D | TSPAN13 | ANG | |
| ZNF275 | RN7SK | IDH1 | PTGES2 | BMP8B | PTGR1 | JUP | CYP26B1 | |
| NAA38 | GADD45B | CUX1 | ENDOG | CAVIN2 | PDE8B | KRT8 | RHOV | |
| SNN | P2RX5 | ITPRIP | ZNRF3 | NBEAL2 | AC008738.1 | RPS29P16 | AKR1C3 | |
| SLC27A4 | DAPK1 | EPHB4 | HMG20B | SCD | FAM19A5 | AL139819.1 | DKK1 | |
| PAQR6 | HIP1R | GMDS-AS1 | COL4A6 | SLC8B1 | PMP22 | NRCAM | CFH | |
| HK1 | CYB5A | SERPINB1 | ZNF618 | SMOC1 | GATS | ADSSL1 | CPVL | |
| GSN | NNMT | GJA1 | CRMP1 | ATP1B1 | RHOU | EPB41L1 | HAS2 | |
| LGR4 | LAPTM4B | ENO3 | AC091390.4 | GRAMD1B | CCDC159 | IL20RB | HHIPL2 | |
| IRS2 | AS3MT | LPCAT1 | SH3BP5 | ZNF714 | ELF3 | KRT81 | OAS1 | |
| SGCE | THSD4 | P4HA3 | GGT1 | NEDD4L | CRB3 | KRT18 | GAL3ST1 | |
| TMEM189 | AC090204.1 | PARD3 | CHAC1 | LINC01703 | GLRX | IGFL2-AS1 | CTH | |
| TMEM205 | AC135048.1 | BIRC7 | ABCA7 | PON3 | LRRN2 | ASS1 | SYT1 | |
| HS1BP3 | CYB5D2 | NDUFA8 | NOTCH1 | BCO1 | SPTBN2 | CLU | SLC45A4 | |
| ARSE | ZNF653 | ZFP36L2 | LIFR | LACTB2 | CACNA2D1 | GLDC | PALMD | |
| UBALD2 | SHARPIN | SEMA3B | TPPP3 | RHOBTB2 | AC006077.2 | LINC01322 | MYO15B | |
| TP53I11 | KIF13B | PITX1 | CTSD | GPRIN3 | PCK2 | RNF19A | ALDH1L2 | |
| CXCL16 | SNHG22 | C16orf74 | GPX4 | ARHGAP27 | CCND3 | NDRG2 | NR4A1 | |
| FAM47E-STBD1 | CEBPB | EPB41L4B | LINC02057 | C9orf172 | FAM69B | MGLL | AKR1B15 | |
| HS6ST1 | TRIML2 | DGAT1 | HSD3B7 | AL355075.4 | SGK1 | KLF4 | BCHE | |
| AC016205.1 | OLFML2A | TRIB3 | DPP7 | SRP14-AS1 | SREBF1 | KRT8P3 | AKR1C1 | |
| IDH3A | ZNF532 | PCDH9 | TBC1D31 | DUSP5 | IFI30 | NPR3 | MAP7 | |
| PIM1 | RASSF2 | HSPB1 | VWA1 | CD38 | SUCLG2-AS1 | KRT85 | NID2 | |
| NUDT7 | PON2 | PC | ABLIM1 | CNNM2 | GLP2R | FIBCD1 | MTUS1 | |
| GSTP1 | DNMT3B | EFNA1 | DGCR5 | L1CAM | COMTD1 | UNC13D | INSL4 | |
| TRUB2 | C17orf51 | AC026979.3 | FAM83H | PLXND1 | LINC01433 | IL1R1 | WISP2 | |
| MAPRE2 | JAG1 | JAKMIP3 | ARRDC1 | KCNK1 | SLC29A2 | RAB20 | CA2 | |
| THRB | KANK1 | LY6E | EGFL7 | ZNF385A | NT5M | IGFBP2 | MUC13 | |

### Down-regulated Genes

| | | | | | | | | | |
|---|---|---|---|---|---|---|---|---|---|
| AL136295.6 | AC015674.1 | NUAK1 | TRAM1L1 | DNMBP | MARCH4 | CCDC186 | ANKRD10 | AL050341.2 | SPRED2 |
| EVI2B | TG | SLC12A8 | AXL | AC139795.2 | ZNF776 | HOXB13 | DTX3 | QPCT | SPRY2 |
| AL512598.1 | PLEK2 | MT2A | LFNG | A1BG | GOLGA7B | DNAJB5 | CHAMP1 | PLAU | SETDB2 |
| RNU1-60P | CRISPLD2 | CES3 | CDCP1 | LCOR | AC020915.2 | ZNF550 | SLC26A2 | SNX19 | KIF3C |
| NPY4R2 | LAMA4 | ZNF697 | JPH3 | AC015982.1 | AC027031.2 | C6orf223 | RAB8B | LINC01588 | NPIPP1 |
| ZBED2 | AC008536.1 | A1BG-AS1 | ANXA8L1 | FOXD1 | USP49 | PDGFC | FAM111B | LINC00941 | IGF2BP2 |
| PKIA | KCTD12 | PDGFB | TANC2 | ARNTL2 | STK17A | ZNF331 | PLEKHG4 | CAVIN1 | SAP30BP |
| AC025580.2 | FBXO32 | TTC3P1 | TSPAN5 | PTX3 | USP31 | ERCC2 | LRRC6 | DIAPH3 | DIS3 |
| AL365356.5 | TENM2 | COL6A2 | ZNF585B | CDIP1 | ZNF8 | AC112220.4 | MCAM | ZSCAN30 | NARF |
| MECOM | EFNB2 | RASSF10 | KIRREL3 | SYNJ2 | SUPT3H | TBC1D10A | FNDC3A | IRAK1BP1 | DYRK3 |
| SNAI2 | WDR66 | MOXD1 | CCL2 | FUT8 | LINC01116 | LY6K | TMEM238 | PDZK1IP1 | AC139887.2 |
| AC012065.3 | ABLIM3 | SH2B3 | ETS1 | EBI3 | GLIPR2 | ALX1 | RIN3 | PGS1 | PXN |
| AC006504.3 | CCBE1 | SORCS2 | UGT8 | IRAK2 | SAMD4A | ZNF300 | CHST7 | AP4S1 | C1QTNF1 |
| AC010809.1 | ZNF155 | GALNT9 | ZNF586 | MT1A | MFGE8 | ZEB1 | TTC30B | HIPK2 | INAFM2 |
| IL7 | SDC2 | GFRA1 | GPAM | PADI2 | NDST1 | ADAM22 | ZNF30 | TSPYL4 | NRG1 |
| ADRB2 | ACTL8 | LAMC2 | EPHB2 | ZNF827 | SPINT2 | TNIP1 | C20orf194 | MTND6P4 | FGD6 |
| AFF3 | FBLN1 | FRMD6 | CXCL1 | GLIPR1 | DCUN1D3 | MACROD2 | RCAN1 | LINC00842 | BTBD11 |
| AC013451.2 | FLG-AS1 | CDK14 | ATP1A3 | C15orf52 | COG3 | MYBL1 | ESD | NOCT | LINC00707 |
| IGFBP7 | RRAD | ARMCX3 | CRTAC1 | C14orf105 | SP100 | SMS | MCL1 | SDC4 | SLC25A37 |
| CDH4 | HBEGF | PODXL | AK5 | NOP14-AS1 | SLC16A2 | DOPEY2 | REV3L | ANXA6 | TMX3 |
| PDE2A | TLE1P1 | AL138756.1 | SOCS3 | AC010422.3 | INPP4B | UCN2 | ASB16-AS1 | PHLDA1 | PIP4K2B |
| FPR1 | CTHRC1 | CALB2 | ZNF112 | AC002310.1 | ZNF773 | C3 | PEAR1 | MOK | IPO5 |
| KCNQ3 | SNORA73B | VANGL2 | ZNF215 | PCDHGC3 | ZNF674-AS1 | AC006058.1 | UVRAG | MAP3K4 | GNA13 |
| ITGB3 | BCL2A1 | ZNF569 | APBB2 | SP6 | DRAM1 | BIVM-ERCC5 | FAT4 | ELF4 | JRKL |
| PRR16 | FYN | GBP3 | G0S2 | PROSER1 | RBMXL1 | ALPK2 | ZC3H13 | ZFP90 | UXS1 |
| ALOX5AP | CADPS2 | KIAA1549L | IFI27L2 | PID1 | ADAM12 | BIRC3 | ZNF45 | ZNF146 | CASK |
| LOXL1-AS1 | NT5E | PLCXD2 | ZMIZ1-AS1 | PNMA8A | CTSS | TRIM34 | ZNF503 | BCAR3 | CASC15 |
| OXCT1 | MOB3B | CD82 | ANKRD13A | LRP4 | B4GALT6 | PTPN14 | ZNF70 | NXPE3 | C17orf100 |
| LRMDA | TCEAL3 | SLIT3 | CLMP | NUDT11 | ZNF324B | ACO1 | PLEKHO2 | CENPBD1 | HENMT1 |
| BACE1-AS | TINAGL1 | FRMD4A | COL4A2 | AL121748.2 | GNB4 | SLC9A7 | MICAL1 | VPS37B | DOCK1 |
| ADGRL2 | ZNF544 | ZNF185 | ZNF551 | INSIG1 | STRA6 | LRIG1 | ZBTB18 | ZNF460 | PLEKHG3 |
| CORO2B | WNT5A | MEG3 | SRPX | SOX9 | DBP | SPEG | G3BP1 | CLEC2D | MYO10 |
| C5orf17 | RTL8B | SLFN5 | MCOLN2 | PMEPA1 | AC135050.6 | CCDC33 | ZNF180 | HEG1 | RAB11FIP2 |
| IGFN1 | SPOCK1 | NNT-AS1 | TPM1 | FAM66C | SHB | ZNF260 | AC098614.1 | GAL | VPS36 |
| LINC01515 | IL11 | PCDHAC2 | TMEM158 | HOXC6 | ITGA5 | HAS3 | GNA12 | TRIM21 | PIGM |
| NPTX1 | COL5A1 | ANKRD44 | GXYLT2 | HAPLN3 | SPDL1 | ALCAM | PAM | PTPN1 | KIRREL1 |
| ANPEP | AL354714.2 | AFAP1L2 | AC073896.2 | LSR | SMIM8 | AC115837.1 | MYH9 | ARL4C | WDFY2 |
| SEMA7A | ZNF239 | SERPINE1 | AC010326.4 | CAMK2N2 | HERC5 | FOXO1 | LINC01232 | ZFAND4 | FBN1 |
| WISP1 | TNFRSF9 | ZNF223 | ARAP2 | TNC | CNOT6 | PPP4R1L | FTSJ1 | DLEU1 | NFKB1 |
| COL13A1 | GPR176 | LINC01006 | KIF5C | ZMAT3 | C3orf52 | AC018629.1 | CCDC18 | MYCBP2 | |
| RBP1 | GTF2IP12 | HMGA2 | TIPARP | DMKN | IL32 | ZNF710 | MBNL2 | CRYBB2P1 | |
| LINC02086 | LHFPL6 | SFMBT2 | LINC02274 | DSE | RCBTB2 | KPNA3 | AMIGO2 | STK40 | |
| PADI3 | RUNX2 | TNFAIP3 | ALOXE3 | MLF1 | PDLIM7 | PLK2 | ZNF101 | RPGR | |
| MAN1A1 | BTNL9 | MPP1 | RNF182 | PLEKHA2 | GPALPP1 | ENTPD7 | DNER | CD44 | |
| CNRIP1 | TNRC6C | PIK3CD | TRAF1 | WDR70 | CDC14A | AP003486.1 | AC022431.1 | SLC6A15 | |
| AC090409.1 | MATN3 | CHGB | PRTFDC1 | TRIO | KLF7 | LTBP3 | TMEM132A | TMCC2 | |
| OTUB2 | SOX7 | STK26 | TNFAIP6 | COL6A1 | CD274 | SLC25A21-AS | AC025265.1 | CASC3 | |
| ADAMTS12 | ZNF283 | SHOX2 | ZNF222 | ELK3 | ITGA3 | ZNF561 | AL136164.4 | APLF | |
| AOX1 | LIMA1 | LTBP2 | FAM228B | ADARB1 | JUN | F11R | DZIP1 | TBC1D4 | |
| AC008687.6 | ADPRH | FOXL2 | AL590617.2 | CCND1 | AC008870.2 | GTF2F2 | UCHL3 | TMEM246 | |
| HK2 | NNT | CTGF | ARMCX6 | C15orf48 | MT1X | ZNF529 | DDX10 | IRGQ | |
| AC124283.3 | SPHK1 | MAN2B1 | TSPYL5 | EVC2 | C16orf45 | C1GALT1C1L | GGACT | MARC1 | |
| CHRNA9 | CA8 | GSPT2 | BDNF | BX322562.1 | HDGFL3 | PRELID2 | LINC00973 | PFDN1 | |
| RAB3B | ITGBL1 | AC073389.1 | CPQ | RASA3 | AC093616.1 | CYR61 | OCIAD2 | SGPL1 | |
| CSMD3 | SAMD9 | KDELC1 | AC131206.1 | ZNF32 | SOBP | ZNF567 | BBX | GNPTAB | |
| SPOCD1 | GLUD2 | NIPAL4 | SMTN | LRRC8C | RASD2 | SLC6A17 | AC087481.3 | TMTC4 | |
| OSGEPL1-AS1 | ZNF22 | NFE2L3 | CXCL8 | RUNX1 | EHD3 | ARHGEF7 | AHRR | LPIN2 | |
| NKILA | MAGEH1 | FHL2 | LINC00511 | AL512488.1 | ZNF354B | TIMP2 | CACHD1 | FAM3C2 | |
| ADAM19 | MAGEA6 | APOL1 | LYPD1 | ATOX1 | ZNF226 | ZFP36L1 | PEA15 | RRAS2 | |
| AC004585.1 | COL4A1 | TMEM59L | ZNF565 | FHDC1 | TCF7L1 | USP40 | MRAS | AATF | |
| MAP3K7CL | MYZAP | STAC | CLTCL1 | ACER2 | CYTH1 | MNS1 | VMP1 | SERTAD2 | |
| IL6 | GSTM3 | PPP1R13L | SMURF2 | ZNF606 | AC124798.1 | EIF3J-AS1 | AC008741.2 | GPX3 | |
| TCEAL8 | SLC10A4 | CLGN | LRRC75A | ITGAX | NFKBIZ | CYLD | SH3KBP1 | KDM4B | |
| CSF2 | PORCN | LRRFIP1P1 | NAV3 | ITGA2 | ZNF585A | RNF2 | ZMIZ1 | WBP4 | |
| AL096865.1 | FHOD3 | XDH | TOB2P1 | CATSPER1 | BACH1 | LINC00847 | SMAD3 | CHFR | |
| AC243773.2 | ZNF548 | GBP1 | HRH1 | FAM111A | ZNF230 | OCLN | ALKBH8 | ARHGAP10 | |

**Supplementary Table S9. List of differentialy expressed genes identified from A549 TS knockdown cells compared to pLKO.1 (control)**

**Up-regulated genes**

| | | | | |
|---|---|---|---|---|
| TMEM144 | TCN2 | KIF12 | SCD | LRRC75B |
| P4HA3 | BCL2L2-PABPN1 | AFF1 | AGMAT | GRB7 |
| CHEK2 | MEX3A | PKD1L2 | SLC9A3R1 | FGFR4 |
| TIRAP | PCSK9 | CTH | ARID3A | GALC |
| CXXC5 | ID1 | AC141586.1 | SUMF1 | NUDT11 |
| TUBAL3 | AP000769.1 | CDIP1 | TMEM254 | ZNF641 |
| TDRKH | FGB | ADGRG2 | VAMP1 | ID2 |
| ING3 | AC099336.2 | FGL1 | CRELD1 | SH3BP5 |
| C12orf73 | IQGAP2 | DNMT3B | GSAP | AKAP12 |
| HNRNPUL2-BSCL2 | C1QL4 | SAMD11 | NCR3LG1 | CAMK2N1 |
| ZNF76 | AKR1C2 | HILPDA | FAM174A | ZNF516 |
| GAPLINC | HSPA2 | YPEL3 | CTDSPL | SMPDL3B |
| CA11 | AP002360.1 | SGK1 | CYP27B1 | WDR91 |
| ZNF397 | SELENBP1 | EPS8 | PAM | ANG |
| ST3GAL2 | HCN3 | NPR3 | KIAA1161 | C1orf115 |
| IGF1R | ADM | DUSP16 | FBXO41 | MTRNR2L12 |
| TLN2 | HOXA-AS2 | GCHFR | CLK4 | TWIST2 |
| CRB3 | AMACR | HGD | PON3 | AL138963.3 |
| ACVR2B | RAP1GAP2 | TM4SF4 | SLC30A3 | SYBU |
| METTL12 | ID2-AS1 | SYNPO2 | IL32 | DOK4 |
| NUP50-AS1 | BTN3A2 | CDH1 | FRAS1 | ADCY9 |
| C17orf97 | RAB20 | CRYBG2 | JUP | CLCN5 |
| PAQR7 | GCA | RAB37 | DMXL2 | ZC3HAV1L |
| EPS8L1 | LINC00632 | CPS1 | DKK1 | DZIP1L |
| B4GALNT1 | METTL7B | VTN | HSD17B14 | SYTL4 |
| ZNF485 | CXCL2 | GAL3ST1 | PHLPP1 | SCARA5 |
| FAM234B | MR1 | ANXA13 | HSD3B7 | PTP4A3 |
| RNF123 | SARM1 | LARGE1 | KREMEN1 | ID4 |
| AC007240.1 | HOXA6 | HNF4A | COLCA2 | CCDC18-AS1 |
| EPDR1 | ANXA4 | NBR2 | MMP7 | BAAT |
| SLC12A6 | NDRG1 | ANKS4B | TOR1AIP1 | EXOC6 |
| KIAA1217 | CYP2R1 | SRR | IGFBP3 | ALX1 |
| ZFYVE1 | LIPH | CRYM | SLC51B | ARRB1 |
| CSAD | CLDND2 | PLIN2 | MICB | FAM13A |
| SNAI1 | MAGEA6 | CP | IRAK2 | STAT4 |
| BBS2 | GDAP1 | SLC16A4 | SH3RF2 | AL590617.2 |
| INSIG1 | FGFR1 | GABARAPL | R3HDM2 | AL021707.6 |
| SESN2 | ERRFI1 | ADAMTS9 | AC008429.1 | AL133351.4 |
| USP25 | TMEM175 | SERPINA6 | ZHX2 | C8orf4 |
| KLHL8 | TMEM37 | SLC7A7 | TBX2-AS1 | SRPX2 |
| DNAJB9 | FAM66C | FBXO27 | APLP1 | CACNA1G |
| ALPK1 | SDHAF4 | APOH | PTPRG | BMP4 |
| ADORA2B | SELENOM | CIDEC | LINC-PINT | KIF21B |
| C3orf52 | LINC00265 | DFNA5 | HLA-DMA | APOL2 |
| LGALSL | TPGS1 | CA9 | DAB2 | OLFML3 |
| PBX1 | SULT2B1 | PDK4 | BDKRB2 | SPX |
| EPS8L3 | CGN | FGFR3 | DEFB1 | USH1C |
| TENM3 | SLC2A3 | F5 | KRT19 | UPK1B |
| PDZK1 | TMCC1 | TRIM31 | LIN7A | SNHG25 |
| FBXO2 | VASH2 | SLFN11 | AC105243.1 | |
| FZD8 | VCAN | DCDC2 | TM4SF20 | |
| FOS | PTGS2 | PRODH2 | NTS | |
| CNKSR2 | AC092336.1 | HLA-DMB | NEFL | |

### Down-regulated genes

| | | | |
|---|---|---|---|
| AC078909.2 | C1orf61 | BCRP3 | EFCAB7 |
| RNU6-33P | CDH4 | AC068631.2 | LYPLAL1 |
| AC005498.2 | PHF21B | ZNF853 | ZNF607 |
| SPARC | GOLM1 | FGFBP1 | REXO5 |
| AC010997.5 | ADGRG1 | PIBF1 | CKS2 |
| AP001469.2 | AC097376.2 | FAM72C | AC243960.2 |
| HIST2H2AB | TSC22D1 | DPYSL3 | C9orf85 |
| LINC00479 | BCAR3 | CEND1 | TFAP4 |
| ACAP1 | NNMT | NPM1P24 | SERTAD1 |
| AC009469.1 | AL162411.1 | ZFP28 | CCDC150P1 |
| AC008878.1 | TRIML2 | FOSL1 | FLNB |
| ALOX5AP | ZRANB3 | KIRREL3 | TTF1 |
| PRDM16 | AC016705.2 | AC006058.3 | USP12 |
| AL355512.1 | SLC25A45 | SYT13 | KLF4 |
| AL645608.5 | GAR1 | AC018629.1 | CAMLG |
| SPOCK1 | LINP1 | COL5A1 | MYH9 |
| AL606834.3 | MSRA | FCGBP | NDUFAF2 |
| LINC01978 | TRPM2 | AC017033.1 | RABEP2 |
| LMO1 | SFN | NPY4R | FAM213B |
| CASC8 | AC010969.2 | HOXA7 | AXL |
| TSPAN1 | TIMP2 | CA8 | HID1 |
| AL365356.5 | TBXAS1 | TAF13 | WAC-AS1 |
| PRMT5-AS1 | TAGLN | TNS4 | CDK2AP1 |
| SIK1 | UCHL3 | AC004585.1 | AC092718.4 |
| SNRPGP15 | RRAS2 | AL583722.1 | ADGRE5 |
| CALB2 | CTNNAL1 | ZNF469 | AC005034.3 |
| PYCARD | PLEKHA2 | RASSF10 | C1orf21 |
| HMGA1P8 | INTS13 | AP000347.1 | MPHOSPH6 |
| AC003002.1 | PHF19 | DOK3 | MTSS1L |
| AL445222.1 | RWDD1 | PIN4P1 | AC007485.2 |
| LINC00707 | HPCAL1 | RPS24P8 | PRIMPOL |
| AC034102.1 | SUMO2 | AC004967.1 | AL391056.1 |
| MUC5AC | SNRPE | AL162386.2 | NOP14-AS1 |
| AC145207.5 | AL359183.1 | FAM84B | TYMS |
| LINC00941 | NDUFAF6 | SLC9A7 | LRRC8A |
| LYPD1 | RPL22L1 | LAMC2 | AL031775.1 |
| ARSI | ZNF324B | MT1X | AC012184.3 |
| PDZK1IP1 | RP11-343N15.5 | STMN3 | DUSP7 |
| KCNMA1 | CAVIN1 | EDN2 | SLC25A51 |
| POLN | YAE1D1 | CHRNA10 | ZNF205 |
| NEXN | OTUB2 | AC048341.3 | POLE3 |
| PREX1 | UBE2W | RPS3AP5 | COA6 |
| PHOSPHO2 | ERH | MARCH4 | PTGES3 |
| ZNF185 | EFEMP1 | VIM | PCED1B |
| JCAD | ZNF8 | CRIP2 | |

**Supplementary Table S10. List of differentialy expressed genes identified from the treatment of A549 cell line with BRQ compared to the control.**

**Up-regulated genes**

| | | | | | | | | |
|---|---|---|---|---|---|---|---|---|
| CLCA2 | RTN1 | PDE2A | ADGRV1 | SLCO4C1 | LAMP3 | PLXNB3 | PDZD3 | IQANK1 |
| LCE1F | VSIG1 | CASP1 | CELF2 | MUC20P1 | PADI1 | CRTAC1 | CDS1 | TP53I3 |
| LCE1E | ALOX5 | CPNE9 | ZNF600 | ANTXRL | CGB7 | LUM | DRD4 | ORAI3 |
| LRRC32 | INPP5D | AP003071.5 | AC025164.1 | RHCG | LINC02577 | AMY2B | PCDHB15 | TBC1D2 |
| DRAXIN | MAGEC2 | CEACAM1 | PIK3IP1 | TMEM270 | CLSTN2 | TNFSF9 | CES1 | PCDHGA4 |
| IMPDH1P4 | SYT8 | MAGEB6 | AC023310.4 | MYPN | COL9A2 | PILRA | LDHD | TLR3 |
| SPINK1 | HMCN2 | AC019117.1 | NCF2 | PSG4 | TNFRSF10C | CATSPERG | PAK3 | AC139149.1 |
| SIGLEC14 | AC097478.1 | PDE6G | C1QTNF1-AS1 | GRM1 | KCNJ2-AS1 | AL592494.3 | AC104407.1 | NMNAT2 |
| CFAP74 | SCN2A | VTCN1 | AC099509.1 | AC113383.1 | ZNF540 | PCDHB14 | ZSCAN18 | PAG1 |
| LCE1B | SLAMF7 | FRZB | LINC00475 | ANKFN1 | B3GALT4 | DEPP1 | DUOXA1 | HNMT |
| LDLRAD1 | RTL9 | ABI3BP | TCTEX1D4 | MR1 | KCNA7 | NTF4 | SULT1C2 | FAM198B |
| BLNK | MAF | DQX1 | MUC20 | ODF3L1 | NPHS1 | WISP2 | NID2 | CDH17 |
| UCA1 | CUX2 | TXNIP | DACT1 | CALML3-AS1 | MCC | CYP4F29P | ZNF813 | SYTL2 |
| IGHV3-32 | TMPRSS3 | ABCA13 | KRTAP2-3 | SH3TC2 | SLC51B | MYO1A | TTLL6 | ABHD1 |
| AL157788.1 | HCP5 | NSG1 | PCDHB13 | FRMPD2 | SELENOP | BEX4 | GBP3 | HOXD9 |
| TPRG1LP1 | NBEAP1 | TNFSF14 | AC069360.1 | CCM2L | C4BPA | IGFALS | FLG | CEACAM6 |
| PINCR | ACTBL2 | CYP4F2 | PRCD | AC005865.2 | CD163L1 | KCNJ16 | DNER | ZNF880 |
| HPN | LINC01297 | GUCA2B | BTBD19 | SYK | FAM198B-AS1 | LINC02280 | PTPRU | ZBED2 |
| AL512638.2 | MAFB | ZNF701 | TSPAN1 | BRINP2 | DUX4L50 | FEZ1 | AC245140.3 | GIPR |
| AC105460.1 | ANKUB1 | NR1I2 | CCDC144NL-AS1 | LINC02532 | VIL1 | KRT19 | CFAP70 | ISLR |
| SUN3 | CD177 | AL445487.1 | PCDHB8 | DDIT4L | LCP1 | MPZ | LRRC36 | NKPD1 |
| LINC01204 | RENBP | PIP | UGT2B15 | CCL26 | TSPAN8 | TSPAN10 | MAN1C1 | ZNF554 |
| AC011294.1 | LINC02328 | LY96 | TSPAN11 | LIVAR | AL035446.1 | APOL1 | ESPNL | CALHM5 |
| AC007342.4 | ITIH5 | SPINT1 | ATG9B | NTN1 | LINC01460 | MALRD1 | VTN | FSCN2 |
| AC016717.2 | THBS4 | FAM13C | RAG1 | AC009690.1 | TMEM45B | VWCE | CLTRN | COL20A1 |
| GGT6 | HSD3BP5 | CCDC187 | FLNC | AC005329.1 | RASGRP4 | SLC7A8 | AC007785.1 | AL162586.1 |
| UNC5B-AS1 | TRIM22 | GRIN2C | NOX5 | LINC02015 | SERPINA1 | FER1L6 | AL138781.1 | KCNQ3 |
| HLA-DOA | LINC02541 | CDH3 | AL033527.5 | DYSF | LYPLAL1-AS1 | WDR63 | FBN1 | FAM212B |
| AC105219.2 | IL1B | FETUB | PTPRVP | CD82 | MANSC4 | EPN3 | AC022784.1 | CCDC180 |
| TNFRSF14 | CNTN5 | AC012236.1 | SORBS2 | H19 | MYO7B | S100P | CDKN1A | BBC3 |
| WDFY4 | CD33 | AL391832.3 | ENAM | HDAC9 | AL354714.3 | PCDH1 | KRT16 | ARHGDIB |
| LINC01732 | AC025569.1 | APOBEC3H | AP001803.2 | LINC01091 | EPHA4 | SCG5 | LOXL4 | TRO |
| RGMA | KLHL30 | SERPINB9P1 | MB | LINC01085 | WEE2-AS1 | MYL9 | SHC4 | BAALC-AS1 |
| KRT9 | NPC1L1 | GPNMB | RYR2 | GNAO1 | LRG1 | PHEX | NANOS1 | CHST2 |
| GUCY2EP | PLA1A | SEMA4A | LINC01146 | COL28A1 | SOWAHB | CDC42BPG | AL138828.1 | AC020928.1 |
| DINOL | MAPK15 | SLC5A10 | EGOT | SHC2 | P2RX6 | TMEM86A | PSMB8-AS1 | AP000769.1 |
| LINC02243 | ABCA8 | PTAFR | MMP11 | SERPINA5 | AC010343.3 | LINC01348 | ATP6V1G2 | SPOCK3 |
| LINC01934 | HSH2D | CST2 | STXBP5-AS1 | AC099684.2 | AL355512.1 | GREB1 | TENM4 | UBA7 |
| AC068985.1 | TCN1 | ZNF91 | ABCA12 | AP000229.1 | CSMD3 | AC087752.3 | DNAJC15 | RASAL1 |
| PTCHD4 | SCT | AL354751.1 | TMEM52 | SCIN | CDA | ABCA1 | CALHM3 | NOX1 |
| AC005865.1 | PRUNE2 | CYGB | GRIP2 | ABCG1 | LINC00853 | TP53INP2 | POPDC2 | DHRS9 |
| AC112236.1 | TFEC | PSTPIP2 | ZNF423 | ITGAX | C1orf116 | PLCD4 | AC128689.1 | KITLG |
| FAM84A | FAM49A | AL162582.1 | COL17A1 | TMEM25 | CFHR3 | AL139039.3 | TRIM54 | HIC1 |
| GAST | SLC52A1 | ALPP | NFE2 | GDF6 | AGTR1 | HHAT | SLC16A12 | GABBR2 |
| MACC1 | ATP6V0D2 | CD79B | TMEM255A | GLS2 | RNA5SP175 | DGKA | SLC2A12 | AC107294.2 |
| LHX3 | GPR143 | TP53INP1 | IFITM10 | CAMK2B | CP | ANXA13 | TMEM255B | RASSF4 |
| TREM2 | PLEKHS1 | AC092167.1 | AL133284.1 | AC020571.1 | ACHE | RGL1 | KIAA1324 | TMEM139 |
| RBP3 | PLEKHG7 | LINC01133 | AC090192.2 | KCP | TMCC3 | TINCR | SCART1 | NYNRIN |
| GPR132 | TIMP3 | ABCA4 | LSMEM2 | SEMA3B-AS1 | SPATA17 | AL391832.2 | VSTM2L | PRR15 |
| AC117500.5 | NLRP1 | AC082651.1 | LINC01559 | FOLR1 | WNT7A | LACTB2-AS1 | AP003068.4 | DPF3 |
| CYP11A1 | CYP2E1 | CNR1 | RGS22 | ANKRD1 | POT1-AS1 | AKR1C4 | LINC00322 | RAPGEF4 |
| CCDC178 | AL589182.1 | AC005256.1 | TNFSF15 | LRRC66 | BTG2 | PCDHGA1 | MMP28 | CXCR4 |
| AC005083.1 | AC084026.1 | SNAI2 | TM4SF20 | TMEM236 | KIAA1755 | PATL2 | AC079466.1 | EPPK1 |
| CYP4F23P | GPR87 | ALDH1L1 | PLEKHB1 | PPP2R2B | Dec-01 | F5 | TNXB | CACNA1A |
| SCN4B | C5orf64 | ABAT | LINC00704 | AC009720.1 | ITIH2 | AL137145.2 | IFI16 | ABCC6 |
| NECTIN4 | KY | IL21R | RINL | CTF1 | CDH16 | CPA4 | SYTL5 | ANXA8L1 |
| TMEM176B | TG | PRSS22 | NPY6R | AL135905.1 | ACTA2 | MAP2 | MFSD4A | IFIT2 |
| MLC1 | LINC00589 | SLC22A2 | DIO1 | TRIM31 | NOV | LINC01468 | GPR35 | VSTM4 |
| LINC01832 | ARC | DUOX2 | LINC01895 | RAB37 | AL158206.1 | FRY | FAAH2 | EBF4 |
| COL24A1 | AL356234.2 | APOD | ARHGAP25 | PGM5 | HES2 | SLC2A1-AS1 | AC022028.2 | PCDHB6 |
| PYHIN1 | KISS1 | HDAC2-AS2 | AP000424.1 | PCDHB11 | ACER2 | ANKRD20A11P | MYO16-AS1 | PLA2G4C |
| SUGCT | RNASE1 | LCN2 | SPATA18 | GPR179 | AC026785.3 | LIPG | COLQ | ACVRL1 |
| PTPRR | ACP5 | AC104248.1 | RASGRP3 | CSTA | JPH2 | FLG-AS1 | PAQR6 | JHDM1D-AS1 |

**Up-regulated genes (Continued)**

| | | | | |
|---|---|---|---|---|
| CPE | AC090772.3 | KCNJ2 | IFIT1 | WNT11 |
| RRAD | SMOC1 | FBXO32 | WNT4 | FLRT2 |
| C3 | ADAMTS7 | PCDHA10 | CDH6 | FN1 |
| LINC00482 | SESN1 | GLP2R | HLA-F | AP003119.1 |
| ADGRF4 | ALS2CL | PEAR1 | LINC00638 | GFOD1 |
| ANKRD2 | IGDCC4 | CHRNB4 | DUSP10 | PARP10 |
| AC027682.6 | SRGAP3 | YPEL2 | SAMD9L | LINC02086 |
| AC147651.1 | SCX | AL590004.4 | ROM1 | HSD17B6 |
| TNFAIP8L3 | L1CAM | SUSD4 | WNT9A | REEP2 |
| ALDOC | AC007611.1 | AC104825.1 | CORO2A | LRRC29 |
| IGFL2 | GPER1 | AC006460.1 | HOXC13 | EPHX4 |
| CARD11 | AC103718.1 | ATP2A3 | SERPINB7 | APOE |
| SEMA3B | REPS2 | SERPINE2 | SYNC | CCDC33 |
| SULF2 | PDGFRA | MXD1 | CFLAR-AS1 | CASC15 |
| AC105446.1 | PTP4A3 | TEX41 | B3GALT5 | POC1B-AS1 |
| CHST1 | ANXA8 | EFCAB5 | PAX8-AS1 | CATSPER1 |
| MUC19 | CDH5 | GAS6-AS1 | AC073370.1 | FOXL2NB |
| LAMA4 | HIST1H2BC | CYSRT1 | CDK18 | AL365356.5 |
| CXXC4 | ZNF28 | TMOD1 | AC093827.4 | PFKFB4 |
| ACY3 | ADRB2 | STYK1 | GAPLINC | KRT15 |
| ZNF525 | ARVCF | PHACTR1 | PPP1R14C | JCAD |
| PCAT6 | CACNG6 | DKK3 | RPGRIP1 | BDH2 |
| SPEG | AC007255.1 | PCDHGB4 | AC062017.1 | DENND1C |
| SIDT1 | DPYSL4 | AC010503.4 | U73166.1 | CTSO |
| Sep-04 | TBX6 | PLEKHG1 | RTL5 | LINC01719 |
| ATP6V0A4 | FGF1 | CNFN | TMEM92 | PINK1 |
| MUC3A | CLDN7 | CHRNA9 | C15orf59 | KLHDC7A |
| MUC4 | AZIN2 | CYP3A5 | ZMAT3 | MILR1 |
| EPS8L3 | SPON2 | ZNF561-AS1 | DENND2D | HABP2 |
| PSORS1C1 | BPIFB1 | BAIAP3 | LAMA2 | EFCAB10 |
| CD22 | SCN5A | GAS7 | ABCG4 | LBH |
| NFATC4 | AOC3 | C1S | CSRNP3 | ATF3 |
| TOB1-AS1 | FILIP1L | THY1 | AKAP6 | |
| CRISPLD2 | INSIG1 | NEFL | GPT | |
| DUSP15 | MFGE8 | UNC5B | RUNX2 | |
| ENG | MAFA | PURPL | TAGLN | |
| PALMD | TMEM154 | TMEM105 | LINC00525 | |
| CATIP | SAMD5 | PRRT2 | ZG16B | |
| LINC01759 | COL15A1 | GPR37L1 | STON2 | |
| CYP2W1 | SLC16A8 | AL390726.6 | SMIM2-AS1 | |
| PCDH12 | ASS1 | NAP1L3 | COL13A1 | |
| WDR66 | IGFBP7 | AC002066.1 | OVGP1 | |
| CCNG2 | LINC02331 | POLH-AS1 | SLC22A17 | |
| PLAT | CD70 | HRNR | AC007325.2 | |
| YPEL3 | VWA7 | CORO6 | FAM214B | |
| BCAS1 | AL512353.1 | AC122710.2 | IL7R | |
| AC068580.4 | LGR6 | CDHR1 | USH1C | |
| LCN12 | AC013652.1 | CST1 | EGLN3 | |
| NPAS1 | GOLT1A | GALNT12 | LINC01285 | |
| FADS2P1 | PGF | ICA1 | ITGA4 | |
| ERVMER61-1 | IGFL1P1 | NR1H4 | RAB3B | |
| CD79A | TRIM2 | NECTIN3-AS1 | IFFO1 | |
| NEURL1 | IFIT3 | CHPF | SESN3 | |
| GJB4 | PRR15L | FOXD4L1 | AL645608.8 | |
| AL021154.1 | TLR4 | VANGL2 | LINC00887 | |
| GPX2 | C1R | SMIM10L2A | AL157935.1 | |
| GALNT5 | CTSH | CSDC2 | LINC00886 | |
| ANK1 | FBXO2 | AC005336.1 | GRIN3B | |
| CREBRF | BANK1 | FMO4 | PTPRB | |
| PDZK1IP1 | PCDHGA2 | ECM1 | RARRES3 | |
| AOX1 | ARMCX1 | AC099489.1 | BEX2 | |
| FAM83E | AMOT | EHF | ISG15 | |
| MMP2 | AP002884.1 | PARM1 | F8 | |
| ITGA11 | AL117339.5 | HSD17B1P1 | CCDC9B | |

#### Down-regulated genes

| | | | |
|---|---|---|---|
| SYT6 | GTSE1 | ASPM | CNTF |
| WDR76 | RRM2 | EEF1A1P13 | SHISA3 |
| RPL6P27 | CCNA2 | RPL13AP5 | DLEU2 |
| MTFR2 | TUBA1C | SNHG19 | LY6K |
| CDKN3 | HJURP | COL26A1 | CADPS2 |
| PRC1 | SMG1P7 | ZNF25 | CHL1 |
| AC006329.1 | NCAPH | PRR11 | FSTL5 |
| CHAF1A | BUB1B | EEF1A1 | MIR17HG |
| RIMS4 | TUBBP1 | GNB1L | AL391988.1 |
| CENPI | KIF18B | MDH1B | C6orf52 |
| BCL2L12 | CDC45 | PALM3 | DMC1 |
| E2F2 | KIF4A | EEF1A1P6 | SNHG5 |
| IGSF10 | SKA1 | SYP | AC093484.4 |
| ERMARD | PBK | RPL10P16 | CR392039.3 |
| CDC25A | ERCC6L | CCDC87 | SNORA73B |
| LMNB1 | PLK4 | RPL13A | |
| WDR4 | RPL23 | EEF1A1P12 | |
| MTHFD2 | LINC00641 | AC010761.1 | |
| CENPF | MCM2 | BUB1 | |
| AC015813.5 | MCM7 | SULT2B1 | |
| RAD54L | EME1 | RPLP0P6 | |
| TTK | UHRF1 | FAM72A | |
| CENPE | RPLP0 | DLGAP5 | |
| AC091167.1 | HMGB1P10 | SNHG12 | |
| CPLX2 | CDCA2 | KLRA1P | |
| SPC24 | XRCC2 | MCM5 | |
| CDC25C | EIF3CL | KIF15 | |
| DIAPH3 | KIFC1 | NTS | |
| NCAPG | AL645608.1 | HNRNPA1P10 | |
| MCM6 | FBXO4 | FAM72B | |
| PIMREG | RPL10 | FBXO36 | |
| KIF2C | AC002116.2 | SNHG25 | |
| SKA3 | NCAPG2 | KCNC3 | |
| AURKB | DEPDC1 | AC026401.3 | |
| RPS10 | PCDH17 | SKA2 | |
| PRIM1 | SAPCD2 | RPL13AP25 | |
| CIP2A | KIF11 | SNHG4 | |
| SGO1 | EEF1A1P19 | CDC20P1 | |
| VPS9D1-AS1 | TEX15 | LEPR | |
| UBE2S | NPTX1 | PLK1 | |
| SPAG5 | E2F8 | POU3F3 | |
| SPC25 | RPL10P9 | AC090498.1 | |
| ZNF558 | CDCA8 | RPL41P1 | |
| POLR3G | GAS5 | SMC4 | |
| DDIAS | CDCA3 | PHF21B | |
| MAGOHB | NUF2 | AC026740.1 | |
| MAD2L1 | CCNB1 | CDC20 | |
| AC068831.7 | AC015813.6 | GRIK2 | |
| KIAA1024 | ORC1 | AC112907.3 | |
| KIF18A | BLM | HPDL | |
| KIF23 | KIF14 | ATP5MC1P4 | |
| FBL | ARHGAP11A | VGF | |
| C1QL4 | MCM10 | TUBA3D | |
| SGO2 | EEF1A1P11 | PIF1 | |
| SNX18P7 | TRIP13 | TERT | |
| RPL41 | MKI67 | ADAM32 | |
| AURKA | EIF3C | RPL22L1 | |
| TUBA1B | UBE2C | PTPRD | |
| POLA2 | LCMT2 | SNHG3 | |
| CENPA | NEK2 | FAM72C | |
| ZFAS1 | EEF1A1P5 | SNHG26 | |
| HMMR | CSKMT | FAM72D | |
| APLN | TUBB | C2CD4C | |
| PARPBP | BIRC5 | GABRA3 | |
| MYBL2 | FANCB | AC132938.3 | |
