## Supplementary Method for "Pan-cancer analysis of pyrimidine metabolism reveals signaling pathways connections with chemoresistance role"

**Supplementary method. Script for calculating the activation scores for the signaling pathway or metabolic processes in cancer and meta-correlation analysis across cancer types.**

```
library(readr)
library(dplyr)
library(RcmdrMisc)
library(Hmisc)
library(gplots)
library(data.table)
library(openxlsx)

ctype <- read_delim("ACC_Infile.txt",delim = "\t",)
```

**### Step 1 - Fold difference (log2), mean and standard deviation estimations for the tumor profiles ###**

```
step1 <- subset(ctype,select= -c(Normal))

step1[2:ncol(step1)] <- step1[2:ncol(step1)]-ctype$Normal

stdev = step1 %>% summarise_if(is.numeric, sd)
meanf = step1 %>% summarise_if(is.numeric, mean)
```

**### Step 2 - Mean estimation for each gene-sets ###**

```
genesets <- read_csv2("genes.csv",col_names = FALSE)

step2 <- as.data.frame(matrix(nrow = nrow(genesets),ncol = ncol(step1)-1))

colnames(step2) <- colnames(step1[2:ncol(step1)])
rownames(step2) <- c(genesets$X1)

for (i in 1:nrow(step2)) {
  gen <- subset(step1,step1$Genes %in% genesets[i,2:ncol(genesets)])
  meanofset <- gen %>% summarise_if(is.numeric, mean)
  step2[i,] <- meanofset
}
```

**### Step 3 - Activation scores calculation for the gene-sets ###**

```
numberofgenes <- read_csv2("Genes_Count.csv",col_names = FALSE)
step3 <- step2

for (i in 1:nrow(step3)) {
  zscore <- (step3[i,]-meanf[1,])/(stdev[1,]/sqrt(numberofgenes[i,2])[[1]])
  step3[i,] <- zscore
}
```

**### Step 4 - Final activation score calculation for signaling pathways or metabolic processes ###**

```
forodd <- seq_len(nrow(step3)) %% 2
```

```
odds <- step3[forodd == 1, ]
evens <- step3[forodd == 0,]

forpym <- rep(0L,nrow(step3))
evens <- rbind(evens,forpym)

actscores <- odds - evens

rownames(actscores)[1:(nrow(actscores)-1)] <- gsub('.{3}$', '',
rownames(actscores)[1:(nrow(actscores)-1)])
```

**### Step 5 - Correlation matrix analysis ###**

```
var1 = as.data.frame(t(actscores),rownames="row_names",colnames= "col_names")
var2 = as.data.frame(t(actscores),rownames="row_names",colnames= "col_names")

cor.mat=cor(var1,var2)

# p and adjusted p values
Dataset <- var1
cor_mat=rcorr.adjust(Dataset, type="pearson")
pvalue = as.data.frame(cor_mat$R$P)
adjpvalue = as.data.frame(cor_mat$P)
correlation = as.data.frame(cor.mat)

# creating xlsx for results
finres <- as.data.frame(bind_rows(ACC_Correlation = correlation$PYRIMIDINE, ACC_Pvalue =
pvalue$PYRIMIDINE, ACC_AdjPvalue = adjpvalue$PYRIMIDINE))

rownames(finres) <- rownames(correlation)

finres$ACC_AdjPvalue <- as.numeric(finres$ACC_AdjPvalue)
finres$ACC_AdjPvalue[is.na(finres$ACC_AdjPvalue)] <- .0001

filtbyadj <- finres %>% filter(ACC_AdjPvalue < 0.05)

filtbycor <- filtbyadj %>% filter((ACC_Correlation > 0.3) | (ACC_Correlation < -0.3))

results <- list("Input_step1" = ctype,"Output_Step1" = step1, "Output_Step2"=step2,
            "Output_Step3"=step3, "Activation_Scores" = actscores, "Correlation" = correlation,
"PValue" = pvalue, "AdjPValue" = adjpvalue,
            "finalresults" = finres, "FilteredbyAdj" = filtbyadj, "FilterbyCor" = filtbycor)

write.xlsx(results, "ACC_Results.xlsx", row.names=TRUE)
```

**### Meta analysis using metacor.DSL ###**

```
library(dplyr)
library(metacor)

corr <- read.csv2("correlation.csv",row.names = 1)
corr[] <- sapply(corr, as.numeric)
```

```
adjP <- read.csv2("adjpval.csv",row.names = 1)
adjP[] <- sapply(adjP, as.numeric)

corr <- corr[!(row.names(corr) %in% "PYRIMIDINE"),]
adjP <- adjP[!(row.names(adjP) %in% "PYRIMIDINE"),]

unfilteredcorrelation <- corr

samplecount <- read.csv2("samplecounts.csv")

for (i in 1:nrow(unfilteredcorrelation)) {

  metcor <- metacor.DSL(r=as.numeric(unfilteredcorrelation[i,(1:32)]),
                      n= as.numeric(samplecount[,2]),
                   label= samplecount[,1])

  unfilteredcorrelation$"r.mean"[i] <- metcor$r.mean
  unfilteredcorrelation$"pval"[i] <- metcor$p
}

for (i in 1:nrow(corr)) {
  corr$posbefore[i] <- (length(which(corr[i,(1:32)]>0)))
  corr$negbefore[i] <- (length(which(corr[i,(1:32)]<0)))
}

for (i in 1:(nrow(adjP))) {
  for (j in 1:(ncol(adjP))) {
    if((adjP[i,j]>= 0.05) || between(corr[i,j],-.3,.3)){
      corr[i,j] <- NA
    }

  }

}

for (i in 1:nrow(corr)) {
  corr$posafter[i] <- (length(which(corr[i,(1:32)]>0)))
  corr$negafter[i] <- (length(which(corr[i,(1:32)]<0)))
}



for (i in 1:nrow(corr)) {
  corr$posrat[i] <- (length(which(corr[i,(1:32)]>0))/32)
  corr$negrat[i] <- (length(which(corr[i,(1:32)]<0))/32)
}

for (i in 1:nrow(corr)) {
  if(length(which(is.na(corr[i,]))) <31 ){
    metcorfiltered <- metacor.DSL(r = as.numeric(corr[i,c(which(!is.na(corr[i,(1:32)])))]),
                          n = as.numeric(samplecount[c(which(!is.na(corr[i,(1:32)]))),2]),
                          label = samplecount[c(which(!is.na(corr[i,(1:32)]))),1])
```

```
    corr$"r.mean"[i] <- metcorfiltered$r.mean
    corr$"pval"[i] <- metcorfiltered$p

  }

else {
    corr$"r.mean"[i] <- 0
    corr$"pval"[i] <- 1
  }
}

write.table(corr,"correlation_filteredwithscoreSignaling.Pym.txt",sep = "\t")
write.table(unfilteredcorrelation,"correlation_unfilteredwithscoreSignaling.Pym.txt",sep = "\t")
```
